# Molecular Mimicry in Inflammatory Bowel Disease: Multi-layered Functional and Sequence-level Analysis of Gut Microbial Proteins Mimicking the Human Proteome

**DOI:** 10.64898/2026.03.20.713231

**Authors:** Ananya Anurag Anand, Payal Mishra, Vejendla Sharmil Srivathsa, Vidushi Yadav, Sintu Kumar Samanta

## Abstract

Inflammatory bowel disease (IBD), encompassing Crohn’s disease (CD) and ulcerative colitis (UC), is a chronic inflammatory disorder whose pathogenesis involves intricate host-microbiome interactions. Molecular mimicry (the structural or functional resemblance between microbial and host proteins) represents a plausible mechanism by which gut microbiota may trigger or perpetuate autoimmune responses. Here we present a comprehensive, multi-layered molecular mimicry in silico pipeline (MMIP) analysis of 39 baseline shotgun metagenomic samples from Human Microbiome Project 2 (HMP2/IBDMDB). Using DIAMOND-based homology search against the SwissProt followed by UniProt Retrieve/ID Mapping (URIM), we characterized microbial protein functional space through three complementary frameworks: (i) normalized GO term frequency comparison across diagnostic groups, (ii) protein family (PFAM) domain enrichment analysis, and (iii) sequence-level mimicry analysis identifying microbial proteins with direct homology to human proteins. Taxonomic profiling using MetaPhlAn3 pre-computed abundance profiles provided further biological context. CD microbiomes exhibited greater enrichment of immune-relevant biological processes and a higher per-sample sequence-level mimicry rate than UC. CD and UC also showed distinct pathobiont profiles, with CD enriched for oral-origin taxa including *Haemophilus parainfluenzae* and UC enriched for *Fusobacterium nucleatum* and *Prevotella* species. Notably, healthy gut microbiome maintains coordinated mimicry of host neuronal, signal-recognition-particle-associated, and antimicrobial peptide machinery, a repertoire dismantled in IBD and replaced by disease-subtype-specific signatures, alongside a candidate mimicry link between CD-enriched bacteria and NOD2. These results represent the first metagenome-wide characterization of molecular mimicry across IBD subtypes using shotgun metagenomic data, offering new mechanistic insight into how microbial dysbiosis may contribute to immune dysregulation in IBD.

## 1. Introduction

Inflammatory bowel disease (IBD) affects more than 10 million individuals worldwide **[Ng et al., 2017]** and is defined by two major subtypes, Crohn’s disease (CD) and ulcerative colitis (UC), which share a common framework of immune dysregulation directed against the gut microbiota but differ markedly in anatomical distribution, depth of mucosal involvement, and clinical course. CD can affect any segment of the gastrointestinal tract with transmural, discontinuous inflammation, while UC is confined to the colonic mucosa. Despite decades of research, the precise mechanisms linking microbial dysbiosis to the initiation and perpetuation of intestinal inflammation remain incompletely understood.

The gut microbiome is increasingly recognized as a central participant in IBD pathogenesis. The landmark Human Microbiome Project 2 (HMP2) multi-omics study of IBD demonstrated profound shifts in microbial community composition and functional potential in IBD patients, including depletion of short-chain fatty acid producers such as *Faecalibacterium prausnitzii* and expansion of pathobionts including *Haemophilus parainfluenzae* and *Escherichia coli* **[Lloyd-Price et al., 2019]**. However, compositional alterations alone do not fully explain the immunological consequences of microbial dysbiosis, and functional characterization of how microbial proteins interact with host biology remains an active area of investigation.

Molecular mimicry — the phenomenon whereby microbial proteins share structural or functional similarity with host proteins — represents a mechanistically compelling route through which gut microbiota may contribute to immune dysregulation. First described in the context of autoimmune diseases triggered by viral or bacterial infections **[Cusick et al., 2012]**, molecular mimicry operates through two distinct mechanisms: sequence-level mimicry, where microbial proteins share linear amino acid similarity with host proteins potentially enabling cross-reactive immune responses, and functional mimicry, where microbial proteins share pathway membership or functional annotations with host counterparts, reflecting evolutionary convergence **[Suliman, 2024]**. In IBD, where the mucosal immune system is in perpetual contact with a dense and diverse microbial community at a compromised epithelial barrier, both forms of mimicry could have profound immunological consequences.

Despite its conceptual importance, systematic computational characterization of molecular mimicry across the gut metagenome in IBD has been limited by methodological constraints. Prior efforts have often relied on 16S rRNA amplicon sequencing, which lacks resolution to identify protein-coding sequences **[Joossens et al., 2011]**. Shotgun metagenomics enables direct recovery and annotation of microbial protein-coding sequences, making it ideally suited for molecular mimicry analysis at metagenome scale **[Quince et al., 2017]**.

Here we apply the Molecular Mimicry In Silico Pipeline (MMIP) to 39 baseline shotgun metagenomic samples from the HMP2 IBDMDB cohort, representing the first application of this framework to whole-metagenome sequencing data from a well-characterized IBD cohort. Our multi-layered analytical framework integrates DIAMOND-based homology search **[Buchfink et al., 2021]**, UniProt functional annotation **[UniProt Consortium, 2025]**, normalized GO term frequency analysis, PFAM domain enrichment **[Mistry et al., 2021]**, and sequence-level mimicry detection, validated against MetaPhlAn3 taxonomic profiles **[Beghini et al., 2021]**. The results reveal systematic differences in the microbial functional mimicry landscape between CD, UC, and healthy individuals, with implications for understanding the mechanisms by which microbial dysbiosis may contribute to immune dysregulation in IBD.

## 2. Materials and Methods

### 2.1 Dataset and Sample Selection

Shotgun metagenomic sequencing data were obtained from the Human Microbiome Project 2 Inflammatory Bowel Disease Multi-omics Database (HMP2 IBDMDB; https://ibdmdb.org) **[Lloyd-Price et al., 2019]**. Pre-computed gene calls generated by the HMP2 Consortium using the Prodigal gene prediction algorithm **[Hyatt et al., 2010]** were downloaded from the IBDMDB data products page. To minimize confounding from disease activity fluctuations and treatment effects, the first available (baseline) sample per participant was selected, yielding a final cohort of 39 samples: 12 non-IBD (healthy), 17 Crohn’s disease (CD), and 10 ulcerative colitis (UC).

### 2.2 Protein Sequence Preparation

Nucleotide gene call sequences, provided as Prodigal-predicted open reading frames (ORFs) in FASTA format, were translated to protein sequences using BioPython (version 1.81) **[Cock et al., 2009]**. Strand directionality encoded in Prodigal FASTA headers (field 4 of semicolon-delimited header: +1 forward, −1 reverse) was used for reverse complementation where appropriate prior to translation. Predicted protein sequences shorter than 10 amino acids were discarded. This yielded 31,071, 31,135, and 24,775 predicted protein sequences for the nonIBD, CD, and UC groups respectively.

### 2.3 Homology Search Against SwissProt Using DIAMOND

A DIAMOND (version 2.1.9) **[Buchfink et al., 2021]** database was constructed from the UniProt Swiss-Prot manually reviewed protein database (574,627 sequences) **[UniProt Consortium, 2025]**. For each of the 39 samples, predicted protein sequences were queried against this database using DIAMOND blastp with the following parameters: sensitive mode (--sensitive), maximum e-value of 1×10⁻⁵ (--evalue 1e-5), one maximum target sequence per query (--max-target-seqs 1), and four CPU threads (--threads 4). The single top-scoring hit per query was retained. Output was generated in tabular format (outfmt 6) including query sequence ID, subject sequence ID, percentage identity, alignment length, number of mismatches, gap openings, query and subject start/end coordinates, e-value, bitscore, and subject title (stitle).

### 2.4 UniProt Functional Annotation via URIM

UniProt accession identifiers were extracted from the sseqid field of DIAMOND output files by parsing the pipe-delimited format (sp|ACCESSION|ENTRYNAME). Accession IDs from all samples within each diagnostic group were pooled and deduplicated, yielding three group-level ID lists. These were submitted to the UniProt Retrieve/ID Mapping (URIM) web service **[Zaru & Orchard, 2023]**, mapping from UniProtKB AC/ID to UniProtKB, with retrieval of functional metadata including Gene Ontology (GO) annotations **[Ashburner et al., 2000]**, GO term identifiers, PFAM domain assignments **[Mistry et al., 2021]**, organism classification, Ensembl gene cross-references, protein sequences, and EC numbers. This constituted URIM Output 1 (all-organism hits), used for functional mimicry analyses.

For sequence-level mimicry, DIAMOND output records where the subject title contained ‘OS=Homo sapiens’ were extracted separately per group. Corresponding UniProt accession IDs were submitted to URIM by the same procedure, yielding URIM Output 2 (human-only hits).

### 2.5 Normalized GO Term Frequency Analysis of URIM Output 1

Gene Ontology identifier strings were extracted from the ‘Gene Ontology IDs’ column of URIM Output 1 TSV files. For each group, GO term occurrence counts were computed across all UniProt entries. Raw counts were normalized by total predicted ORFs per group and multiplied by 10⁵ to yield normalized GO term frequencies (per 100,000 ORFs), accounting for differences in total predicted protein sequences between groups (nonIBD: 31,071; CD: 31,135; UC: 24,775).

Inter-group comparisons were performed by computing delta normalized frequency (Δfreq = freq₂ − freq₁) and log₂ fold change [log₂FC = log₂((freq₂ + 0.01)/(freq₁ + 0.01))] for three comparisons: CD vs nonIBD, UC vs nonIBD, and CD vs UC. A pseudocount of 0.01 was added to prevent division by zero. GO term names and namespace assignments were added by cross-referencing identifiers against the Gene Ontology Basic (go-basic.obo) ontology file **[Gene Ontology Consortium, 2023]**. Human-specific term sizes were retrieved from the Gene Ontology Annotation (GOA) human annotation file (goa_human.gaf, EBI FTP) **[Huntley et al., 2015]**. Differential analysis applied a three-filter approach: (i) retention of biological process (GO:BP) terms only; (ii) exclusion of overly generic metabolic or cellular process terms, as follows: ‘metabolic’, ‘biosynthetic’, ‘cellular process’, ‘primary’, ‘compound’, ‘nucleic acid’, ‘macromolecule’, ‘organization’, ‘component’, ‘system process’, ‘homeostasis’, ‘development’, ‘binding’, ‘membrane’, ‘cytoplasm’; (iii) retention of terms with human term size below 500 genes. A two-filter analysis was also performed using only the first two filters from the three-filter approach. Additionally, a no-filter approach was also applied.

### 2.6 g:Profiler Enrichment Analyses

Two complementary g:Profiler-based enrichment analyses were performed to statistically evaluate the significance of GO term representation in each diagnostic group against the human genome background, using the gprofiler-official Python package (version 1.0.0) **[Raudvere et al., 2019]**.

#### 2.6.1 Direct GO ID Enrichment (Analysis A)

In the first approach, GO identifiers extracted from URIM Output 1 were submitted directly as query input to g:Profiler using the profile() function with organism=’hsapiens’, sources=[’GO:BP’, ‘GO:MF’, ‘GO:CC’], and a significance threshold of p < 0.05. This approach tests whether the GO terms represented in the microbial protein hits are statistically overrepresented relative to what would be expected by chance in the human genome annotation. Results were obtained for nonIBD (189 enriched terms), CD (191 enriched terms), and UC (187 enriched terms). Pairwise differential comparisons between groups were performed using the delta -log10(p-value) approach: enriched terms from two groups were merged on native GO ID, and Δ(-log₂FC p-value) was computed as the difference in -log₁₀(p-value) between groups. Three-filter analysis was applied as described in Section 2.5.

#### 2.6.2 Ensembl Gene ID-based Enrichment (Analysis B)

In the second approach, GO identifiers from URIM Output 1 were first converted to Ensembl human gene identifiers using the g:Profiler convert() function with target_namespace=’GO’, mapping each GO term to the set of human genes annotated with that term. This yielded 22,941, 22,938, and 22,939 unique Ensembl gene IDs for nonIBD, CD, and UC respectively. These gene ID lists were submitted to g:Profiler profile() with organism=’hsapiens’, sources=[’GO:BP’, ‘GO:MF’, ‘GO:CC’], and p < 0.05, yielding 184, 188, and 191 enriched terms respectively. Differential gene representation between groups was assessed by computing normalized gene frequencies (count per 100,000 ORFs) and log₂ fold changes between diagnostic groups. Genes with |log₂FC| > 1 were designated as differentially represented and submitted separately to g:Profiler for targeted enrichment analysis.

### 2.7 PFAM Domain Enrichment Analysis

PFAM domain assignments were extracted from the ‘PFAM’ column of URIM Output 1 TSV files **[Mistry et al., 2021].** Raw PFAM domain counts were normalized per 100,000 ORFs for each group. Differential enrichment was assessed by computing delta normalized frequencies and log₂ fold changes between groups. Domains enriched in IBD groups relative to nonIBD were functionally categorized to assess the prevalence of eukaryote-like architectural features.

### 2.8 Sequence-level Mimicry Analysis

DIAMOND output records matching Homo sapiens proteins (stitle containing ‘OS=Homo sapiens’) were isolated per group. Total human-matching hit counts were normalized per 100,000 ORFs to enable cross-group comparison. UniProt accession identifiers from URIM Output 2 were used to characterize the human proteins most frequently targeted by microbial sequence mimics. Human sequence mimicry rates were normalized by mean predicted ORFs per sample within each diagnostic group rather than group-level ORF totals, to account for differences in sample number between groups (nonIBD: n=12, CD: n=17, UC: n=10) and enable per-sample-level comparison. The frequency of each unique human UniProt accession across all samples within a group was computed and normalized per 100,000 ORFs to identify the human proteins most recurrently targeted by microbial sequence mimics.

### 2.9 Taxonomic Abundance Analysis

Taxonomic abundance profiles were obtained from the pre-computed merged MetaPhlAn3 table from IBDMDB and filtered to the 39 study samples **[Beghini et al., 2021]**. Per-sample relative abundance profiles were separated by diagnostic group and filtered to species-level features, defined as rows containing ‘s ‘ in the lineage string, excluding entries labelled UNKNOWN. Mean relative abundances were computed per species per diagnostic group across all samples within that group, with missing values (species absent from a sample) treated as zero prior to averaging. Log₂ fold changes between diagnostic groups were calculated for three pairwise comparisons — CD vs. nonIBD, UC vs. nonIBD, and CD vs. UC — using the formula log₂FC = log₂((mean_abundance₂ + 0.01) / (mean_abundance₁ + 0.01)), where a pseudocount of 0.01 was added to both numerator and denominator prior to log₂ transformation to prevent division by zero for species absent in one group, consistent with the pseudocount approach employed in the GO term frequency analyses (Section 2.5). Species with |log₂FC| > 1 relative to nonIBD in either the CD vs. nonIBD or UC vs. nonIBD comparison were designated as differentially abundant and reported as IBD-enriched or nonIBD-enriched accordingly. All analyses were performed in Python using pandas (version 2.x) **[McKinney, 2010]** and numpy **[Harris et al., 2020]**.

### 2.10 Resolve-to-Taxon Sequence Mimicry Analysis (Bottom-Up Validation Analysis)

While group-level pooled identifier strategies provide a wide-lens summary of overall functional shifts, they obscure the precise microbial sequences and specific operational taxonomic units (OTUs) driving individual mimicry signals. To address this limitation and establish clear, sequence-level causality, we deployed a parallel, single-ORF resolution discovery approach. This pipeline traces sequence-level homology profiles at the single-molecule scale directly back to both their specific sample of origin and their exact bacterial donor species.

All predicted protein open reading frames (ORFs) from the 39 study samples were serialized in parquet format alongside contextual metadata (Sample_ID, Disease_Group). Sequences were converted to FASTA format and partitioned into chunks of 200,000 for high-throughput distributed computation. A custom reference framework was constructed using a local DIAMOND (version 2.1.11) blastp database built exclusively from the UniProt Swiss-Prot reviewed human proteome (downloaded via UniProt REST API, organism_id:9606, reviewed:true). Individual sequence chunks were queried under stringent alignment bounds (--evalue 1e-5, --max-target-seqs 1, --outfmt 6, --threads 4). Alignment pairs were discarded if they exhibited a percentage identity < 35% or an alignment length < 8 amino acids.

Because the query headers retained implicit indexing of individual sample origins, we computed sample-wise human target abundance vectors. Human protein targets were then assigned to categorical enrichment classes using a majority-voting scheme determined by the dominant diagnostic group contributing to that sequence cluster: *CD_Enriched*, *UC_Enriched*, *Healthy_Enriched*, *Common_Disease*, or *Low_Evidence* (n < 3 global ORFs). Group-wise differential abundance was evaluated via Log₂ fold changes using a pseudocount of 0.5.

To rigorously confirm the microbial provenance of these hits and strip away potential host-sequencing contamination, a secondary sequence-level verification pass was performed. Every mimicking ORF was back-queried against the full, multi-kingdom UniProt Swiss-Prot database. An aggressive cross-kingdom exclusion filter was implemented: any ORF whose ultimate top-scoring Swiss-Prot hit aligned to a eukaryotic reference (including *Homo sapiens*, *Mus musculus*, *Arabidopsis thaliana*, *Rattus norvegicus*, *Bos taurus*, *Oryza sativa*, *Drosophila melanogaster*, *Saccharomyces cerevisiae*, *Caenorhabditis elegans*, *Danio rerio*, *Xenopus spp.*, *Sus scrofa*, *Gallus gallus*, *Dictyostelium discoideum*) was removed. Only sequences showing bacterial alignment roots (n = 45,088 ORFs) were taken for downstream taxonomic mapping. Final outputs recorded the exact bacterial organism, protein name, gene identifier, and NCBI TaxID to enable high-resolution organismal attribution.

### 2.11 Statistical Analysis and Visualization

All normalization and differential analyses were performed in Python 3.13 with pandas **[McKinney, 2010]** and NumPy **[Harris et al., 2020]**. GO term name mapping used local go-basic.obo and goa_human.gaf files for reproducibility. Figures were generated using Python **[Hunter, 2007]**.

**Figure 1** details the complete Molecular Mimicry In Silico Pipeline (MMIP).

**Figure 1.**
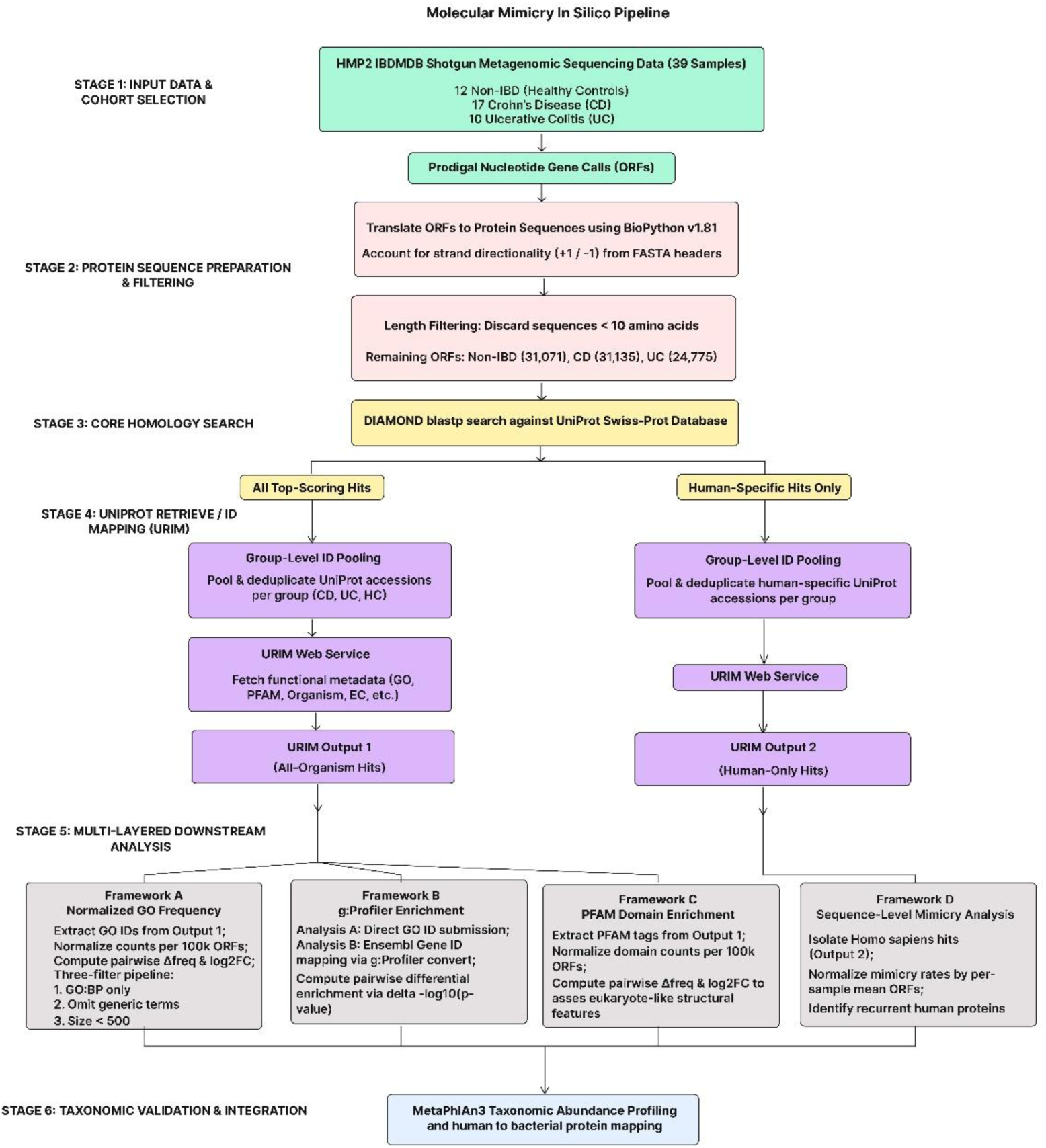
Molecular Mimicry In Silico Pipeline (MMIP) workflow for IBD metagenomics

## 3. Results

### 3.1 Dataset Overview and ORF Statistics

Thirty-nine baseline shotgun metagenomic samples from the HMP2 IBDMDB cohort were analyzed: 12 healthy non-IBD individuals, 17 Crohn’s disease patients, and 10 ulcerative colitis patients. Translation of pre-computed Prodigal gene calls yielded a total of 1,789,492 predicted protein sequences across all samples. The mean number of predicted ORFs per sample differed substantially between groups: nonIBD samples contained a mean of 58,388 ORFs per sample (range: 29,762–92,425), compared to 40,134 ORFs per sample in CD (range: 168–78,522) and 40,657 ORFs per sample in UC (range: 65–83,687) (Figure 2).

**Figure 2.**
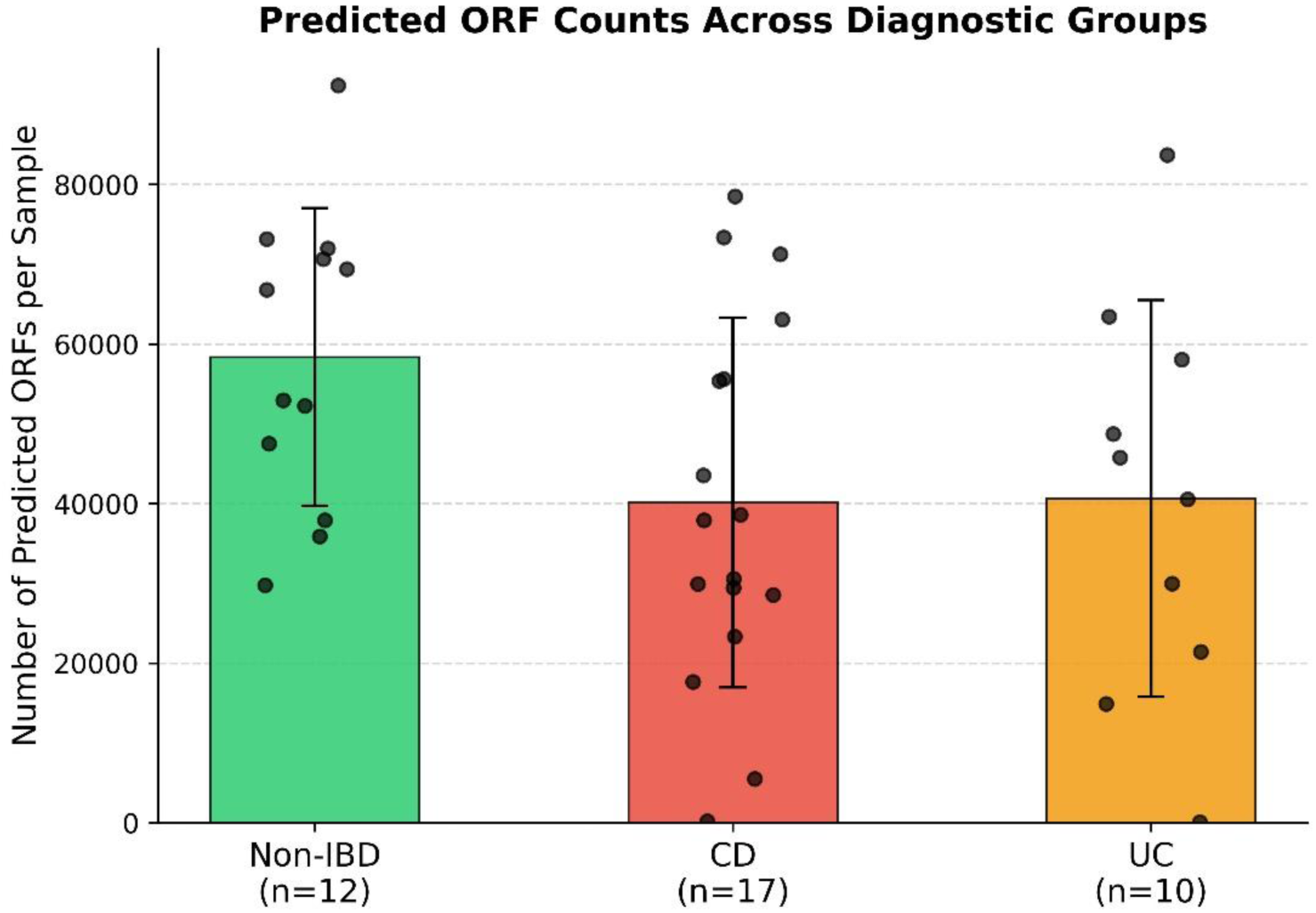
— ORF counts per diagnostic group with individual sample values overlaid.

This notable difference in per-sample ORF counts, with nonIBD samples containing more predicted proteins on average than IBD samples, reflects either higher microbial biomass, greater sequencing depth, or both in healthy individuals, and motivated the per-sample normalization approach used throughout all quantitative analyses **[Gevers et al., 2014].**

The CD and UC cohort had 1 sample each with extremely low ORF counts (might be due to lesser sequencing depth). These were therefore removed for a better comparative analysis of the mean, standard deviation and other statistics, which were then re-calculated **(Table 1)**. The results showed that the mean was slightly affected after the removal of under-represented samples. However, the nonIBD samples still showed more predicted ORFs than UC and CD.

**Table 1.** Predicted ORF counts per sample across diagnostic groups, before and after removal of under-represented samples.

| Dataset | Group | Count | mean | std | min | max |
| --- | --- | --- | --- | --- | --- | --- |
| Original (all samples) | nonIBD | 12 | 58387.58 | 18673.72 | 29762 | 92425 |
| Original (all samples) | CD | 17 | 40133.65 | 23175.36 | 168 | 78522 |
| Original (all samples) | UC | 10 | 40656.9 | 24857.43 | 65 | 83687 |
| Filtered (underrepresented removed) | nonIBD | 12 | 58387.58 | 18673.72 | 29762 | 92425 |
| Filtered (underrepresented removed) | CD | 16 | 42631.5 | 21442.13 | 5492 | 78522 |
| Filtered (underrepresented removed) | UC | 9 | 45167.11 | 21593.54 | 14878 | 83687 |

### 3.2 DIAMOND Homology Search Against SwissProt

DIAMOND blastp homology search of all predicted protein sequences against the SwissProt database (574,627 reviewed sequences, e-value ≤ 1×10⁻⁵) yielded significant hits for approximately 29–30% of predicted ORFs per sample across all three diagnostic groups (Figure 3A). The proportion of ORFs with significant SwissProt hits was comparable between groups (nonIBD: mean 29.5%, SD 1.8%; CD: mean 30.8%, SD 4.3%; UC: mean 27.9%, SD 9.4%), indicating broadly similar rates of homology to reviewed proteins across all conditions. The higher variance in CD and UC reflects greater inter-sample heterogeneity in these disease groups. After removing the two under-represented samples, recalculated statistics **(Table 2)** showed that UC variance dropped substantially (SD: 9.43% → 3.79%), confirming that the originally elevated UC standard deviation was driven primarily by the single low-depth outlier sample (MSM9VZLZ), rather than reflecting genuine inter-sample heterogeneity within the UC cohort. In contrast, CD variance remained largely unchanged (SD: 4.34% → 4.37%) following removal of its outlier sample, indicating that CD’s higher variability relative to nonIBD reflects true biological heterogeneity rather than a single low-quality sample. After filtering, CD and UC showed comparable variance (SD 4.37% and 3.79%, respectively), both modestly higher than the tightly distributed nonIBD cohort (SD 1.78%).

**Figure 3A.**
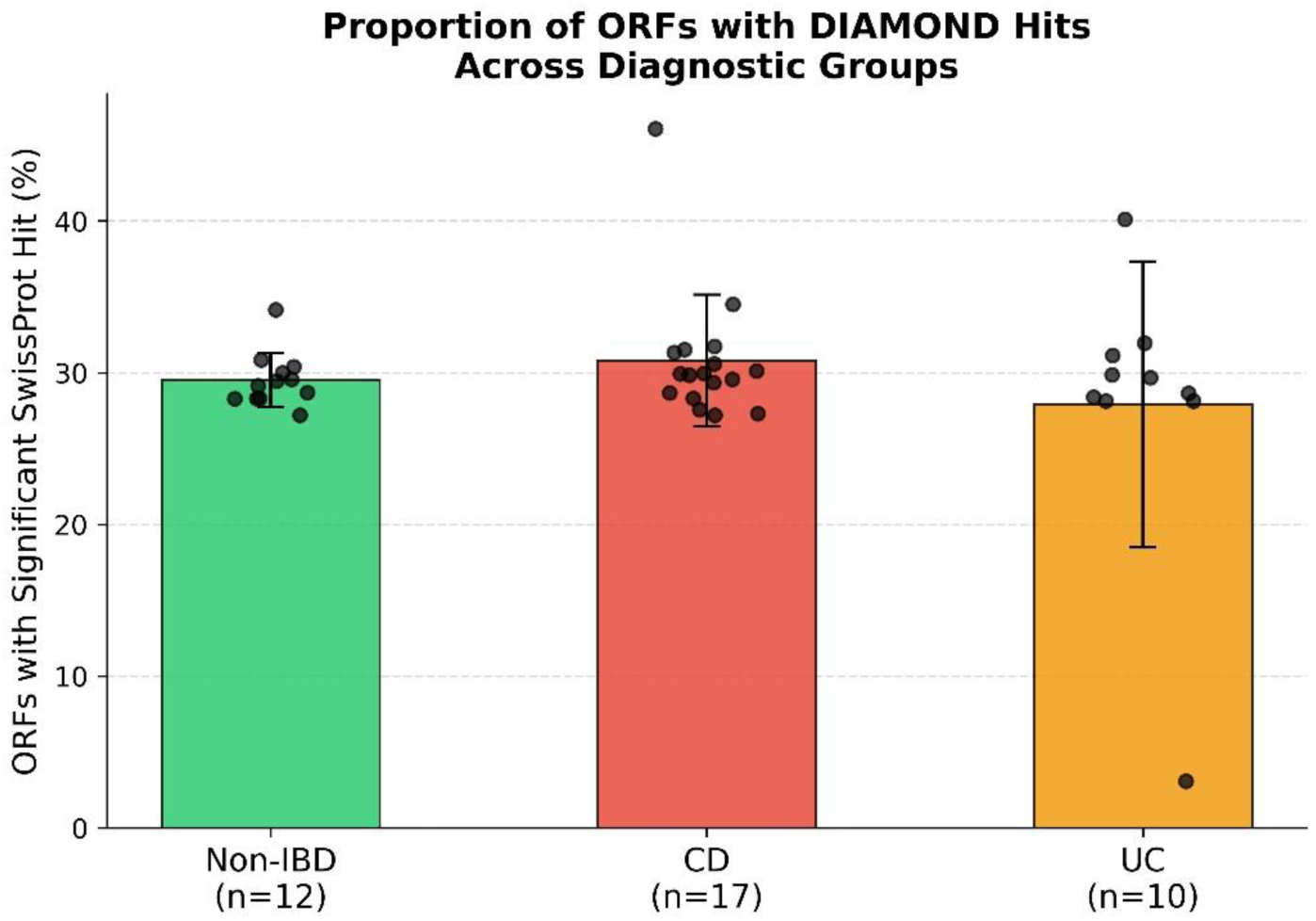
— Proportion of ORFs with significant SwissProt DIAMOND hits per group.

**Table 2.** Percentage of ORFs with significant SwissProt hits: original vs. filtered cohorts.

| Group | Dataset | count | mean | std | min | max |
| --- | --- | --- | --- | --- | --- | --- |
| nonIBD | Original (all samples) | 12 | 29.53 | 1.78 | 27.19 | 34.15 |
| CD | Original (all samples) | 17 | 30.8 | 4.34 | 27.2 | 46.07 |
| UC | Original (all samples) | 10 | 27.92 | 9.43 | 3.08 | 40.12 |
| nonIBD | Filtered (underrepresented removed) | 12 | 29.53 | 1.78 | 27.19 | 34.15 |
| CD | Filtered (underrepresented removed) | 16 | 30.56 | 4.37 | 27.2 | 46.07 |
| UC | Filtered (underrepresented removed) | 9 | 30.68 | 3.79 | 28.14 | 40.12 |

The distribution of percentage identity between microbial query sequences and their top SwissProt hits was nearly identical across all three groups, with median values of approximately 40–41% identity (nonIBD: mean 47.3%, CD: mean 47.9%, UC: mean 48.1%) and comparable interquartile ranges (**Figure 3B**, **Table 3**). This consistency indicates that the degree of sequence divergence between gut microbial proteins and their SwissProt homologs does not differ systematically between diagnostic groups, and that differences observed in downstream analyses reflect more of the quantitative rather than qualitative changes in the microbial functional repertoire.

**Table 3.** Distribution of DIAMOND hit percentage identity across diagnostic groups.

| Group | Mean % Identity to SwissProt hits | std | min | 25% | 50% | 75% | max |
| --- | --- | --- | --- | --- | --- | --- | --- |
| **CD** | 47.87 | 21.54 | 17.6 | 31.9 | 40.6 | 56.8 | 100 |
| **UC** | 48.1 | 21.15 | 17.8 | 32.5 | 41.3 | 56.9 | 100 |
| **Non-IBD** | 47.29 | 20.33 | 17.6 | 32.2 | 40.9 | 56 | 100 |

**Figure 3B.**
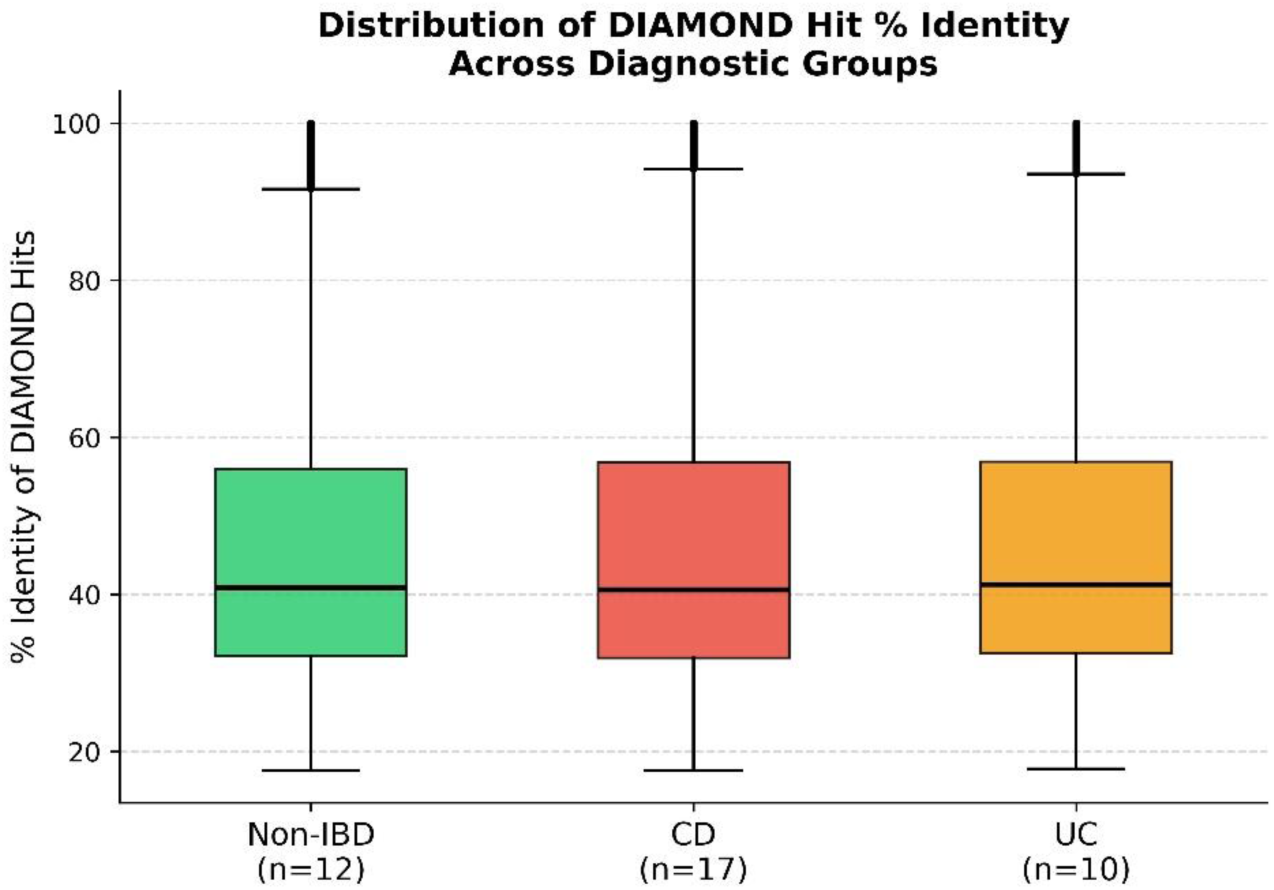
— Distribution of % identity of DIAMOND hits across groups.

### 3.3 Normalized GO Term Frequency Analysis Reveals Differential Functional Mimicry

A total of 9,372 unique GO terms were detected across the combined dataset after extraction from URIM. After normalization by per-sample ORF counts (per 100,000 ORFs), pairwise differential analysis was performed between all three diagnostic group pairs. The three-filter pipeline (biological process terms only; exclusion of generic metabolic and cellular process terms; term size < 500 human genes) yielded 2,403 specific biological process terms for comparative analysis. Several GO terms were found exclusive to a single cohort-*577 IBD-exclusive,* 327 CD-exclusive and 239 UC-exclusive BP terms (**Supplementary File 1**). Three-filter analysis retained GO:Biological Process terms only, excluding generic metabolic and cellular process terms, with human term size < 500 genes. Enrichment direction indicates the cohort with higher normalized GO term frequency in each pairwise comparison **(Table 4)**.

**Table 4.** Pairwise differential GO term counts following three-filter analysis.

| Comparison | Direction | Count |
| --- | --- | --- |
| CD vs nonIBD | Enriched in CD | 292 |
| CD vs nonIBD | Enriched in nonIBD | 571 |
| UC vs nonIBD | Enriched in UC | 315 |
| UC vs nonIBD | Enriched in nonIBD | 527 |
| CD vs UC | Enriched in CD | 402 |
| CD vs UC | Enriched in UC | 341 |

#### 3.3.1 GO Terms Enriched in CD vs Non-IBD

The Crohn’s disease-associated microbiome showed enrichment of multiple specific biological process terms relative to healthy controls, with top terms including adaptive immune response (log₂FC = 10.33), regulation of leukocyte cell-cell adhesion (log₂FC = 9.33), regulation of leukocyte tethering or rolling (log₂FC = 9.33), bone remodeling (log₂FC = 9.33), cellular response to misfolded protein (log₂FC = 9.33), and mitochondrial electron transport (log₂FC = 9.33) (Figure 4A). These terms span immune regulation, cellular stress responses, and metabolic processes, collectively suggesting that CD-associated microbial communities harbor a disproportionate functional representation of proteins related to processes relevant to intestinal inflammation and systemic disease manifestations.

**Figure 4A.**
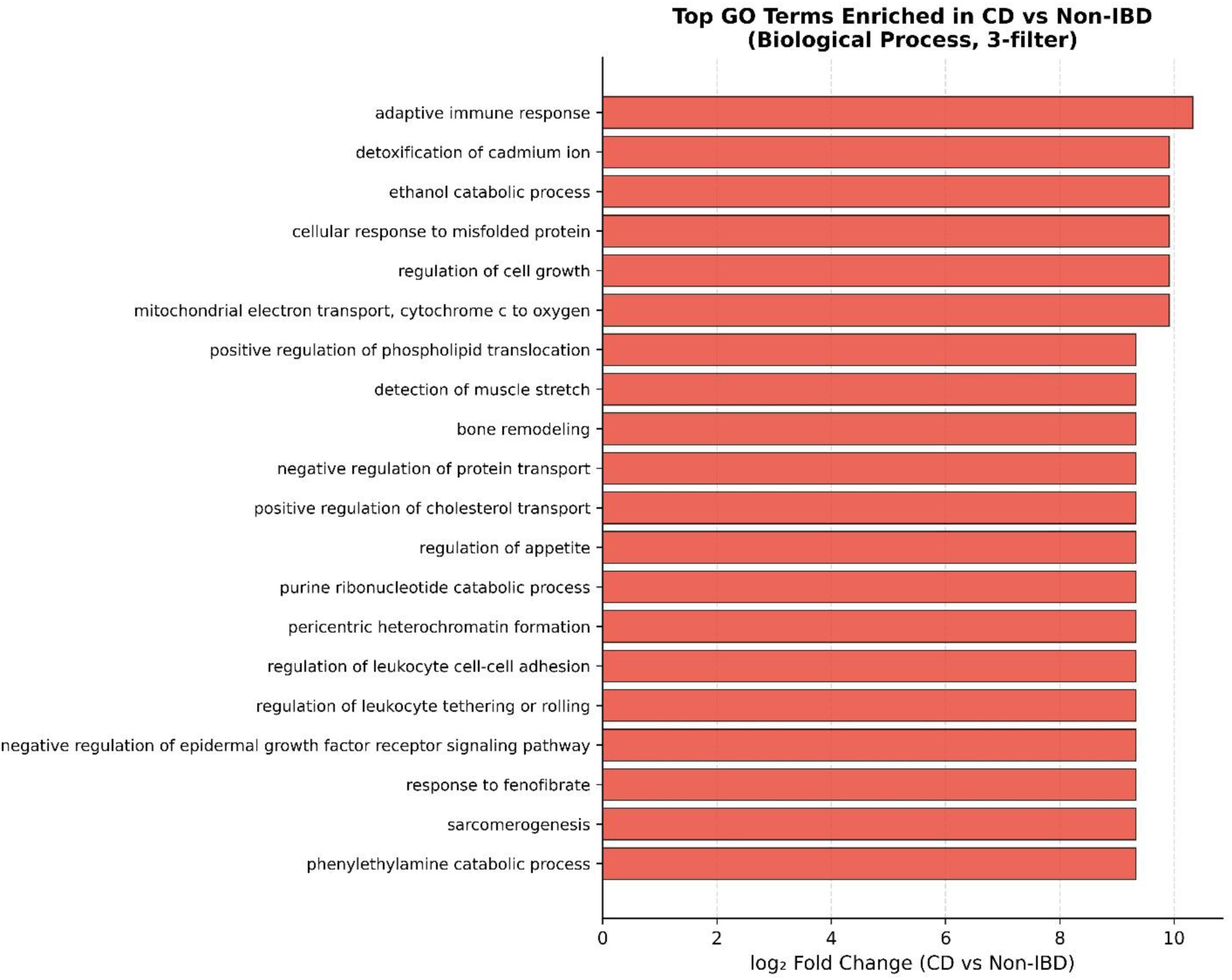
— Top 20 GO terms enriched in CD vs non-IBD (3-filter, biological process).

#### 3.3.2 GO Terms Enriched in Non-IBD vs CD (Protective Functional Signatures)

Conversely, healthy non-IBD microbiomes were enriched in distinct biological process terms relative to CD, with the most strongly depleted-in-CD terms including RTX prokaryotic toxin activity (log₂FC = −10.65), Class-III pyridoxal-phosphate-dependent aminotransferase activity (log₂FC = −10.33), and multiple terms related to basic metabolic enzyme activities (Figure 4B). These represent functional categories enriched in the healthy gut microbiome that are relatively depleted in CD, potentially reflecting differences in microbial community composition between healthy and CD-associated gut environments..

**Figure 4B.**
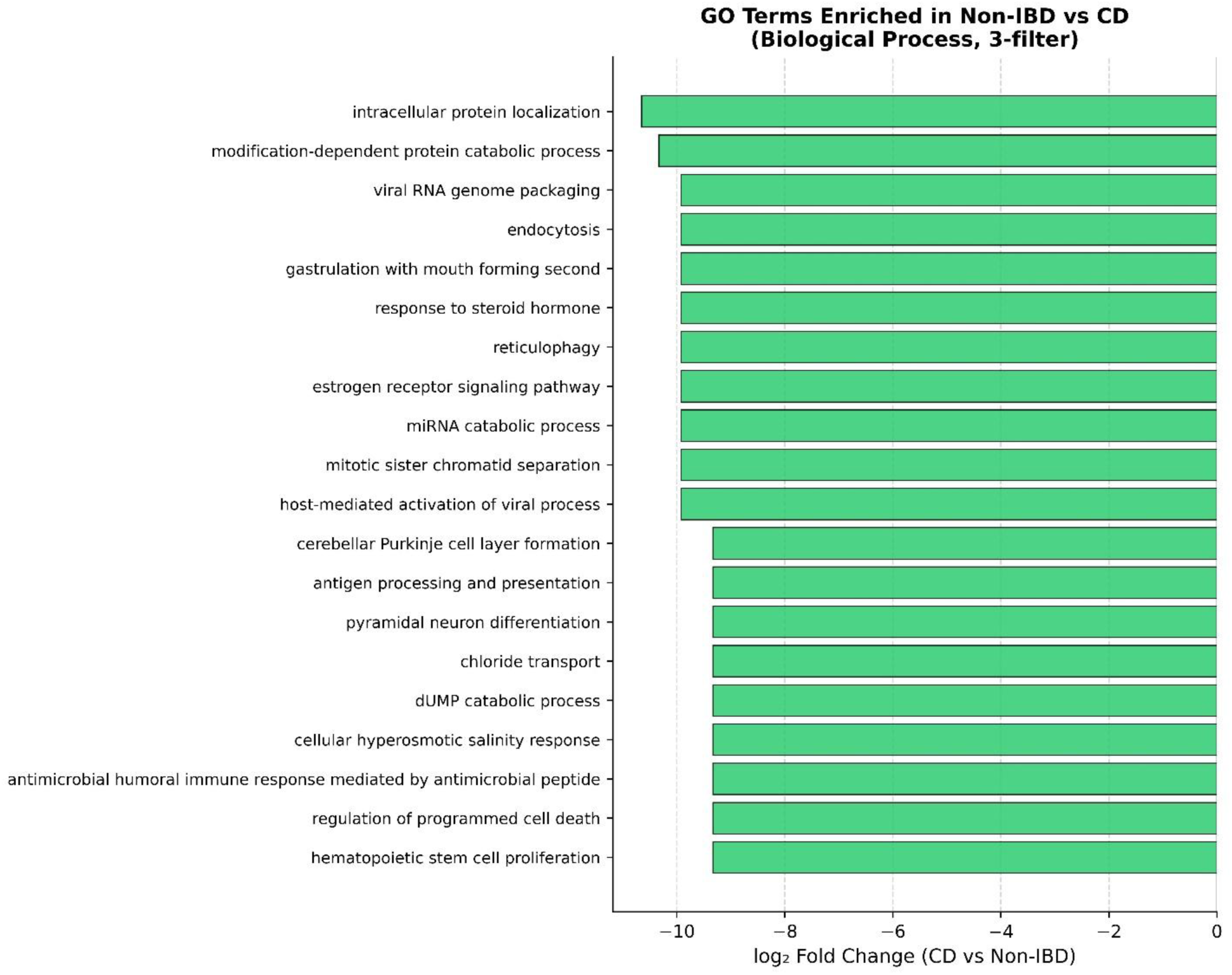
— Top 20 GO terms enriched in non-IBD vs CD.

#### 3.3.3 GO Terms in UC vs Non-IBD: Attenuated Differential Signal

Top UC-enriched terms included regulation of cell growth (log₂FC = 10.24), lipopolysaccharide-mediated signaling pathway (log₂FC = 9.66), cytokine-mediated signaling pathway (log₂FC = 9.66), and T cell receptor signaling pathway (log₂FC = 9.66) (Figure 4C). Healthy-enriched terms relative to UC are shown in Figure 4D. Both CD and UC microbiomes showed comparable numbers of differentially represented GO terms relative to healthy controls after 3-filter analysis (CD: 863 total; UC: 842 total), with healthy microbiomes showing greater functional enrichment relative to both disease groups (571 terms enriched in nonIBD vs CD; 527 terms enriched in nonIBD vs UC) than vice versa (292 enriched in CD vs nonIBD; 315 enriched in UC vs nonIBD).

**Figure 4C.**
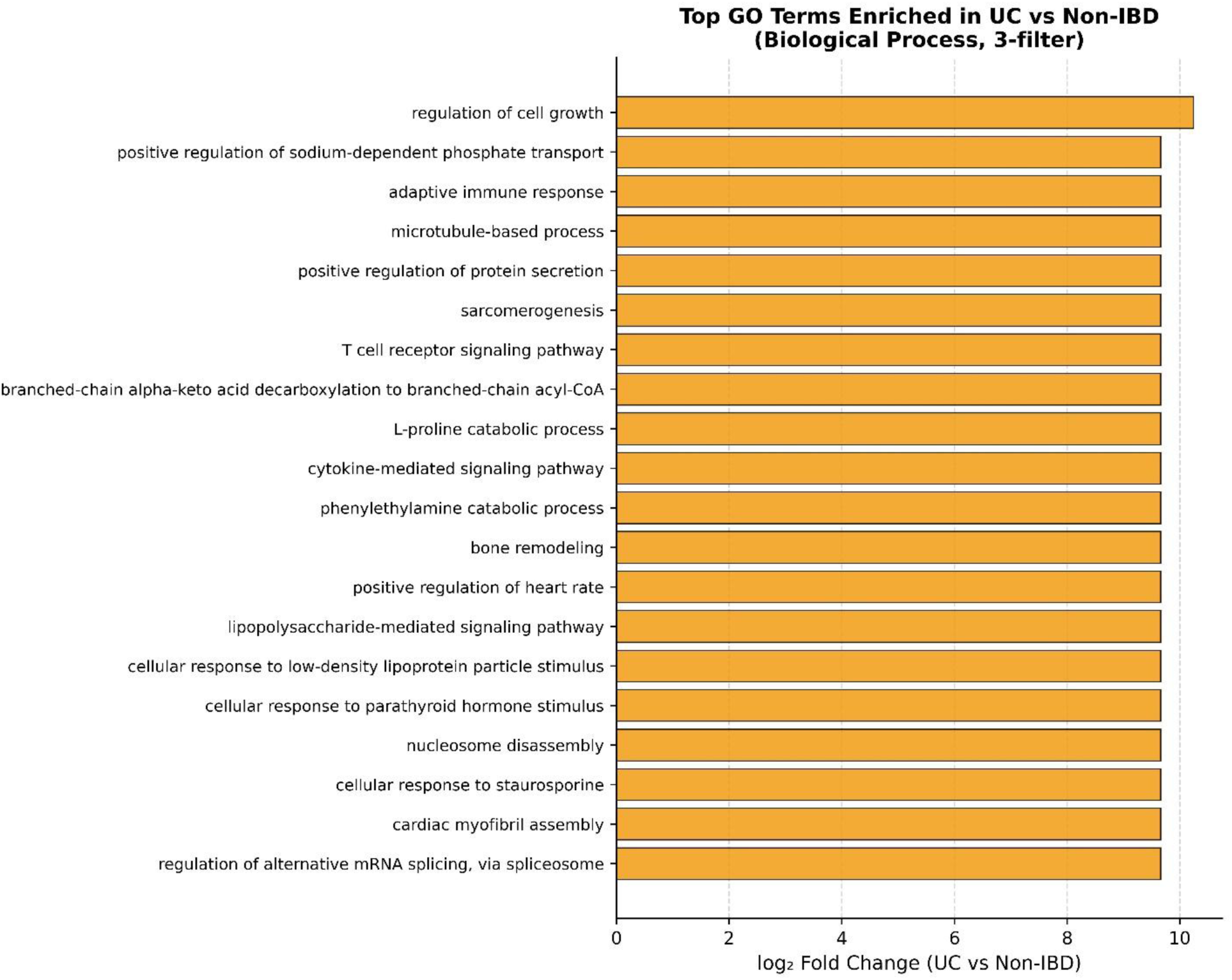
— Top 20 GO terms enriched in UC vs non-IBD.

**Figure 4D.**
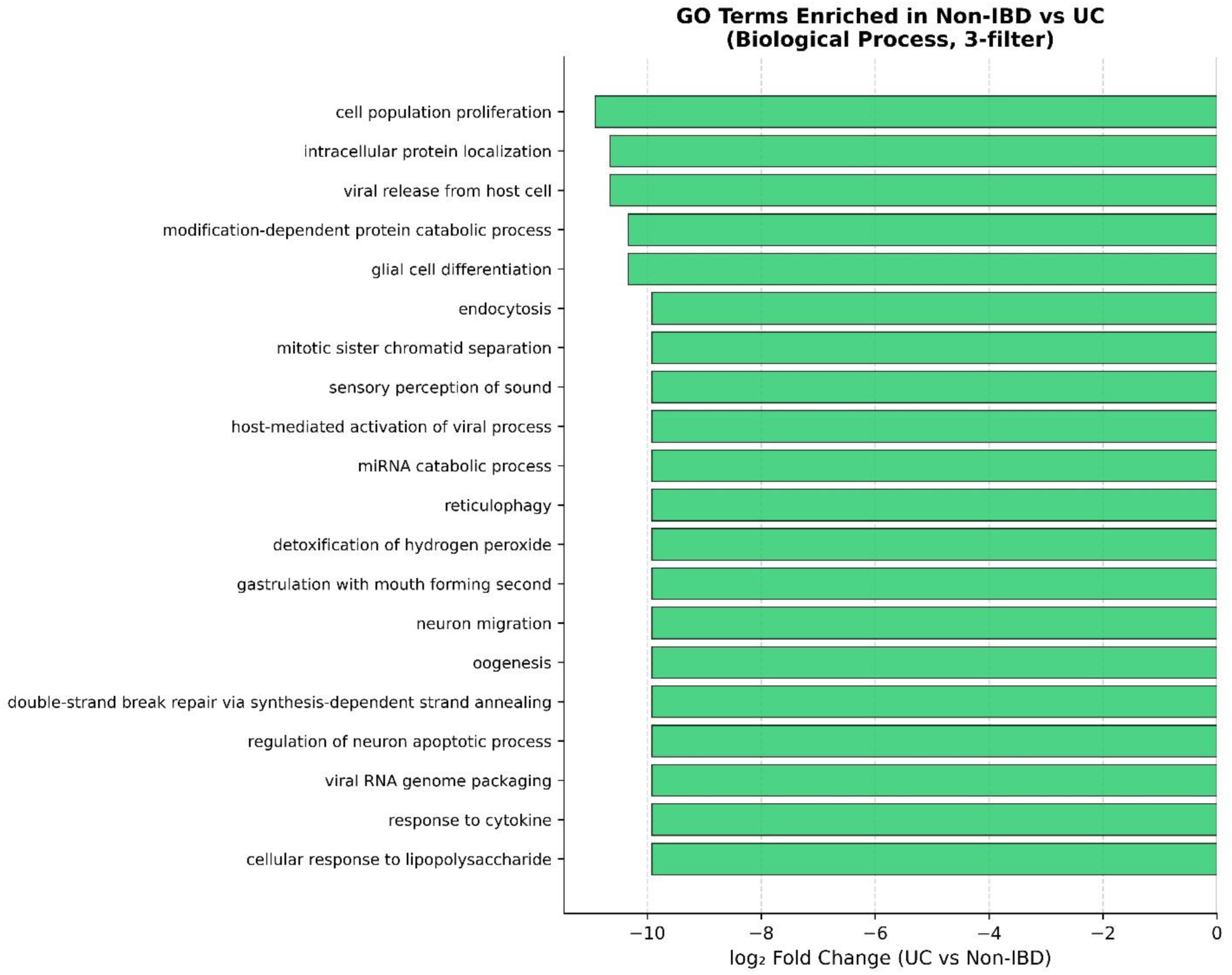
— Top 20 GO terms enriched in non-IBD vs UC.

#### 3.3.4 CD vs UC Direct Comparison

Direct comparison between CD and UC microbiomes confirmed differential functional representation between the two IBD subtypes (Figure 4E). Terms enriched in CD relative to UC included processes related to immune cell regulation and cellular stress responses. Figure 4F shows terms enriched in UC relative to CD. Direct CD vs UC comparison identified 743 differentially represented terms, with CD showing modest enrichment over UC (402 vs 341 terms).

**Figure 4E.**
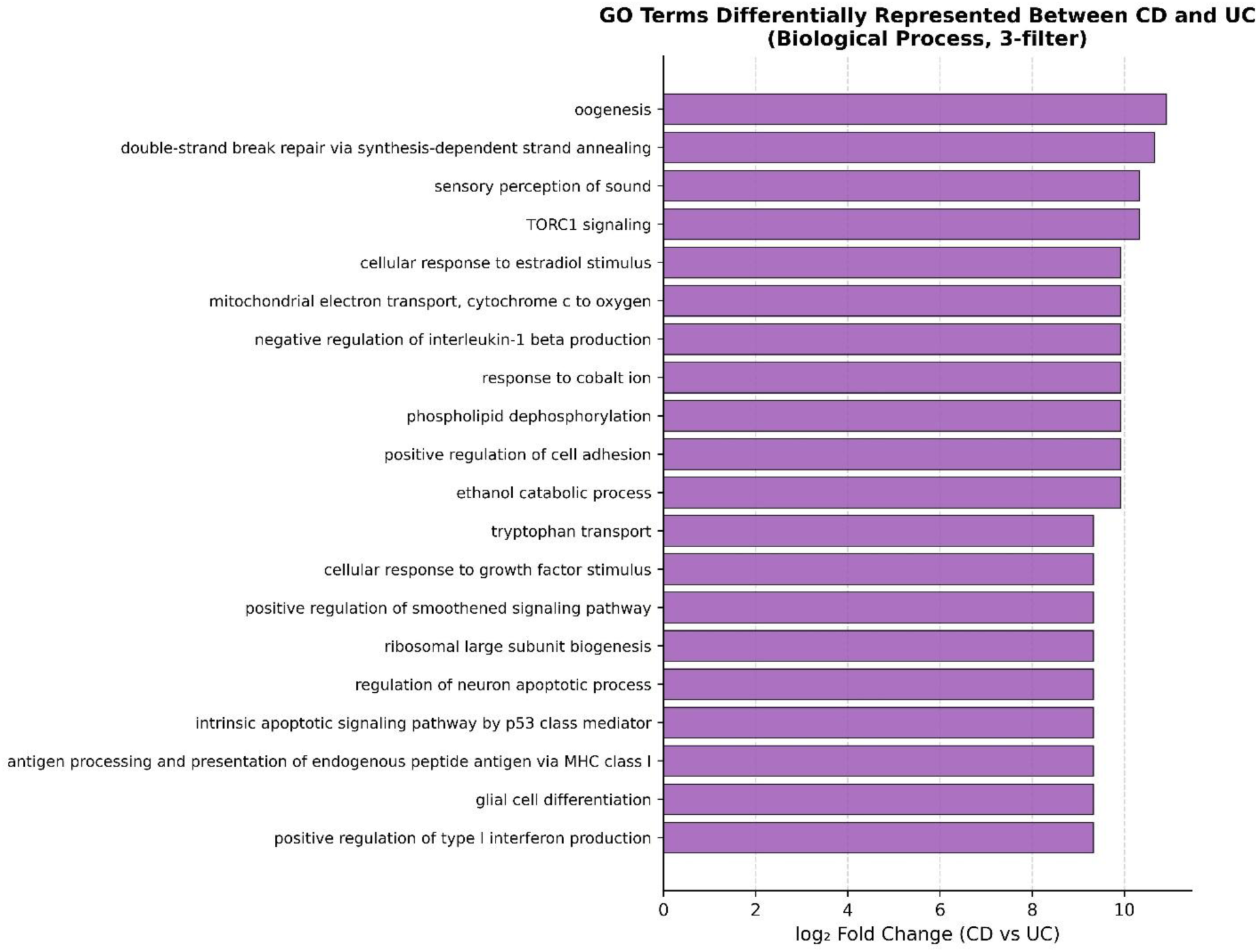
— GO terms differentially represented between CD and UC.

**Figure 4F.**
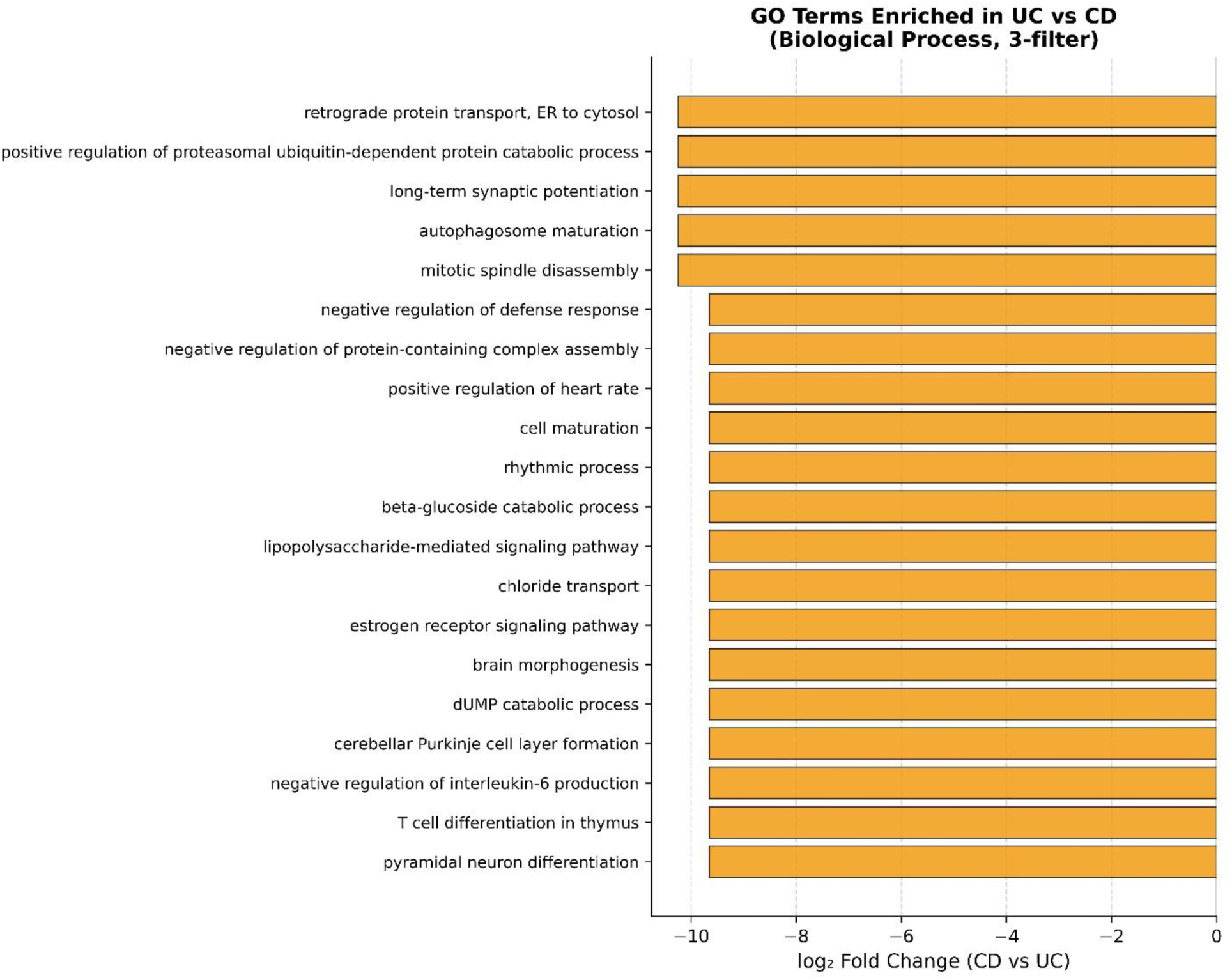
— GO terms enriched in UC vs CD.

Additionally, several neuronal and SRP-related terms were also found to be differentially expressed, which have been detailed in **Supplementary Files 2 and 3**, respectively and which have been discussed in detail in Section 3.11 with greater emphasis.

### 3.4 g:Profiler Enrichment Analyses

#### 3.4.1 Direct GO ID Enrichment

Statistical enrichment analysis using GO identifiers directly submitted to g:Profiler (human genome background) identified 189, 191, and 187 significantly enriched terms (p < 0.05) for nonIBD, CD, and UC groups respectively. The near-identical term counts and the dominance of high-level terms (term size 3,909–12,743) reflect broad functional convergence between the gut metagenome and human biology rather than disease-specific pathway enrichment (Figure 5A, no filter). Application of two-filter analysis (BP only; remove generic keywords) identified 43 differentially enriched terms in the CD vs nonIBD comparison and only 2 terms in UC vs nonIBD (Figure 5B). Three-filter analysis further refined the CD signal to 17 specific terms while UC retained 2 terms — ‘localization’ and ‘response to stimulus’ — which represent broad functional categories without specific disease relevance (Figure 5C). The stark contrast between CD (43 terms at 2-filter and 17 at 3-filter) and UC (2 terms at both 2-filter as well as 3-filter) provides additional evidence for greater functional divergence in CD.

**Figure 5A.**
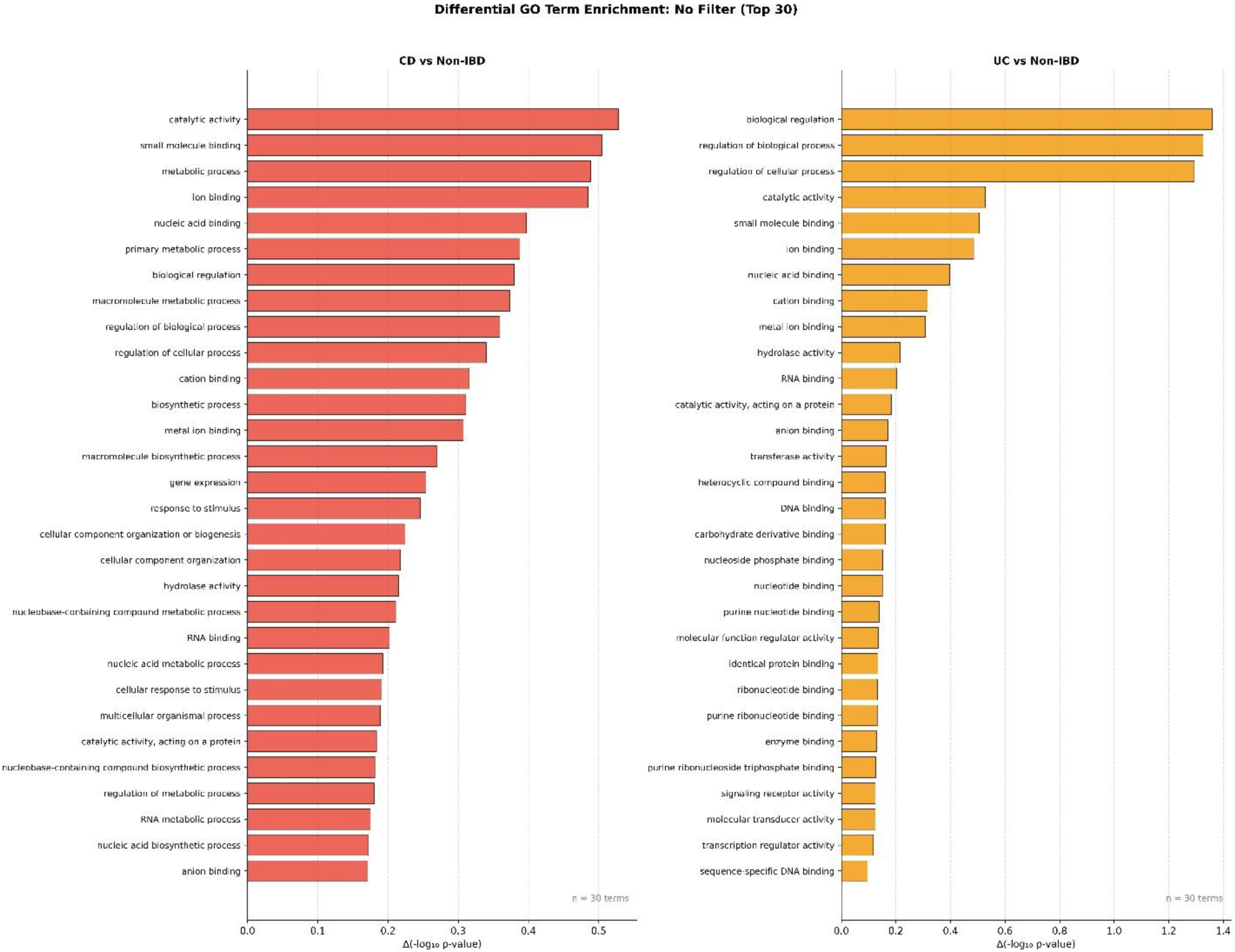
— No-filter g:Profiler enrichment (top 30).

**Figure 5B.**
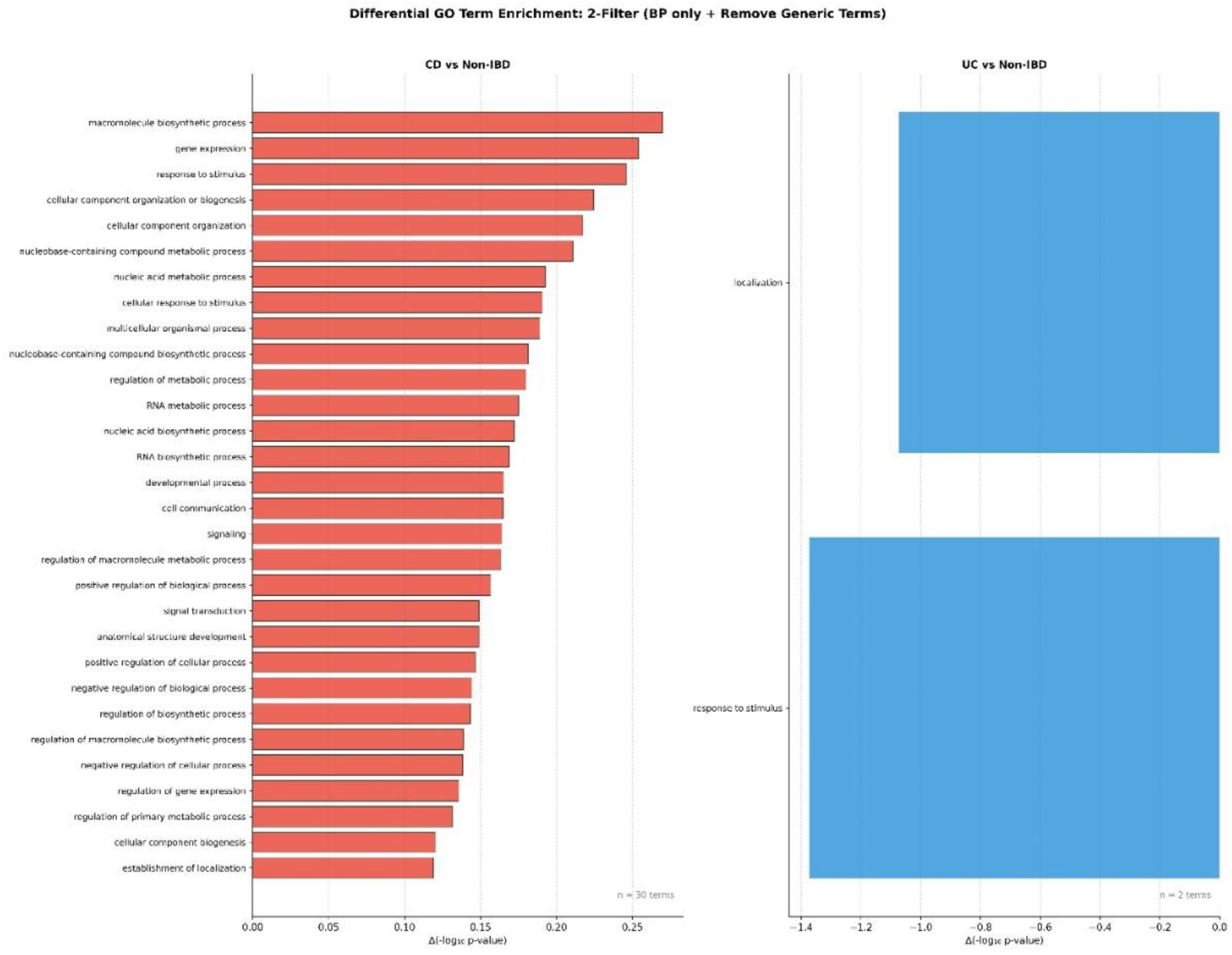
— 2-filter comparison.

**Figure 5C.**
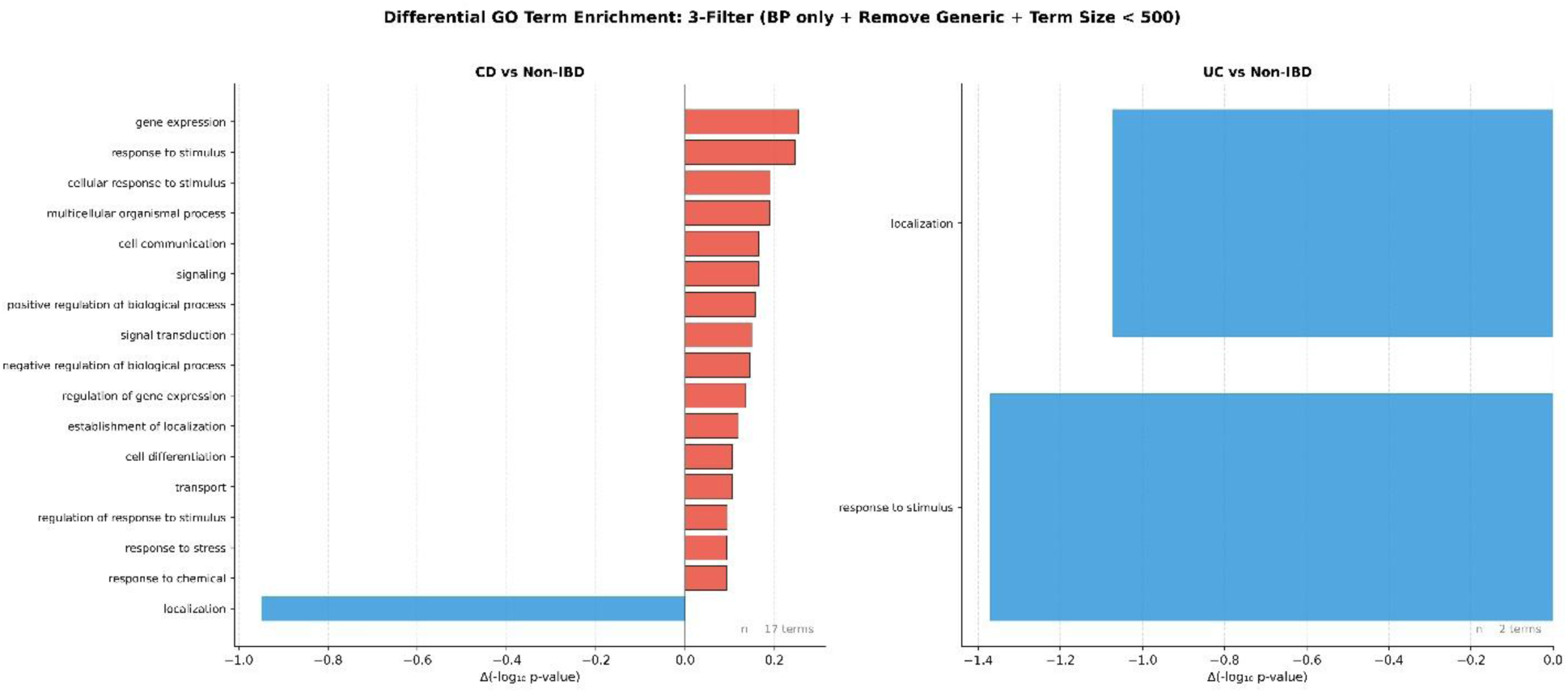
— 3-filter comparison.

#### 3.4.2 Ensembl Gene ID-based Enrichment and Gene-level Analysis

Conversion of GO identifiers from URIM to Ensembl human gene IDs via g:Profiler yielded 22,941, 22,938, and 22,939 unique gene IDs for nonIBD, CD, and UC respectively, representing near-complete coverage of the annotated human protein-coding genome. Differential gene analysis restricted to genes with |log₂FC| > 1 identified 0 genes enriched in CD vs nonIBD, 13 genes enriched in UC vs nonIBD, and sparse signals in other comparisons (Figs. 5D–5F). This near-complete human genome coverage indicates that the gut metagenome collectively samples a very broad range of human functional biology, and that the relevant discriminator between diagnostic groups is quantitative frequency of GO term representation rather than categorical presence or absence of specific gene functional orthologs.

Targeted g:Profiler analysis of the differentially represented genes identified phosphotyrosine residue binding (GO:0001784), protein phosphorylated amino acid binding (GO:0045309), and phosphoprotein binding (GO:0051219) as depleted in both CD and UC relative to healthy controls (**Table 5**). This convergent depletion of phosphorylation-related functional annotations in both IBD subtypes may reflect the loss of specific commensals contributing these functional capabilities.

**Figure 5D.**
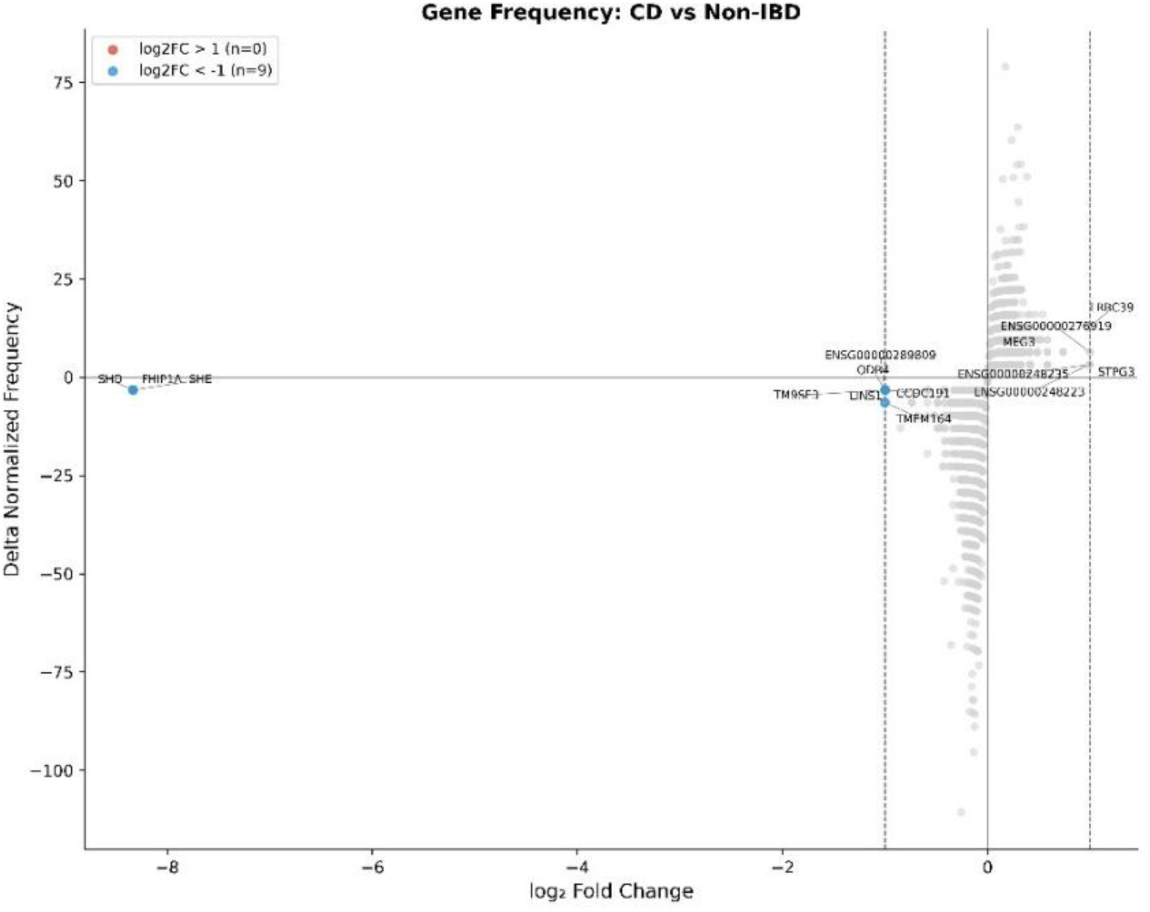
Volcano plot for CD vs non-IBD comparison.

**Figure 5E.**
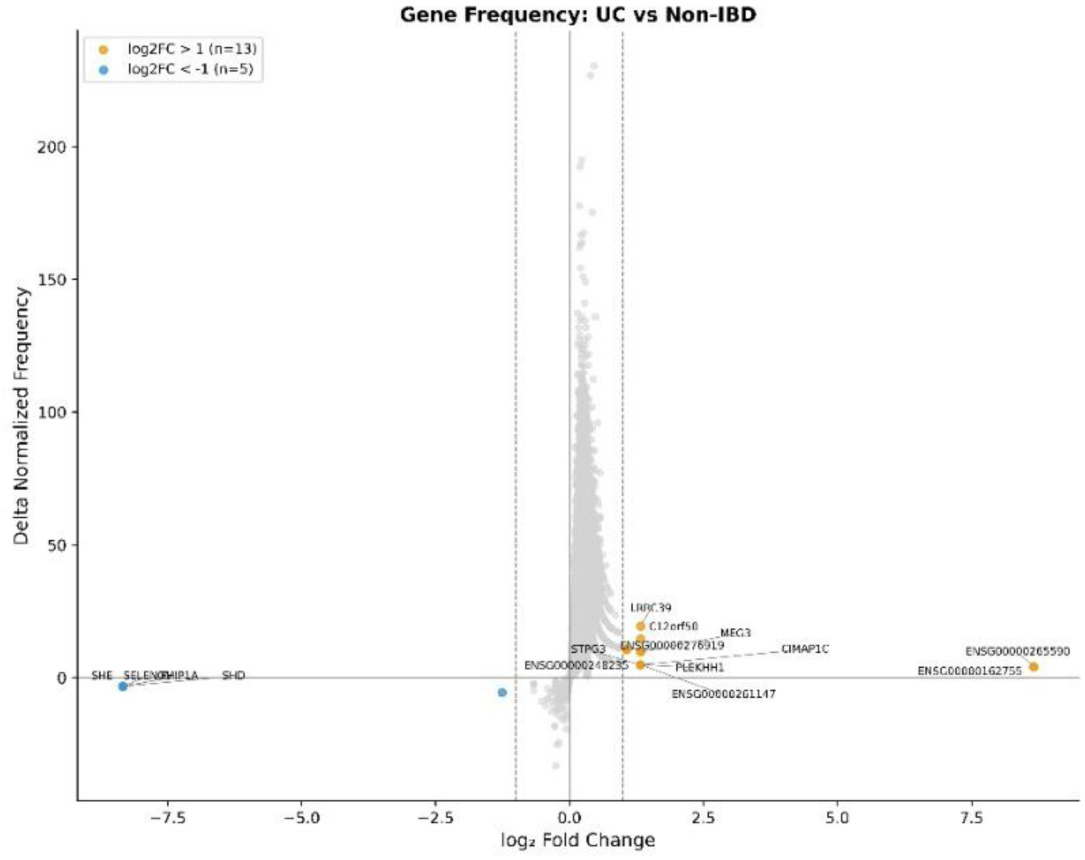
Volcano plot for UC vs non-IBD comparison.

**Figure 5F.**
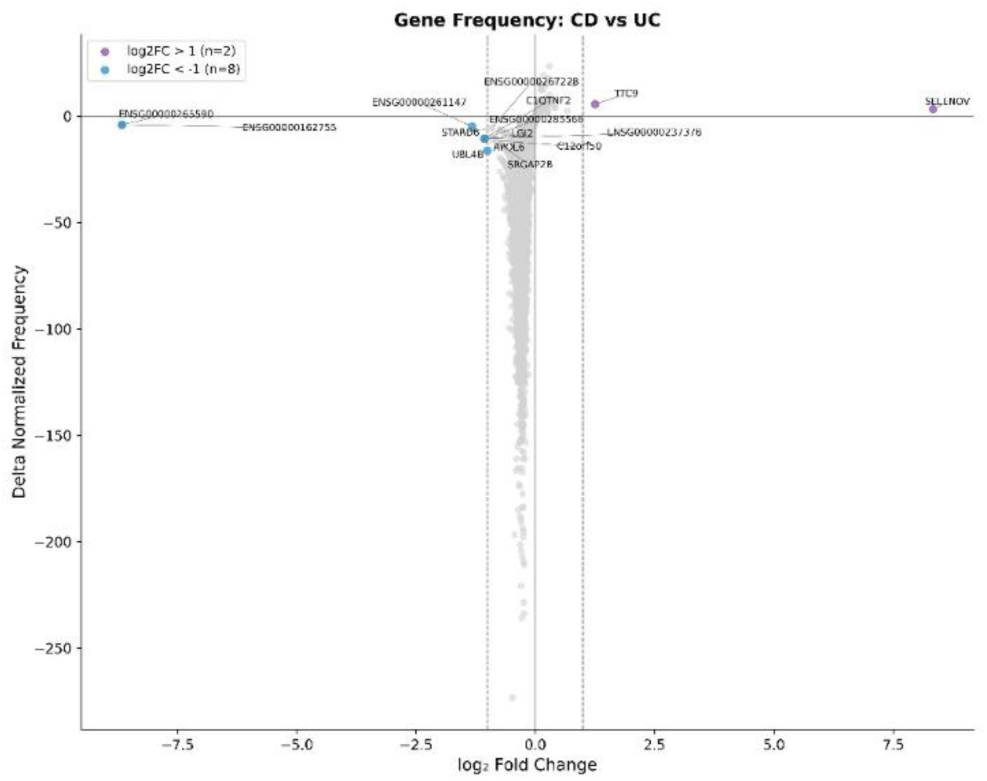
Volcano plot for CD vs UC comparison.

**Table 5.** Differentially enriched GO terms identified by g:Profiler analysis of Ensembl gene IDs derived from GO ID conversion (p < 0.05).

| Comparison | Direction | Significant GO Terms ($p < 0.05$) | Key terms |
| --- | --- | --- | --- |
| CD vs nonIBD | Enriched in CD | 0 | — |
| CD vs nonIBD | Depleted in CD | 3 | Phosphotyrosine residue binding, Protein phosphorylated amino acid binding, Phosphoprotein binding |
| UC vs nonIBD | Enriched in UC | 1 | Negative regulation of cell growth |
| UC vs nonIBD | Depleted in UC | 3 | Phosphotyrosine residue binding, Protein phosphorylated amino acid binding, Phosphoprotein binding |
| CD vs UC | Enriched in CD | 0 | — |
| CD vs UC | Depleted in CD | 0 | — |

*Note: The convergent depletion of phosphorylation-related binding terms in both CD and UC relative to healthy controls suggests loss of specific commensals contributing these functional capabilities during IBD-associated dysbiosis.*

### 3.5 PFAM Domain Enrichment Reveals Differential Protein Family Architecture

A total of 3,991 unique PFAM protein families were detected across the combined dataset. After per-sample normalization, differential enrichment analysis identified multiple protein families with divergent representation between diagnostic groups.

The top PFAM domains enriched in CD relative to healthy controls included LpoA family (log₂FC = 10.65), Class I-like SAM-binding methyltransferase superfamily CmoB family (log₂FC = 10.33), Major facilitator superfamily oligosaccharide transporter family (log₂FC = 10.33), Peptidase C56 family (log₂FC = 10.33), and SMC family RAD50 subfamily (log₂FC = 10.33) (Figure 6A). The enrichment of RAD50 subfamily — a DNA repair protein family — and peptidase families in CD microbiomes is noteworthy, as these domains are present in eukaryotic organisms and their enrichment in microbial proteins may reflect either horizontal gene transfer from host cells or the expansion of taxa harboring eukaryote-like domain architectures.

**Figure 6A.**
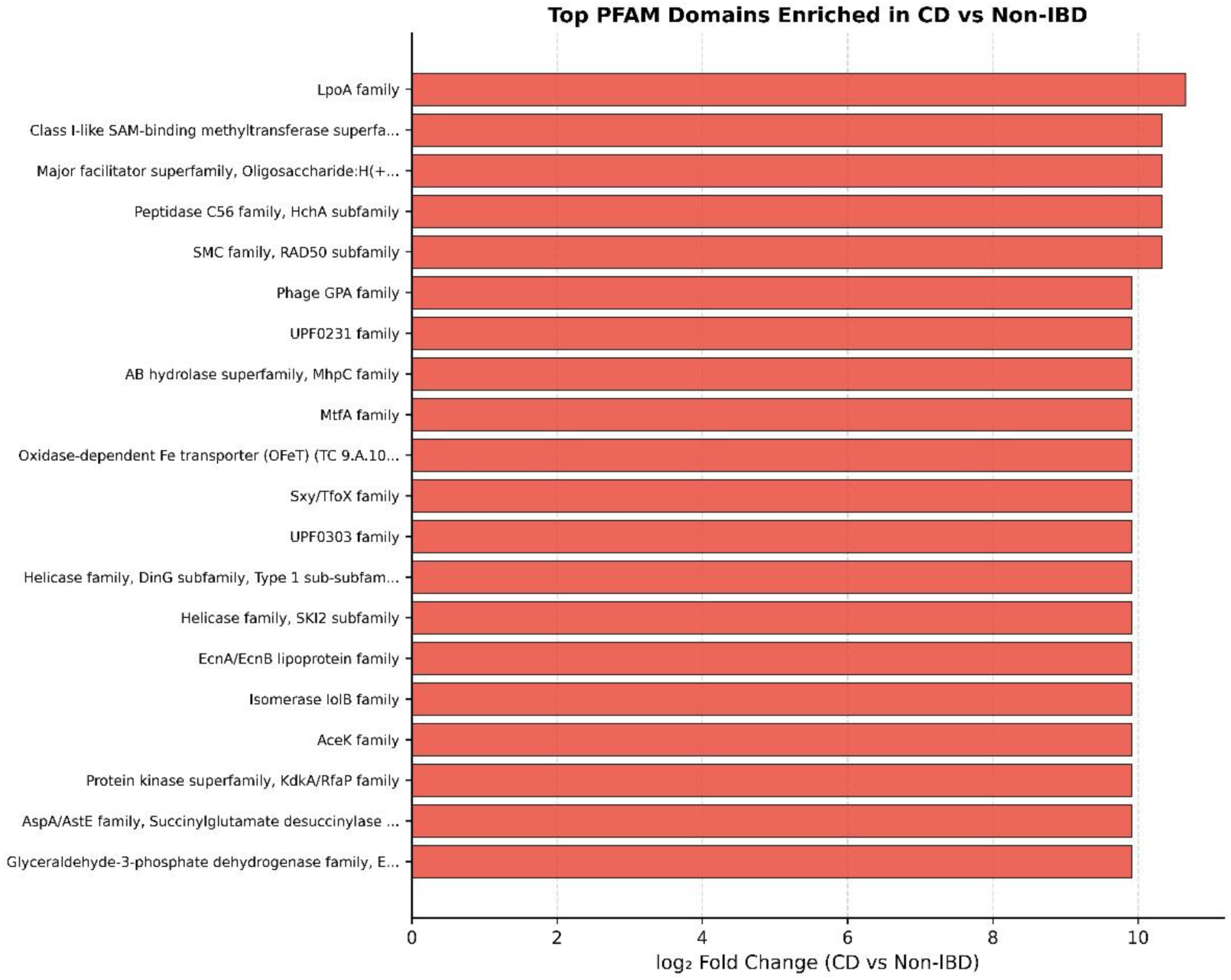
— CD-enriched PFAM domains.

Healthy non-IBD microbiomes were enriched in distinct PFAM families relative to CD, including RTX prokaryotic toxin family (log₂FC = −10.65), Class-III pyridoxal-phosphate-dependent aminotransferase family (log₂FC = −10.33), and Inovirus G1P protein family (log₂FC = −9.92) (Figure 6B). UC-enriched domains included UPF0159 family (log₂FC = 10.66), CofE family (log₂FC = 10.24), and Glycosyltransferase 2 family GalNAc-T subfamily (log₂FC = 10.24), the latter being of particular interest given the known role of GalNAc-type glycosylation in host-microbe interactions (Figure 6C) **(Lei et al., 2025).** Protective PFAM domains enriched in healthy vs UC are shown in Figure 6D.

**Figure 6B.**
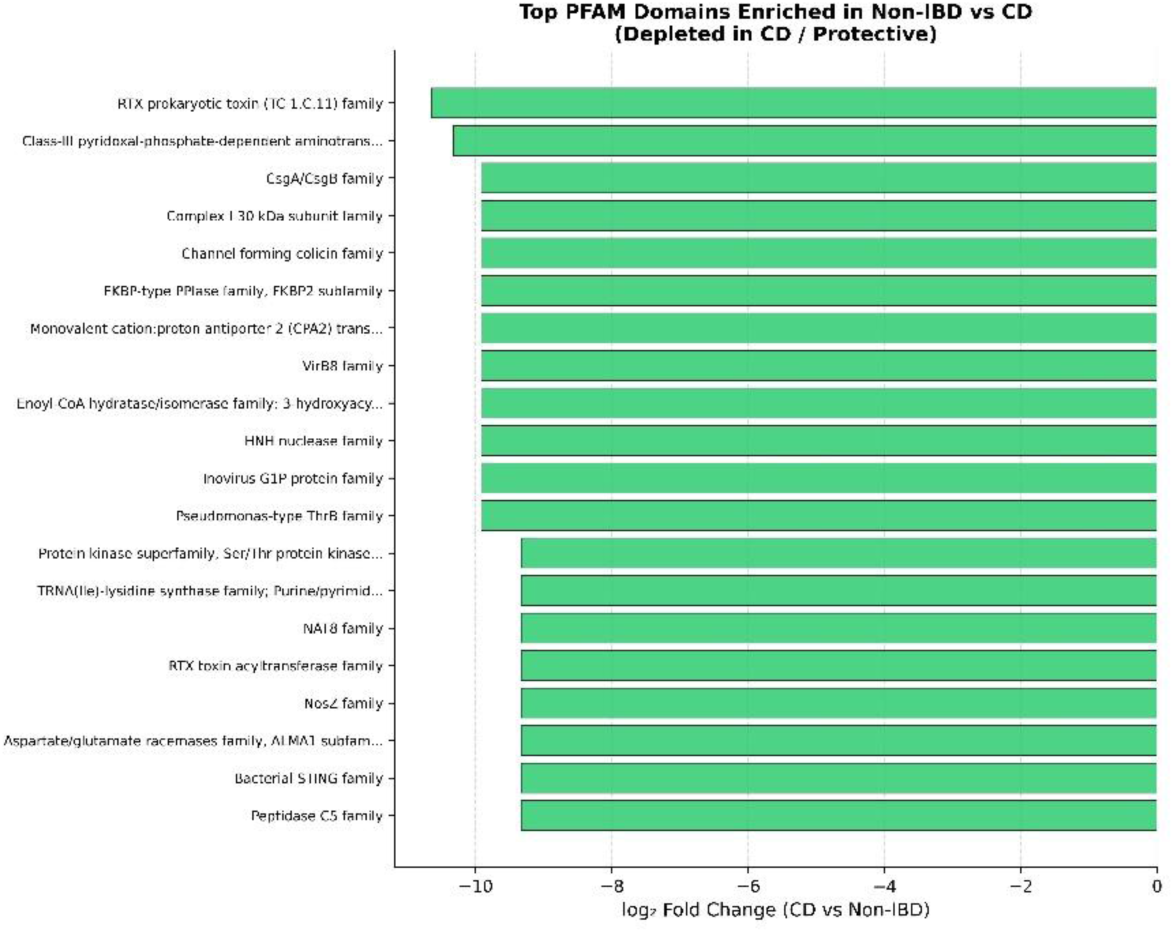
— Domains enriched in non-IBD vs CD.

**Figure 6C.**
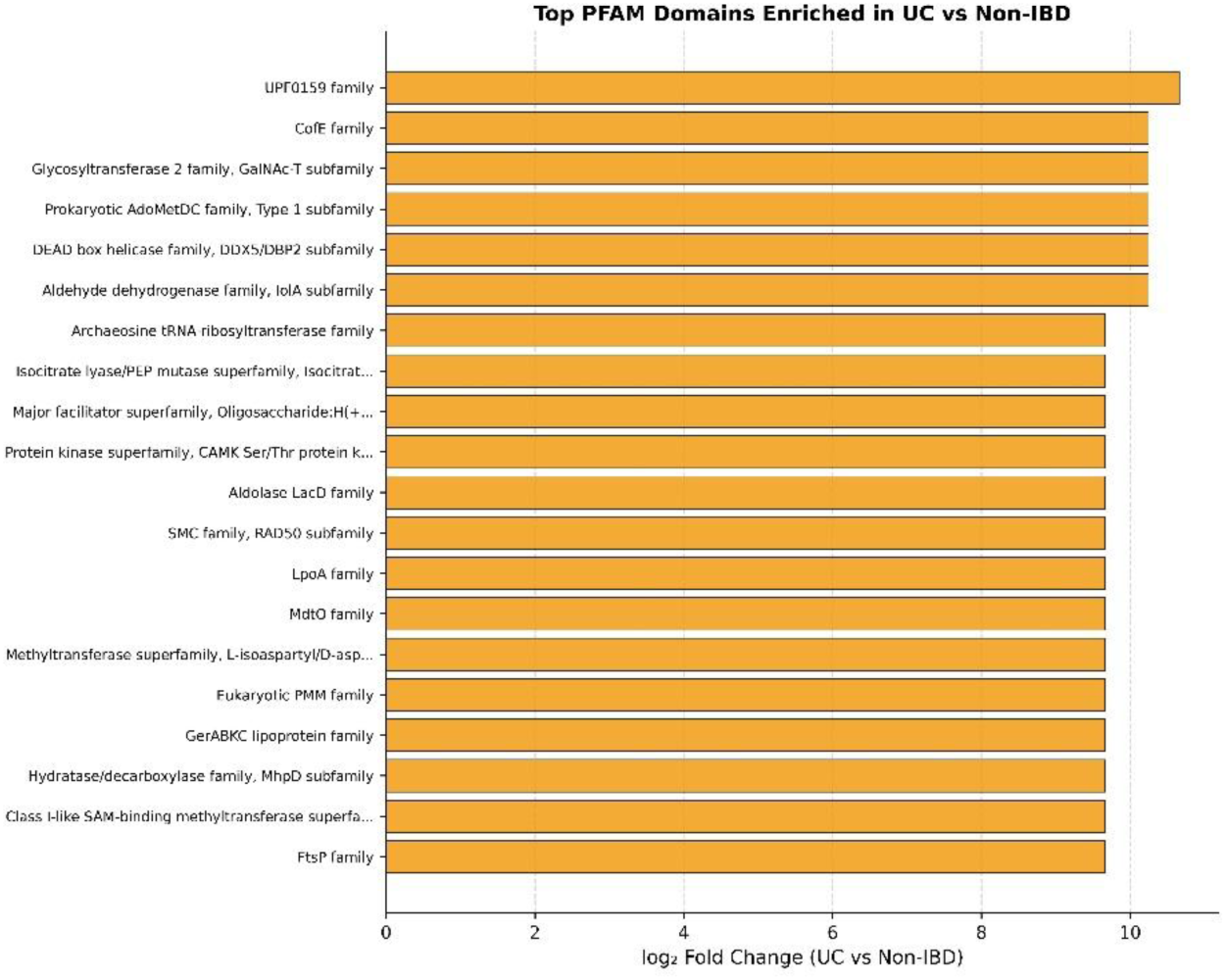
— UC-enriched PFAM domains.

**Figure 6D.**
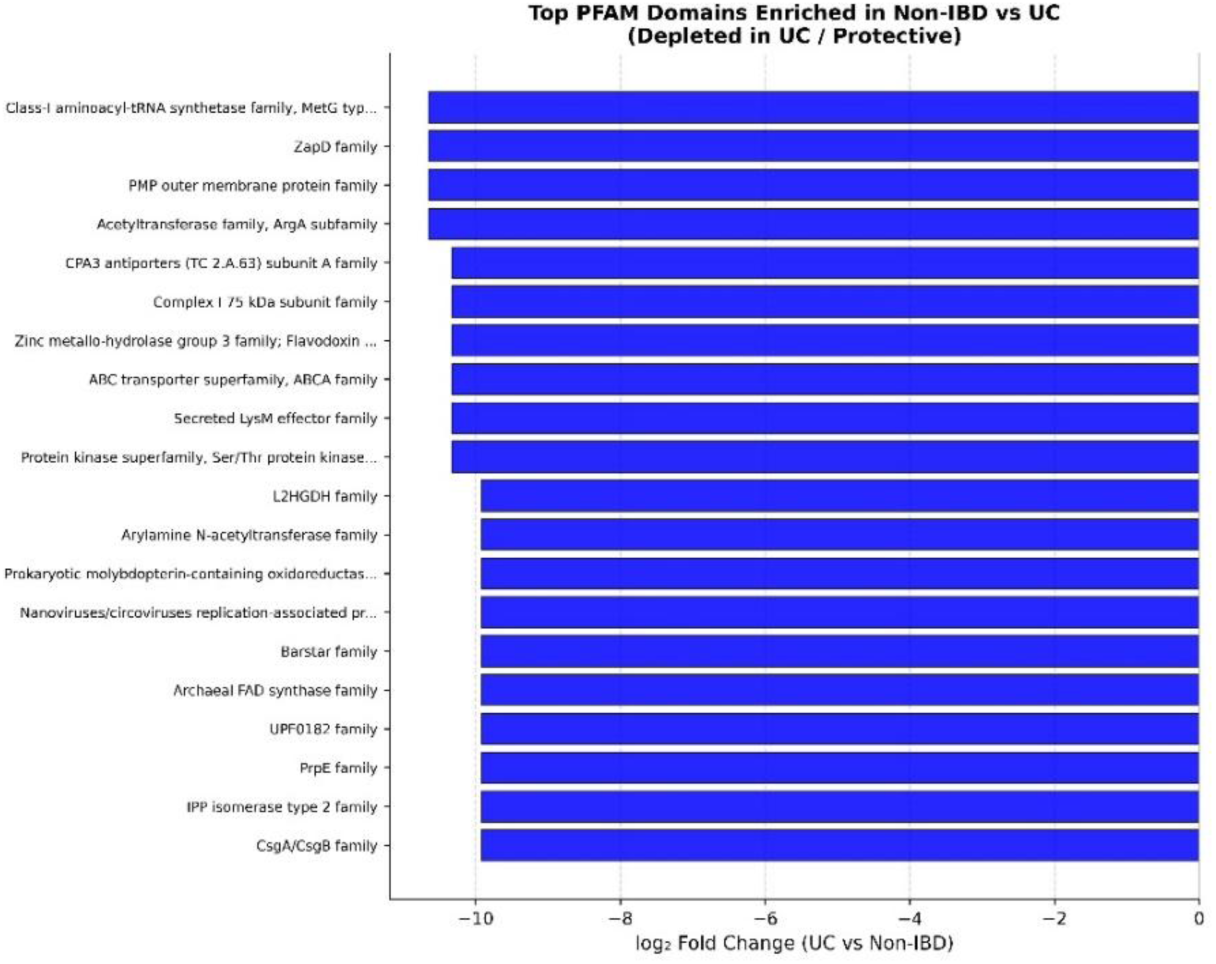
— Domains enriched in non-IBD vs UC.

### 3.6 Sequence-level Mimicry: Microbial Proteins with Direct Human Sequence Homology

Isolation of DIAMOND hits to Homo sapiens proteins identified 905 human-matching hits (238 unique human UniProt accessions) in the nonIBD group, 903 hits (219 accessions) in CD, and 544 hits (187 accessions) in UC.

#### 3.6.1 Normalization Strategy and Group Comparisons

Human sequence mimicry rates were normalized by mean ORFs per sample within each group to account for the substantial difference in per-sample ORF counts between groups (nonIBD: 58,388 vs CD/UC: ∼40,000 ORFs per sample). CD microbiomes showed the highest normalized human mimicry rate (2,250 per 100,000 ORFs), followed by nonIBD (1,550) and UC (1,338) (Figure 7A). Percentage identity of microbial-to-human alignments was low and consistent across groups (mean 34–35%) (Figure 7B).

**Figure 7.**
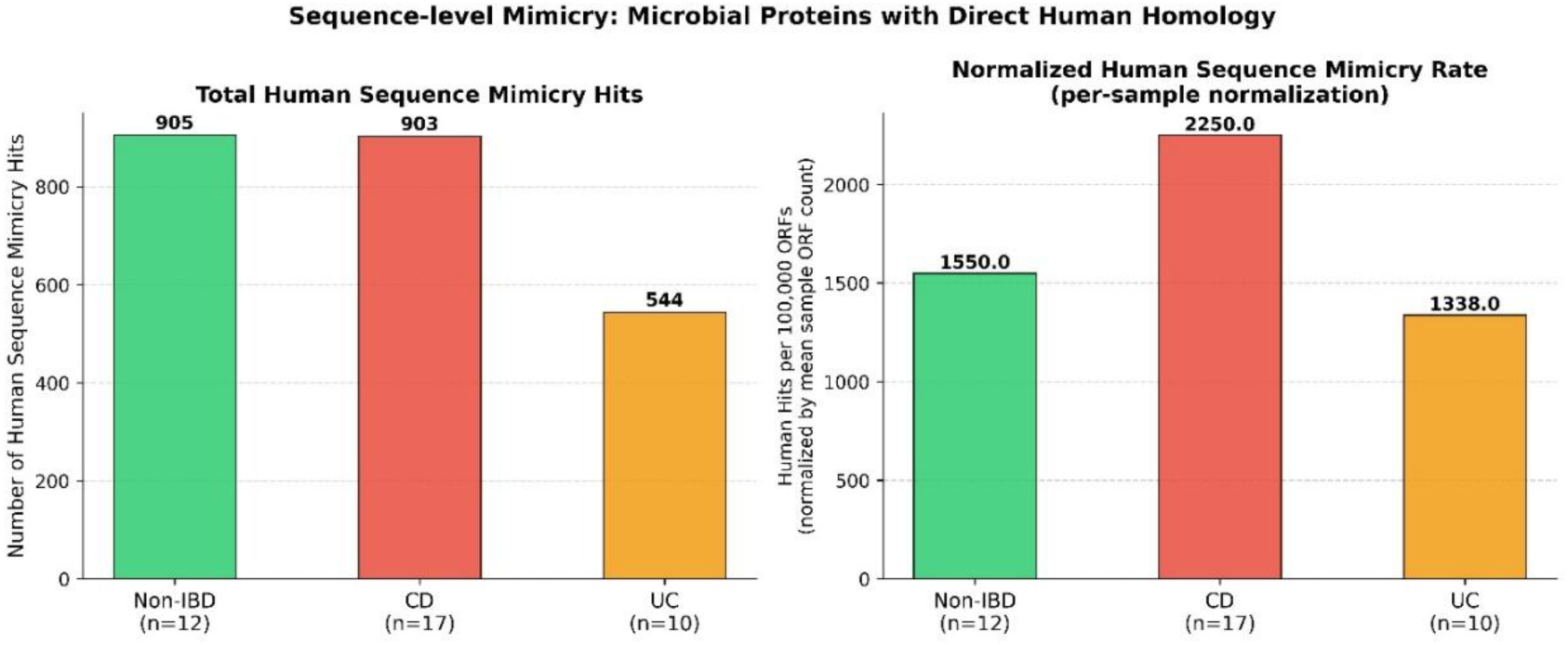
A Per-sample mean ORF normalization.

**Figure 7B.**
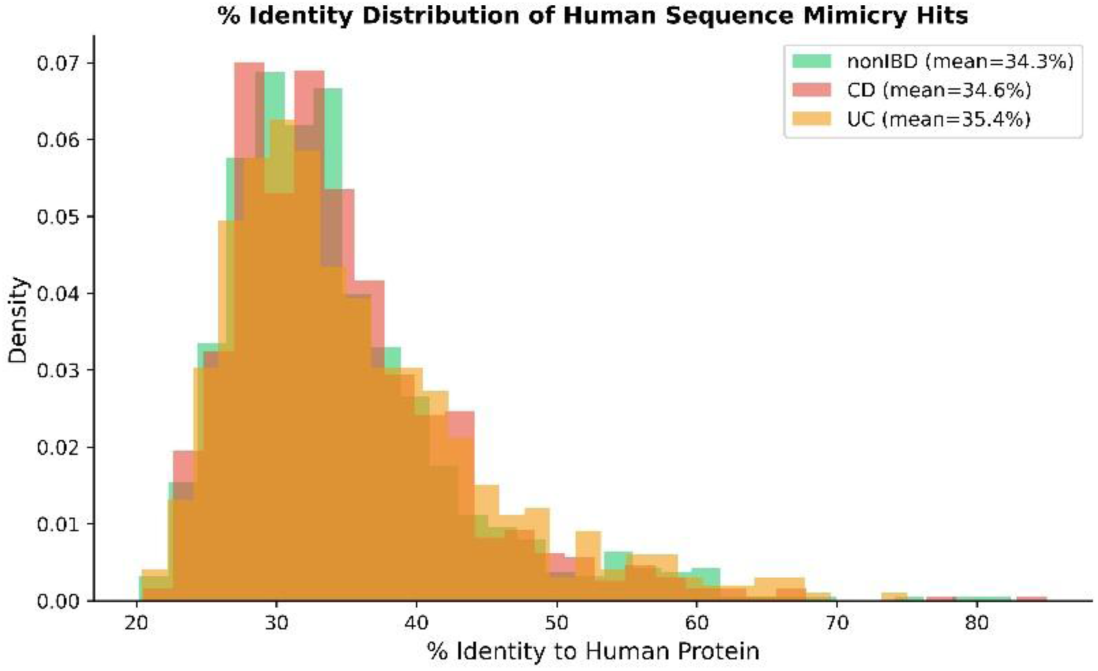
Percentage identity of microbial-to-human alignments

#### 3.6.2 Sequence Mimicry Rates and Human Protein Targets

UniProt accession IDs extracted from the human-filtered DIAMOND output were submitted to URIM, yielding 238, 219, and 187 unique human UniProt accessions for nonIBD, CD, and UC respectively, confirming that CD microbiomes access a broader repertoire of human sequence homologs than UC.

The most frequently matched human proteins across all diagnostic groups included NAGLU (alpha-N-acetylglucosaminidase), HGSNAT (heparan sulfate glucosamine N-acetyltransferase), SIAE (sialate O-acetylesterase), PIF1 (DNA helicase), FUCA2 (alpha-L-fucosidase 2), and PEPD (Xaa-Pro dipeptidase) (Figure 7C). NAGLU and HGSNAT are lysosomal enzymes involved in glycosaminoglycan catabolism, SIAE participates in sialic acid metabolism, and FUCA1/FUCA2 are fucosidases — collectively, these targets suggest preferential microbial mimicry of human carbohydrate-processing enzymes, consistent with the importance of glycan utilization in gut microbial ecology **[Tailford et al., 2015; Fedele, 2015].** CD microbiomes showed notably elevated mimicry rates for NAGLU (CD: 154.5, nonIBD: 78.8, UC: 46.7 per 100,000 ORFs) and CDA (cytidine deaminase; CD: 74.7, nonIBD: 29.1, UC: 24.6), while SLC5A11 (sodium-dependent glucose transporter) was detected only in nonIBD and CD but was essentially absent in UC.

**Figure 7C.**
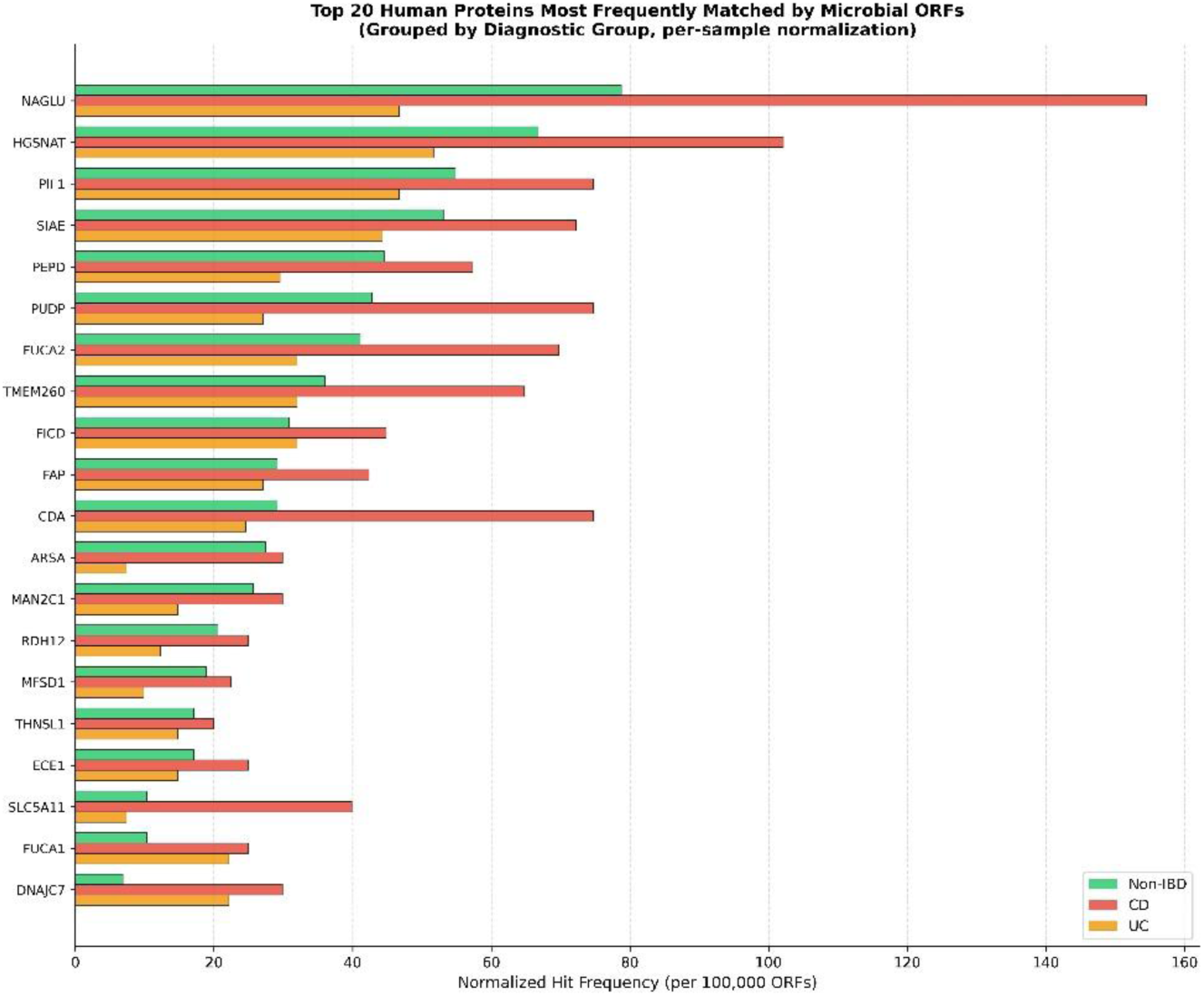
— Grouped horizontal bar chart of top 20 human protein targets.

### 3.7 Taxonomic Profiling: Dysbiosis Patterns Corroborate Functional Mimicry Signals

MetaPhlAn3 taxonomic profiling of the 39 study samples revealed distinct species-level composition between diagnostic groups, with CD showing more pronounced dysbiosis than UC relative to healthy controls (Figure 8A). Full species-level abundance comparisons across all three pairwise contrasts are provided in **Supplementary File 4**.

**Figure 8A.**
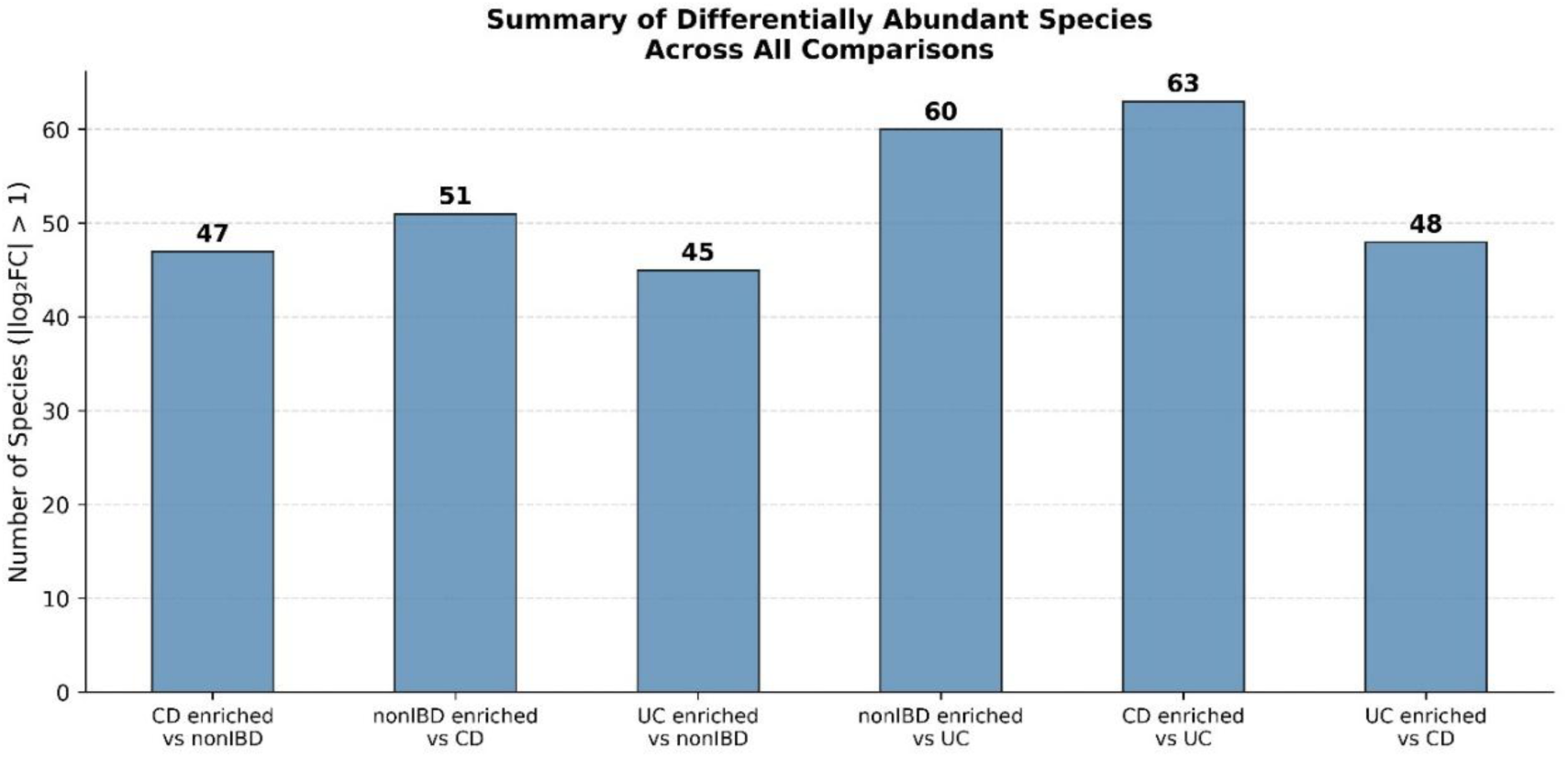
— Summary of differentially abundant species across all comparisons.

#### 3.7.1 CD-enriched Species

CD samples were enriched in several known pathobionts relative to healthy controls (Figure 8B). The most strongly enriched species included *Streptococcus thermophilus* (log₂FC = 7.28, present exclusively in CD), *Clostridium butyricum* (log₂FC = 6.21, CD-exclusive), *Klebsiella michiganensis* (log₂FC = 6.07), *Haemophilus parainfluenzae* (log₂FC = 5.96), *Bacteroides sp. CAG_144* (log₂FC = 5.04), *Acidaminococcus intestini* (log₂FC = 5.04), *Bacteroides fragilis* (log₂FC = 4.19), and *Clostridium perfringens* (log₂FC = 3.30). The enrichment of oral-origin bacteria (*S. thermophilus, H. parainfluenzae*) in the CD gut is consistent with the oral-gut microbial translocation hypothesis of CD pathogenesis **[Sohn et al., 2023].**

**Figure 8B.**
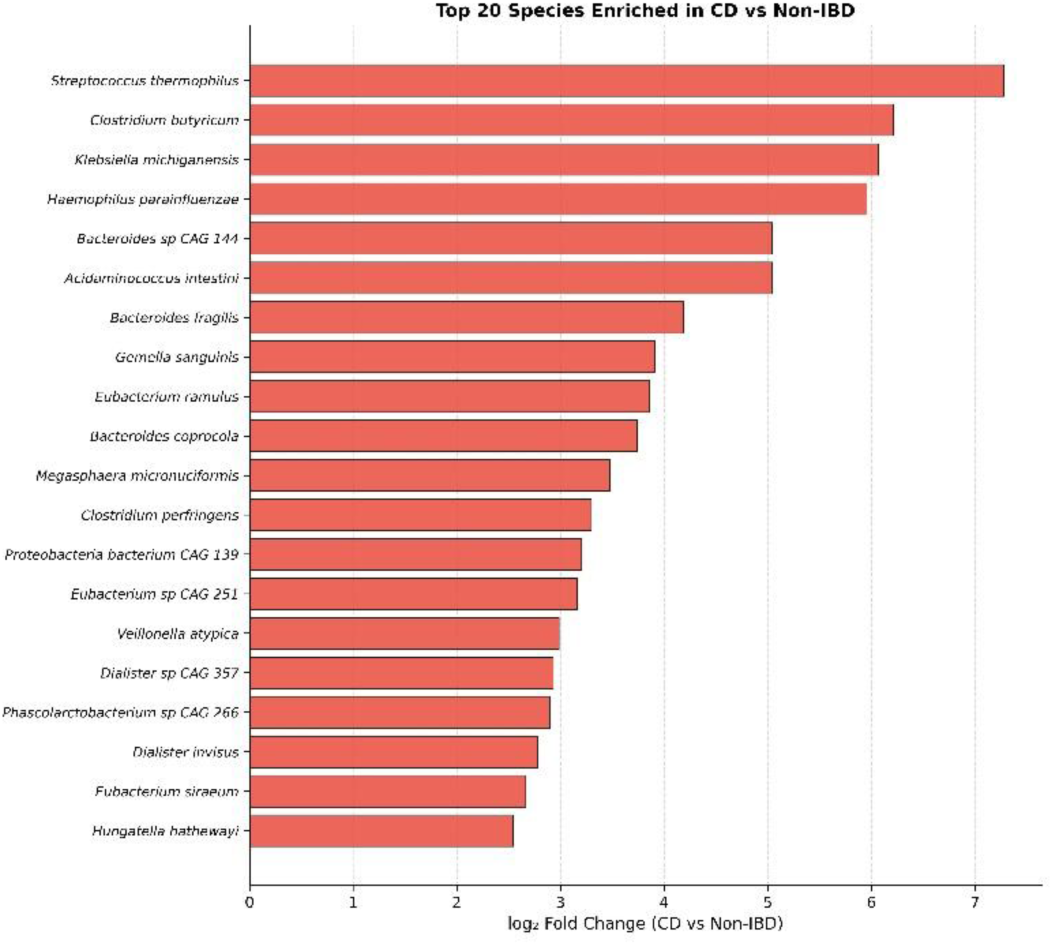
— Top 20 species enriched in CD vs non-IBD.

#### 3.7.2 Healthy-enriched (Protective) Species

Healthy non-IBD microbiomes were enriched in species associated with intestinal homeostasis, including *Bacteroides plebeius* (log₂FC = −7.14 in CD), *Akkermansia muciniphila* (log₂FC = −5.65 in CD), *Bacteroides finegoldii* (log₂FC = −6.43 in CD), and *Phascolarctobacterium succinatutens* (log₂FC = −5.31 in CD) (Figure 8C). *Akkermansia muciniphila* in particular is a well-established keystone commensal associated with mucosal barrier integrity, and its depletion in CD is consistent with prior reports **[Zheng et al., 2023].** These protective commensals were similarly depleted in UC relative to nonIBD, though the magnitude of depletion was generally smaller (Figure 8D).

**Figure 8C.**
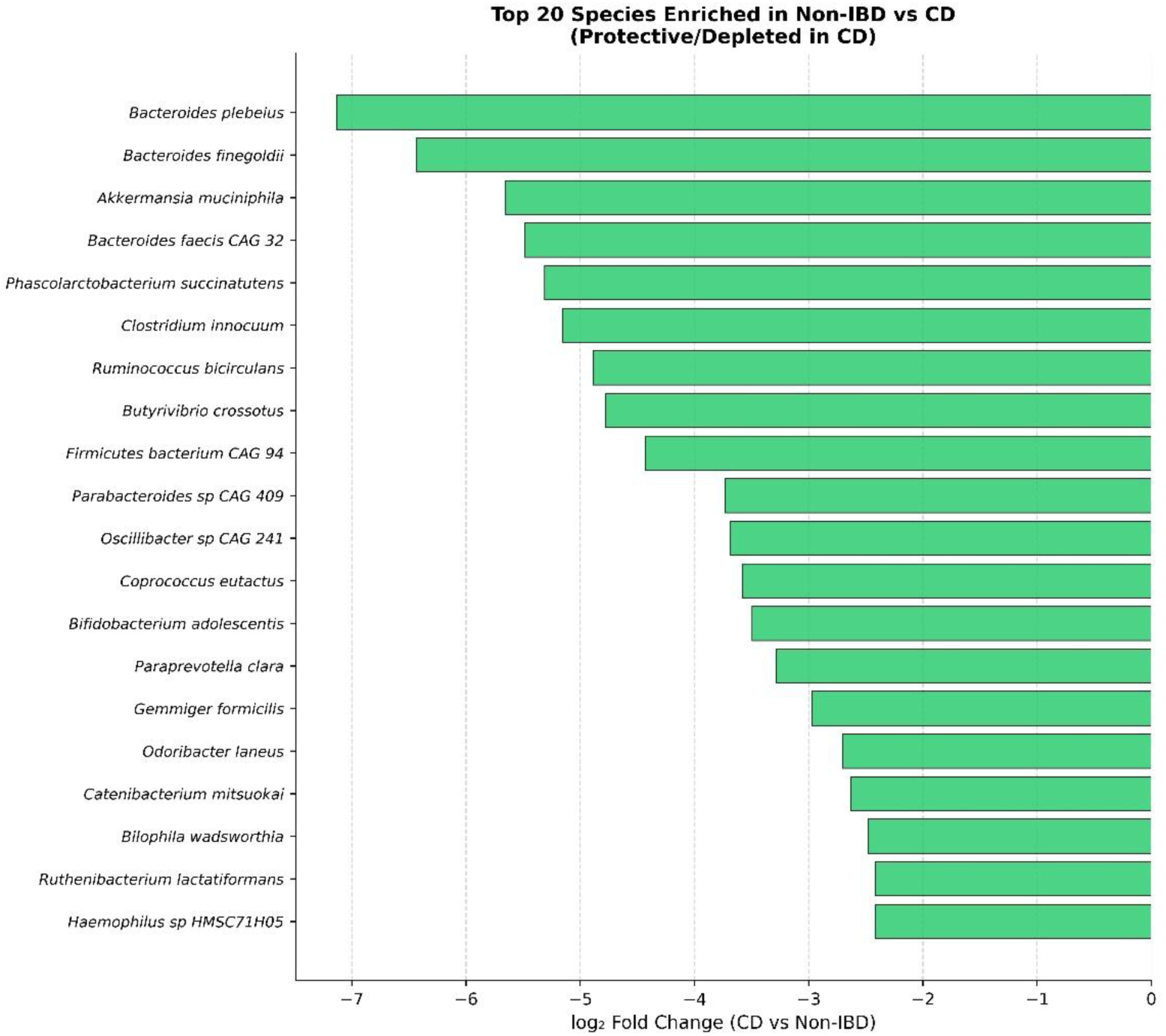
— Top 20 species enriched in non-IBD vs CD.

**Figure 8D.**
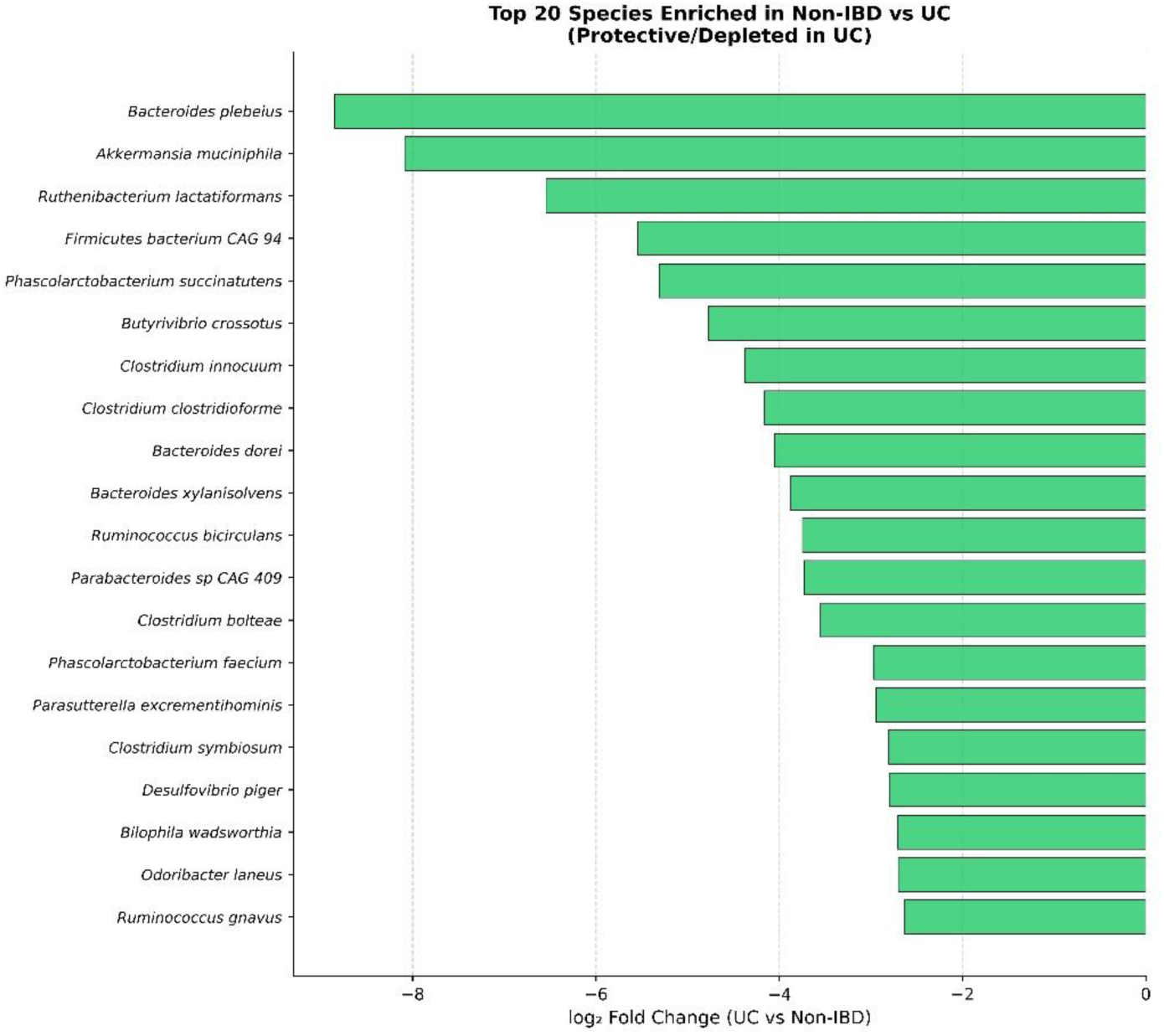
— Top 20 species enriched in non-IBD vs UC.

#### 3.7.3 UC-specific Dysbiosis

UC samples showed a distinct but less extreme dysbiosis profile compared to CD (Figure 8E). The summary comparison (Figure 8A) quantifies the differential abundance: CD showed more species significantly enriched relative to nonIBD than UC, consistent with the greater functional perturbation observed across all mimicry analyses. CD vs UC direct comparison further illustrated the taxonomic differences between the two IBD subtypes (Figure 8F-G), with several species exclusively or predominantly enriched in one disease subtype.

**Figure 8E.**
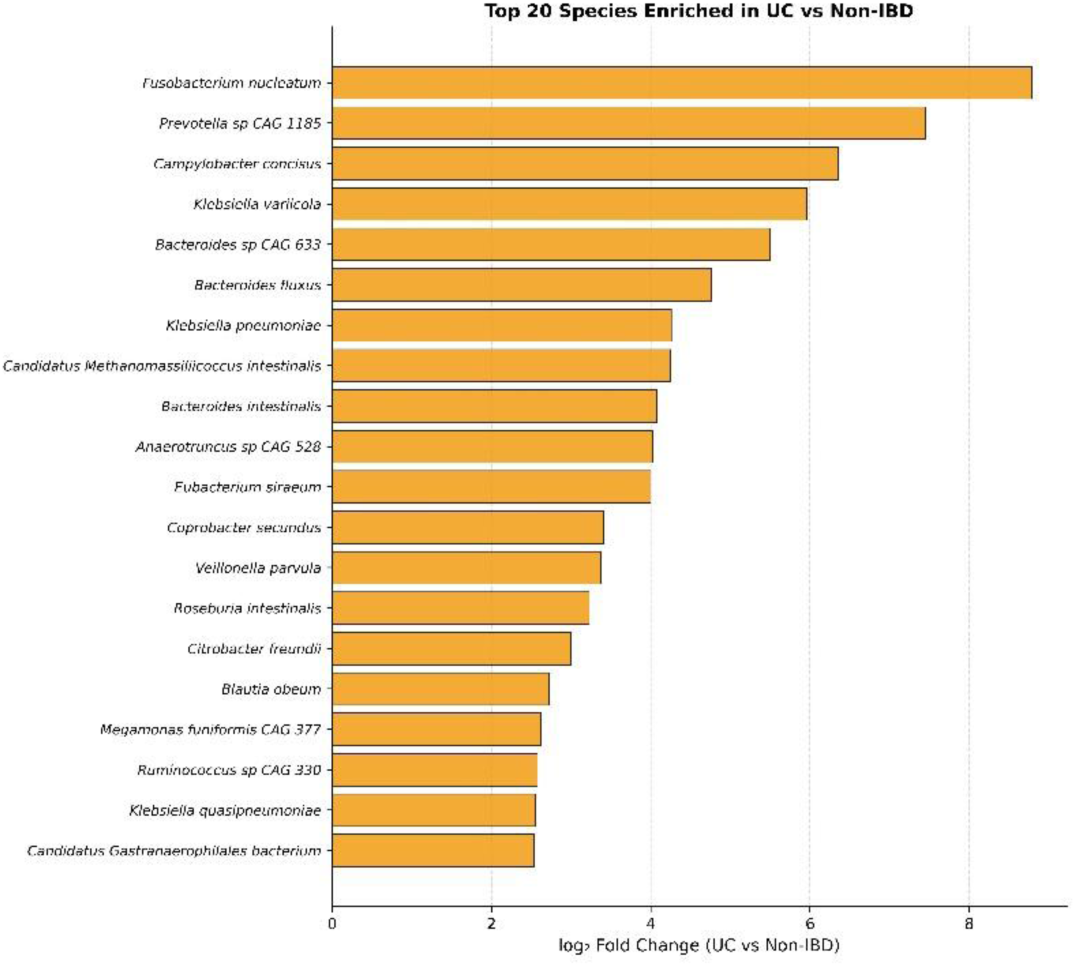
— Top 20 species enriched in UC vs non-IBD.

**Figure 8F.**
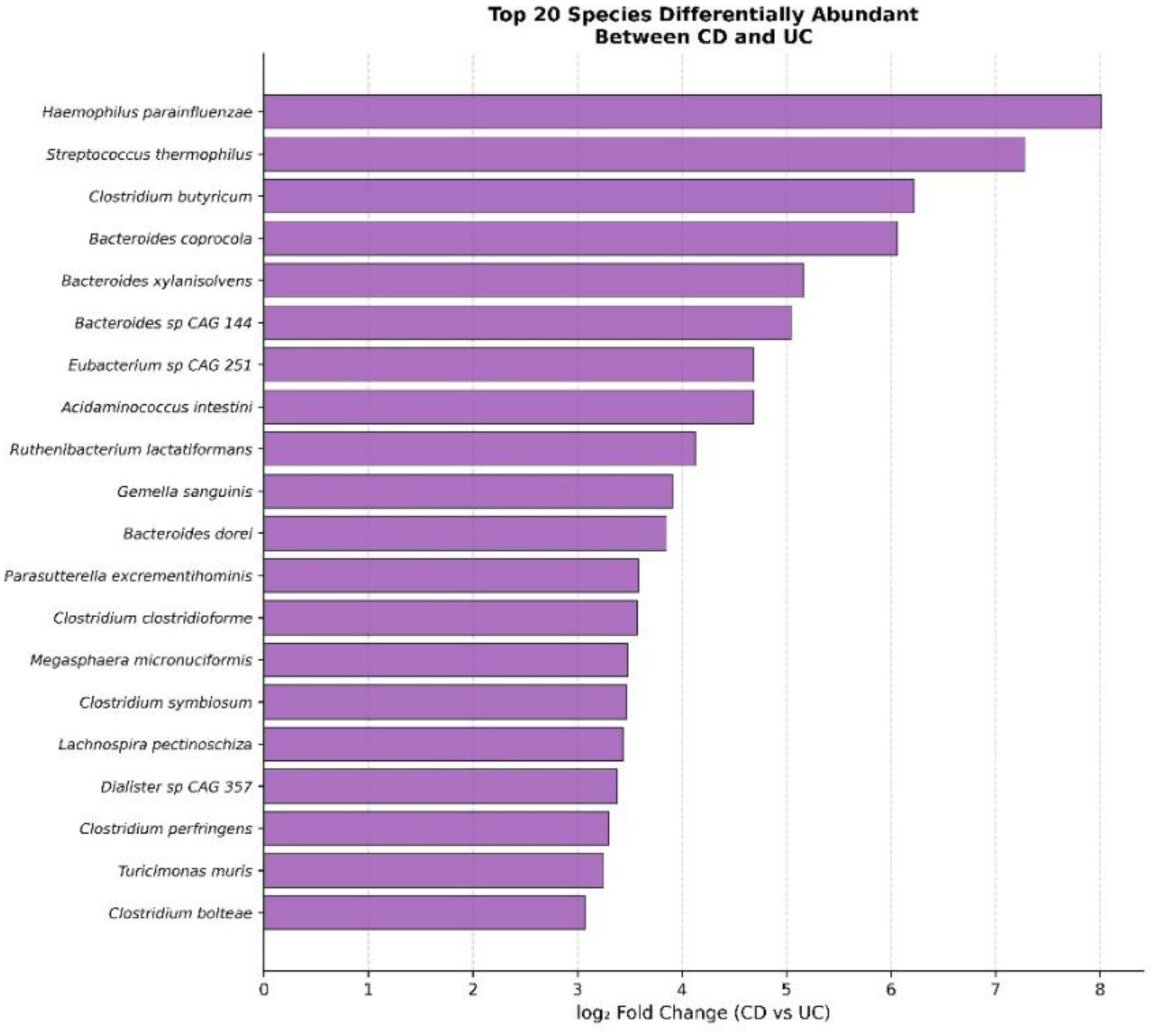
— CD vs UC direct species comparison: CD enriched.

**Figure 8G.**
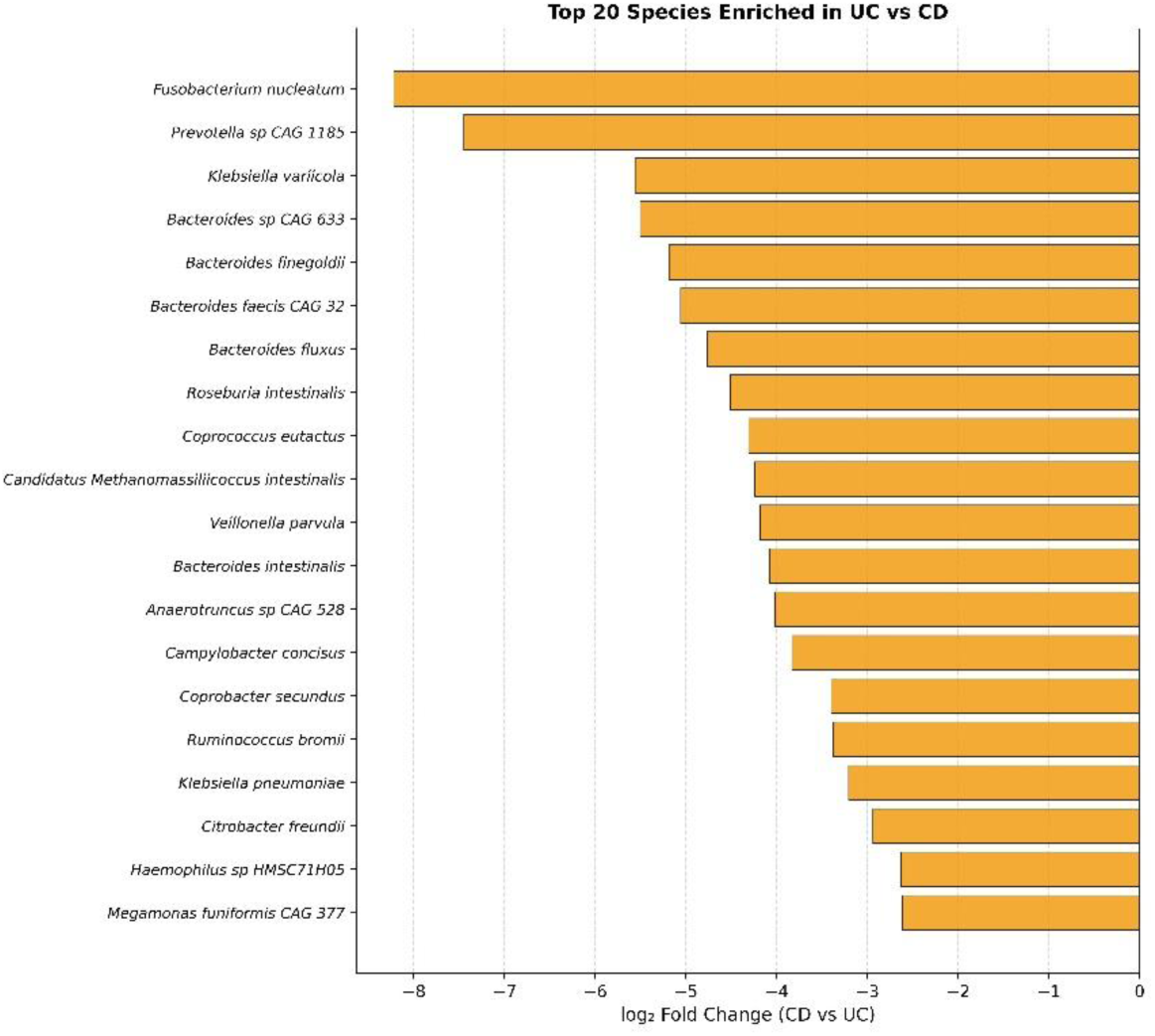
— CD vs UC direct species comparison: UC enriched.

### 3.8 Healthy gut microbiome carries out 577 exclusive biological processes

Direct GO analysis of the URIM output reveals findings which showcase a comprehensive molecular portrait of what a healthy human gut microbiome is doing for us, and what is catastrophically dismantled when IBD takes hold.

577 biological processes are exclusively present in the healthy microbiome and completely absent from both CD and UC **(Supplementary File 1 Sheet 1**). These were found to be not depleted, not reduced, but gone, that is, zero in both disease states. Thus, the healthy gut microbiome is not just a passive collection of bacteria. It is actively mimicking — and therefore potentially supporting — a vast array of host biological processes. When dysbiosis strikes, this entire layer of microbial contribution vanishes simultaneously in both forms of IBD. The non-IBD-exclusive processes read like a blueprint for gut homeostasis. The most important ones have been discussed below.

#### 3.8.1 Immune regulation

The exclusive presence of the term ‘**Antimicrobial humoral immune response mediated by antimicrobial peptide**’ in the healthy cohort indicates that healthy bacteria mimic AMP-response pathways. They are participating in the host’s own antimicrobial defence. This is gone in IBD. Figure 4B shows the depletion of the same in CD. Depletion of the same in UC can be studied in supplementary table **(Supplementary File 1 Sheet 1**), as the GO plots shown previously include only the top 20 terms. Additionally, the presence of the term ‘Cellular response to interferon-beta’ in only healthy individuals indicates that the healthy microbiome carries proteins mimicking antiviral interferon signalling, which are lost in IBD.

#### 3.8.2 Autophagy and protein quality control

The enrichment of the terms ‘Macroautophagy’, ‘reticulophagy’, and ‘intracellular protein localization’ indicate that the healthy microbiome mimics the host’s own cellular housekeeping machinery. These are completely absent in IBD. The gut loses its microbial contribution to protein turnover and organelle quality control simultaneously with disease onset.

#### 3.8.3 Neuroenteric homeostasis

The exclusive enrichment of terms ‘Norepinephrine metabolic process’ and ‘Gamma-aminobutyric acid (GABA) import’ in the Healthy cohort indicates that healthy microbes mimic norepinephrine metabolism. This is gone in IBD. GABA is the primary inhibitory neurotransmitter of the enteric nervous system **[Rao & Gershon, 2018].** Healthy microbiota mimic GABA import machinery. This is again entirely absent in both CD and UC.

#### 3.8.4 Digestive and metabolic

The terms ‘Digestive tract development’, ‘triglyceride metabolic process’, ‘response to hormone’, and ‘response to vitamin D’ were found to be healthy-exclusive. The microbiome is participating in the host’s own developmental and metabolic sensing. IBD erases this.

### 3.9 What CD gains that nobody else has (327 CD-exclusive BP terms)

Direct GO analysis of the URIM output reveals that CD bacteria are not just opportunistic **(Supplementary File 1 Sheet 2)**. They are acquiring entirely new functional capabilities: (i) Mitochondrial electron transport, cytochrome c to oxygen — CD-enriched bacteria mimic the terminal step of oxidative phosphorylation. This is striking. In the oxygen-rich, inflamed CD gut, bacteria that can mimic mitochondrial electron transport machinery have a massive survival advantage — and may actively interfere with host mitochondrial signalling; (ii) Ethanol catabolic process — CD bacteria can process ethanol. This connects to barrier disruption and mucosal permeability; (iii) Intrinsic apoptotic signalling pathway by p53 class mediator — CD bacteria mimic p53-mediated apoptosis regulators. Bacteria that can interfere with host cell death decisions have profound implications for epithelial turnover in Crohn’s transmural inflammation. (iv) Neuron cellular homeostasis — CD-exclusive; (v) Bile acid secretion — CD bacteria mimic bile acid secretion pathways. The ileum is the primary site of bile acid reabsorption and also the most common site of CD **[Baumgart & Sandborn, 2012];** (vi) Regulation of appetite — this is extraordinary. CD bacteria are mimicking host appetite regulation pathways. This may partially explain the profound anorexia and weight loss characteristic of Crohn’s disease; (vii) Positive regulation of nitric oxide biosynthetic process — NO is both a signalling molecule and an antimicrobial. CD bacteria mimicking NO production pathways could be subverting host innate defence; (viii) NOD2 signalling pathway — CD bacteria mimic NOD2 signalling. NOD2 is the most strongly associated genetic risk locus for Crohn’s disease **[Hugot et al., 2001].** The fact that CD-enriched bacteria mimic the very pathway whose mutation predisposes to CD is not coincidental. This is mechanistically explosive; (ix) Symbiont-mediated perturbation of host process — present only in CD. CD bacteria encode proteins that mimic the process of microbial host perturbation itself. They have evolved to interfere.

### 3.10 What UC gains that nobody else has (239 UC-exclusive BP terms)

Direct GO analysis of the URIM output reveals that UC is associated with immune hyperactivation and mucosal inflammatory signalling: (i) Cytokine-mediated signalling pathway (term_size=224) — the largest and most significant UC-exclusive term **(Supplementary File 1 Sheet 3)**. UC bacteria massively mimic host cytokine signalling. This is how UC maintains chronic mucosal inflammation — through microbial proteins that look like cytokine pathway components to the host immune system; (ii) T cell receptor signalling pathway (term_size=167) — UC bacteria mimic T cell receptor signalling. This connects directly to the aberrant T cell activation that drives UC pathology **[Kałużna et al., 2022];** (iii) Lipopolysaccharide-mediated signalling pathway — UC bacteria, despite being Gram-negative LPS producers themselves, also mimic LPS sensing pathways. They are simultaneously the stimulus and the receptor mimic; (iv) Histamine secretion and histamine biosynthetic process — UC-exclusive. UC bacteria mimic histamine production pathways. Histamine is a key mediator of mucosal oedema, mast cell activation, and visceral hypersensitivity. UC patients notoriously have elevated mucosal histamine **[Barcik et al., 2020];** (v) Neutrophil mediated immunity — UC bacteria mimic neutrophil immune functions. This aligns with UC’s hallmark neutrophilic mucosal infiltration; (vi) Dendritic cell antigen processing and presentation — UC bacteria mimic dendritic cell antigen presentation machinery. This is molecular mimicry at the level of antigen presentation itself; (vi) Germinal centre B cell differentiation — UC bacteria mimic B cell maturation pathways. UC is associated with mucosal IgG production and anti-neutrophil cytoplasmic antibodies (ANCA) **[Joossens et al., 2011].** (vii) Morphogenesis of an epithelium — UC bacteria mimic epithelial morphogenesis. UC causes profound crypt architectural distortion, and bacteria mimicking epithelial structural programmes could contribute to this; (viii) Response to amphetamine — this is unexpected and worth noting carefully. The dopaminergic system is involved in gut motility and visceral pain **[Sun et al., 2025]**. UC bacteria mimicking amphetamine response pathways suggests interference with dopaminergic neurotransmission in the gut.

### 3.11. Neuronal, signaling and SRP terms

#### 3.11.1 Neuronal and signaling terms

**Supplementary File 2** contains the details of all the Neuronal related GO terms and their log2fc

##### 3.11.1.1 Neural and Neuronal terms

The dominant pattern is severe depletion in both IBD subtypes — neural crest cell migration, regulation of synaptic assembly at neuromuscular junction, perineurial glial growth are all lost in both CD and UC (log2fc ≈ −8.33). Most striking divergence: neuronal action potential is strongly CD-specific (+8.33 CD, 0 UC), while neuron migration and regulation of neuron apoptotic process are UC-depleted but CD-neutral. Neuronal stem cell division is the only term enriched in both IBD subtypes.

The finding that neuronal stem cell division (GO:0036445) is the sole neuronal GO term enriched in both CD and UC relative to the nonIBD microbiome warrants careful consideration. Unlike the subtype-divergent neuronal signatures described above, this pan-IBD enrichment indicates a conserved feature of IBD-associated dysbiosis that transcends the mechanistic differences between Crohn’s disease and ulcerative colitis. Enteric neural progenitor cells — the enteric nervous system’s resident stem cell population — have been documented in the adult human gut and are increasingly recognised as contributors to enteric nervous system plasticity and repair **[Metzger et al., 2009; Azan et al., 2011].** Their division is regulated by kinases, cytoskeletal remodelling proteins, and cell fate determinants that are structurally conserved across eukaryotes and, at the sequence level, share homology with bacterial regulatory proteins **[Shih and Rothfield, 2006 ; Manuse et al., 2016].** The enrichment of GO:0036445 in both IBD microbiomes therefore suggests that bacteria enriched in the IBD gut — irrespective of whether the disease manifests as CD or UC — carry proteins structurally resembling the molecular machinery that governs enteric neural progenitor division. Whether such mimicry influences enteric neurogenesis or neural progenitor fate decisions in vivo remains to be established experimentally; however, the finding is consistent with emerging evidence that the gut microbiome actively modulates enteric nervous system plasticity [**Cryan et al., 2019]**, and raises the possibility that IBD-associated dysbiosis may impair the regenerative capacity of the enteric nervous system through molecular interference with neural stem cell division pathways. Notably, the simultaneous loss of broad neuronal projection and differentiation mimicry in both IBD subtypes — occurring alongside this conserved gain — suggests a shift from mimicry of mature neuronal function toward mimicry of progenitor-level cell division, potentially reflecting a dysbiosis-driven remodelling of the microbiome’s interface with the enteric nervous system from homeostatic support to proliferative pathway interference.

##### 3.11.1.2 Action potential

Every action potential term passes the log2fc threshold of 1, confirming this is a highly differential category. The CD-specific signature is unmistakable — neuronal action potential, membrane depolarization, cardiac muscle action potential, and voltage-gated sodium channel activity are all CD-exclusive (+8.33 CD, 0 UC). Critically, regulation of neuronal action potential is depleted in UC (−9.33) while enriched in CD — a clean, sharp divergence.

##### 3.11.1.3 Synapse

UC-specific enrichment dominates in synapse assembly, synaptic vesicle exocytosis, acetylcholine catabolic process in synaptic cleft, negative regulation of synaptic transmission, regulation of synaptic activity, postsynaptic modulation of chemical synaptic transmission with all these terms +8.66 in UC while zero in CD. CD-specific terms include regulation of synapse pruning, synaptic target inhibition, and regulation of postsynapse organization. Long-term synaptic potentiation shows the sharpest contrast of all: UC +1.91 vs CD −8.33. Meanwhile, glutamatergic synaptic transmission and synaptic vesicle endocytosis are depleted in both — lost from the healthy microbiome.

##### 3.11.1.4 Signaling

This was found to be the largest and most complex category. CD-specific enrichments include regulation of oxidative stress-induced neuron intrinsic apoptotic signaling pathway, intrinsic apoptotic signaling pathway by p53 class mediator (+9.33 CD only), TORC1 signaling, DNA damage response signaling, hippo signaling, and symbiont-mediated suppression of host toll-like receptor signaling — all CD-exclusive. UC-specific enrichments include cytokine-mediated signaling pathway (+9.66 UC), T cell receptor signaling pathway (+9.66 UC), lipopolysaccharide-mediated signaling pathway (+9.66 UC), acetylcholine receptor signaling pathway (+8.66 UC), and Toll signaling pathway — consistent with UC’s mucosal immune character. Integrated stress response signaling and positive regulation of calcium-mediated signaling are shared by both IBD types.

##### 3.11.1.5 Acetylcholine

Acetylcholine was found to be entirely UC-enriched. Acetylcholine catabolic process in synaptic cleft, acetylcholine binding, and acetylcholine receptor signaling pathway are all +8.66 in UC with zero CD signal. Acetylcholinesterase activity is mildly UC-enriched (+1.32). Cellular response to acetylcholine is depleted in UC (−1.26). Together this points to UC-specific disruption of the cholinergic anti-inflammatory reflex — bacteria in UC mimic the entire cholinergic machinery.

##### 3.11.1.6 Dendritic terms

Anterograde dendritic transport is the only shared enrichment (+8.33 CD, +8.66 UC) — the one IBD-conserved dendritic signal. UC shows enrichment of apical dendrite, distal dendrite, proximal dendrite compartments and dendritic cell antigen processing (+8.66 each). CD shows enrichment of dendrite cytoplasm (+9.33). Dendrite morphogenesis and dendrite membrane are lost in CD but neutral in UC — another clean subtype divergence.

Overall, the healthy microbiome maintains broad neuronal and synaptic mimicry; in IBD this is dismantled and replaced by subtype-specific signatures — CD acquires stress, apoptosis, and action potential mimicry while UC acquires cholinergic, synaptic vesicle, and immune signaling mimicry — and these two subtypes are as different from each other as either is from healthy.

#### 3.11.2 SRP terms

**Supplementary File 3** contains the details of all the SRP related GO terms and their log2fc

##### 3.11.2.1 Shared DOWN in both IBD (8 terms) — the core

This is the most important category. Protein targeting to membrane (GO:0006612, term_size=63) is severely depleted in both CD and UC (log2fc = −8.33 each) — this is the central SRP function, and it is lost from the IBD microbiome. The entire ER-associated machinery follows: ER-Golgi intermediate compartment membrane (−9.92 both), host cell ER membrane (−8.33 both), mitochondria-associated ER membrane contact site (−8.33 both), and ER stress-induced pre-emptive quality control (−8.33 both) are all lost. Small GTPase binding (−1.58 CD, −1.26 UC) and ceramide translocation (−9.33 both) round out this picture. Therefore, SRP-mediated protein targeting machinery mimicry is a feature of the healthy microbiome that is comprehensively dismantled in IBD regardless of subtype.

##### 3.11.2.2 Shared UP in both IBD (2 terms)

GTPase activator activity (+9.33 CD, +8.66 UC) and protein targeting to vacuole (+8.33 CD, +8.66 UC) are enriched in both IBD subtypes. GTPase activator activity is particularly interesting — bacteria in IBD appear to upregulate mimicry of GTPase regulation, which could reflect enhanced bacterial motility, vesicle trafficking manipulation, or interference with host Rab/Ras GTPase signaling.

##### 3.11.2.3 CD-specific UP (9 terms) — ER stress and secretory pathway hijacking

CD exclusively enriches ER chaperone complex (+8.33), cytosol to ER transport (+8.33), rough ER lumen (+8.33), perinuclear ER lumen (+8.33), cortical ER lumen (+8.33), protein targeting to peroxisome (+8.33), activation of GTPase activity (+8.33), positive regulation of phospholipid translocation (+9.33), and positive regulation of regulated secretory pathway (+8.33). This is striking — CD bacteria specifically acquire mimicry of the ER protein folding and secretory machinery, while simultaneously losing the core SRP targeting function. This could represent a bacterial strategy to interfere with host ER-dependent immune signaling specifically in the transmural inflammatory context of Crohn’s disease.

##### 3.11.2.4 UC-specific UP (2 terms)

ER unfolded protein response (UPR; +1.32 UC) and positive regulation of GTPase activity (+8.66 UC) are UC-exclusive. The UPR enrichment in UC is biologically coherent — UC involves chronic mucosal ER stress in goblet and epithelial cells, and UC-enriched bacteria appear to carry proteins mimicking the host UPR machinery, potentially exploiting or amplifying ER stress signaling in the colonic epithelium.

##### 3.11.2.5 CD-specific DOWN (5 terms)

ER-Golgi intermediate compartment (−1.00 CD), ER stress-induced intrinsic apoptotic signaling (−1.00 CD), and three obsolete mitochondrial targeting terms are specifically lost in CD. The ER stress-apoptosis depletion in CD is notable given that CD simultaneously gains ER chaperone mimicry — suggesting CD bacteria suppress apoptotic outcomes of ER stress while maintaining chaperone-like interactions.

##### 3.11.2.6 UC-specific DOWN (6 terms)

Translocation of peptides or proteins into host (−1.67 UC) is particularly meaningful — this term directly represents bacterial protein injection into host cells, and its depletion in UC suggests UC-enriched bacteria rely less on direct translocation and more on surface/secreted mimicry. Phospholipid translocation (−1.26 UC), positive regulation of protein targeting to membrane (−1.26 UC), and UC-specific ER membrane terms are also lost.

Overall, the healthy microbiome mimics the full SRP-mediated protein targeting axis; IBD dismantles this shared machinery and replaces it with subtype-specific ER manipulation — CD bacteria hijack the ER chaperone and secretory pathway while UC bacteria engage the unfolded protein response — both representing distinct bacterial strategies to interface with host ER stress signaling in the inflamed gut.

### 3.12 Single-ORF Sequence Mimicry: High-Resolution Traceback and Organismal Attribution

While the broad group-level sequence screening outlined in Section 3.6 provided an overview of human sequence homologs, its low average identity (34–35%) and lack of single-molecule resolution left open the possibilities of background alignment noise or minor host sequencing contamination. To definitively overcome these limitations and anchor our macro-level GO-term and PFAM-domain observations to specific bacterial taxa, we executed a parallel, high-stringency bottom-up validation pipeline.

This deep, single-ORF sequence search initially yielded 50,161 total human-homologous mimicry hits mapping across 1,286 unique human proteins. Eukaryotic filtering removed 5,073 ORFs whose top SwissProt hit originated from non-bacterial organisms, yielding 45,088 bacterially-sourced mimicry ORFs mapping onto 1,179 unique human protein targets. Across these confirmed microbial sequences, the mean sequence identity reached 44.3% (±7.9% SD) with a robust median alignment length of 212 amino acids. These alignment statistics indicate that the observed signals do not arise from fleeting, superficial motifs, but represent deep, structurally meaningful cross-kingdom sequence homology capable of driving true biomimetic host-microbe interactions.

**Figure S1 of Supplementary File 5** shows the distribution of molecular mimicry events. **Figure S2 of Supplementary File 5** shows top 30 most frequently mimicked human proteins. **Figure S3 of Supplementary File 5** shows top 30 bacterial proteins mimicking the human proteins.

**Supplementary File 6** contains the enrichment analysis of all human proteins mimicked by bacteria across all 3 cohorts, while **Supplementary File 7** contains the annotations for the same. However, **Supplementary File 8** contains the enrichment analysis of human proteins mimicked by proteins from all sources (including bacteria) across all 3 cohorts, and **Supplementary File 9** contains the annotations for the same.

#### 3.12.1 Healthy-Enriched Sequence Homology is Maintained by Canonical Commensal Clusters

The single largest sequence class isolated by the pipeline comprised 553 human proteins targeted by 26,855 individual microbial ORFs overrepresented in healthy controls (59.6% of the verified mimicry catalog). This category showed high alignment quality (mean sequence identity: 44.6%, mean *E*-value: 5.15 × 10^−8^). Top sequence-validated human mimicry targets included the metabolic enzymes GBA3 (glucosylceramidase beta 3; log_2_ FC = 4.09 vs CD), AL1A3 (aldehyde dehydrogenase; log_2_ FC = 3.70), GNL1 (log_2_ FC = 3.46), and SIAE (sialate O-acetylesterase; log_2_ FC = 3.17) (**Figure S4 of Supplementary File 5**).

Taxon-resolved attribution revealed that this baseline protective mimicry is explicitly sustained by a core network of canonical gut mutualists (**Figure S5 of Supplementary File 5**). *Bacteroides thetaiotaomicron* contributed the largest number of healthy-enriched mimicry ORFs (*n* = 2,262), closely supported by *Bacillus subtilis* (*n* = 2,124), *Phocaeicola vulgatus* (*n* = 1,235), *Parabacteroides distasonis* (*n* = 1,103), and *Bacteroides fragilis* (*n* = 1,072). Prominent short-chain fatty acid (SCFA) producers, including *Agathobacter rectalis* and *Lachnoclostridium phytofermentans*, emerged as key structural contributors. This clear taxonomic clustering indicates that host-microbiome homeostasis relies on a baseline structural repertoire provided by protective mutualists, which is systemically lost during inflammatory disruption.

#### 3.12.2 Crohn’s Disease Shifts the Mimicry Landscape toward Active Pathobiont Profiles

In Crohn’s Disease, the pipeline exposed an entirely rearranged sequence landscape: 349 human proteins were selectively targeted by 17,424 CD-specific ORFs (38.6% of the global catalog; mean sequence identity: 44.0%, mean *E*-value: 5.73 × 10^−8^). The most aggressively enriched human targets under this disease state included oxidative stress and structural maintenance factors: MSRB3 (methionine sulfoxide reductase B3; log_2_ FC = 3.70 vs nonIBD), NQO2 (log_2_ FC = 3.17), TOP2B (log_2_ FC = 3.17), TREA (log_2_FC = 3.17), and HEXA (log_2_ FC = 3.17) (**Figure S6 of Supplementary File 5**).

When we parsed the taxonomic origins of these CD-associated sequence signatures, the donor profile diverged sharply from health (**Figure S7 of Supplementary File 5**). While the resilient generalist *Bacteroides thetaiotaomicron* retained a high footprint (*n* = 2,398), the remaining space was dominated by classic inflammatory pathobionts. *Bacteroides fragilis* expanded its structural sequence contribution (*n* = 1,294), accompanied by prominent appearances of *Escherichia coli O157:H7* (*n* = 563) and *Haemophilus influenzae* (*n* = 294).

Furthermore, oral-translocated species like *Porphyromonas gingivalis* surfaced as major contributors to this CD-enriched mimicry ORFs. This direct sequence mapping confirms our global taxonomic profiling data by reinforcing the findings of Section 3.7, establishing that the expanded mimicry of human tissue machinery in Crohn’s Disease is explicitly driven by an invading consortium of Gram-negative pathobionts and those that escaped from oral-cavity (oral-origin species consistent with the oral-gut translocation hypothesis).

#### 3.12.3 Ulcerative Colitis Displays a Sparse, Fragmented Sequence-Mimicry Profile

The sequence-level landscape in Ulcerative Colitis was in stark contrast to Crohn’s Disease, accounting for just 54 unique human proteins targeted by a minor pool of 297 ORFs (0.66% of the validated dataset). These alignments displayed reduced structural affinity (mean identity: 41.4%, mean *E*-value: 3.37 × 10^−7^). Top UC-enriched targets included MRP2 (log_2_ FC = 3.17 vs nonIBD), CTR1 (log_2_ FC = 2.81), and HELLS (log_2_FC = 2.32) (**Figure S8 of Supplementary File 5**).

Mimicry ORFs were distributed across unrelated, environmentally derived taxa—such as *Rhodopseudomonas palustris*, *Methanocaldococcus jannaschii*, and *Aquifex aeolicus*—rather than localized within a disease-associated gut community (**Figure S9 of Supplementary File 5**). This sparse signature mirrors the minimal functional divergence observed in our GO-term and PFAM-level analyses, confirming that unlike the extensive functional restructuring seen in CD, the UC microbiome preserves a sequence-level mimicry profile more similar to healthy controls than to CD.

**Figure S10 of Supplementary File 5** shows top 30 human proteins commonly mimicked by ORFs from both CD and UC gut.

#### 3.12.4 Conserved Housekeeping Enzymes Form the Primary Structural Bridge Between Kingdoms

The single most pervasive mimicking bacterial sequence groups belonged to highly conserved metabolic and structural catalysts (**Figure S3 of Supplementary File 5**).

The absolute dominant drivers included:

- GALM_ACICA (aldose 1-epimerase from *Acinetobacter calcoaceticus*; *n* = 277 ORFs)
- YCHF_BACSU (ribosome-binding ATPase YchF from *Bacillus subtilis*; *n* = 263)
- RECQ_ECOLI (ATP-dependent DNA helicase RecQ from *Escherichia coli*; *n* = 216)
- ASPG1_ECO57 (L-asparaginase 1 from *E. coli O157:H7*; *n* = 212)
- MUTB_PORGI (methylmalonyl-CoA mutase large subunit from *Porphyromonas gingivalis*; *n* = 205)

The heavy concentration of these core mutases, epimerases, helicases, and ATPases highlights the structural conservation underpinning cross-kingdom mimicry. This direct sequence verification matches our macro-scale GO findings, confirming that ancient, highly conserved housekeeping enzymes provide primary basis for sequence-level molecular mimicry in the gut.

## 4. Discussion

### 4.1 Multi-layered Molecular Mimicry Framework

This study presents a systematic, metagenome-wide characterization of molecular mimicry across IBD subtypes using shotgun metagenomic data from the HMP2 IBDMDB cohort. By applying three complementary analytical frameworks — normalized GO term frequency analysis, PFAM domain enrichment, and sequence-level human homology detection — we capture molecular mimicry at multiple levels of biological granularity. This multi-level design is intentional: functional mimicry (shared GO term space) and sequence-level mimicry (direct amino acid similarity to human proteins) represent mechanistically distinct phenomena with different potential immunological consequences, and examining both provides a more complete picture of host-microbe functional relationships in IBD.

A consistent finding across all three analytical layers is that CD-associated microbiomes show substantially greater divergence from healthy controls than UC. This convergence of signals from independent analyses — more differentially represented GO terms, more enriched PFAM domains, and a higher per-sample sequence mimicry rate in CD — strengthens confidence in the biological interpretation and is not attributable to a single analytical choice.

### 4.2 CD Shows Greater Functional Divergence Than UC

The greater functional divergence of CD from healthy controls, observed consistently across GO, PFAM, and sequence-level analyses, is biologically coherent with the known pathophysiology of the two IBD subtypes. CD is a transmural, discontinuous inflammatory condition affecting any gastrointestinal segment, while UC is confined to the colonic mucosa. The deeper structural disruption and broader anatomical involvement of CD likely creates conditions permissive for colonization by a more diverse range of pathobionts and functionally distinct microbial communities.

This interpretation is supported by the taxonomic data, which revealed more pronounced species-level dysbiosis in CD, including the enrichment of oral-origin bacteria (*Haemophilus parainfluenzae, Streptococcus thermophilus*) and known pathobionts (*Bacteroides fragilis, Clostridium perfringens, Klebsiella michiganensis*) not seen at comparable levels in UC. The enrichment of oral bacteria in particular is consistent with the oral-gut microbial translocation hypothesis, which proposes that breakdown of upper gastrointestinal barrier function in CD allows oral commensals to establish in the lower gut, where they may drive inflammatory responses through multiple mechanisms including molecular mimicry **[Sohn et al., 2023]**.

### 4.3 Biological Significance of GO Term Findings

Among the GO terms most enriched in CD microbiomes, the top hit — adaptive immune response — is directly and mechanistically relevant to IBD pathogenesis. The enrichment of microbial proteins with functional annotations related to adaptive immunity processes raises the possibility that gut bacteria in CD harbor proteins capable of engaging or modulating host adaptive immune pathways. This could occur through direct molecular mimicry of immune signaling components, through shared enzymatic activities that modify immunologically active substrates, or through structural domains recognized by pattern recognition receptors.

The enrichment of bone remodeling as a GO term in CD microbiomes is also noteworthy, as osteopenia and osteoporosis are recognized extraintestinal manifestations of CD, occurring in a significant proportion of patients **[Baban et al., 2021]**. While causality cannot be established from these correlative data, the functional representation of bone remodeling-related proteins in the CD microbial proteome suggests a possible mechanistic link between microbial functional mimicry and skeletal complications of CD. Similarly, the enrichment of cellular response to misfolded protein — an endoplasmic reticulum stress pathway — is consistent with the known role of ER stress in Paneth cell dysfunction in CD, a process increasingly recognized as central to CD pathogenesis **[Adolph et al., 2013]**.

The UC-specific enrichment of cytokine-mediated signaling pathway and T cell receptor signaling pathway terms is consistent with the T cell-driven mucosal immune activation characteristic of UC, though the modest number of differentially represented terms in UC overall tempers strong conclusions.

### 4.4 PFAM Domain Enrichment: Structural Underpinning

The differential PFAM domain profiles between diagnostic groups provide structural context for the GO term findings. The enrichment of SMC family RAD50 subfamily proteins in CD is of particular interest: RAD50 is a eukaryotic DNA repair protein, and bacterial proteins with RAD50-like domains have been described but are not ubiquitous. Their enrichment in CD microbiomes suggests expansion of taxa harbouring these eukaryote-like structural architectures, which could facilitate direct protein-protein interactions with host DNA damage response components. The enrichment of Major facilitator superfamily oligosaccharide transporter family proteins in both CD and UC points to altered carbohydrate transport capabilities in IBD microbiomes, consistent with the known role of microbial carbohydrate metabolism in shaping the intestinal environment. The depletion of RTX prokaryotic toxin family proteins in both IBD groups relative to nonIBD controls is of uncertain biological significance. RTX toxins are produced predominantly by oral-origin pathobionts including *Actinobacillus*, *Aggregatibacter*, and *Kingella* species, which are not major constituents of the healthy gut microbiome [**Linhartová et al., 2010].** Their detection in the nonIBD metagenome likely reflects transient oral microbial translocation to the gut, and their depletion in IBD may reflect altered mucosal barrier dynamics or compositional shifts in oral-origin taxa rather than loss of a protective commensal function. Taxon-resolved analysis would be required to interpret this finding definitively.

### 4.5 Sequence-level Mimicry: Implications and Limitations

The identification of 219–238 unique human proteins matched by microbial sequences across diagnostic groups represents a set of candidate molecular mimicry targets. The preferential targeting of human carbohydrate-processing enzymes — particularly lysosomal glycosidases (NAGLU, HGSNAT, FUCA1, FUCA2) and sialic acid metabolism enzymes (SIAE) — is biologically plausible given that gut bacteria extensively utilize host glycans as carbon sources and have evolved carbohydrate-active enzymes with structural similarities to host enzymes through convergent evolution or horizontal gene transfer.

The elevated per-sample sequence mimicry rate in CD (2,250 vs 1,550 per 100,000 ORFs in nonIBD, upon normalization) suggests qualitative enrichment of human-mimicking protein families in the CD microbial community, beyond what is explained by total ORF differences. The mechanistic interpretation of this difference — whether it reflects expansion of specific human-mimicking taxa in CD, upregulation of horizontally transferred eukaryotic-like genes, or other processes — cannot be resolved from these data alone and requires experimental validation.

An important caveat is that sequence similarity at the 34–35% mean identity observed here represents distant homology, falling in the ‘twilight zone’ of sequence alignment where structural and functional relationships may not be conserved. Cross-reactive immune responses in this identity range are possible but cannot be assumed. Structural validation using AlphaFold2-predicted models and epitope prediction would be necessary to assess the immunological relevance of these mimicry candidates.

### 4.6 The Attenuated UC Signal

The consistently attenuated differential signals in UC across all analytical layers — fewer differentially represented GO terms, less extreme PFAM enrichment, lower per-sample sequence mimicry rate, and less severe taxonomic dysbiosis — is one of the most consistent findings of this study. This pattern is consistent with the clinical and pathological characteristics of UC, which typically presents with more predictable disease extent and course than CD, and with epidemiological evidence suggesting a stronger genetic component (higher concordance in twins) and potentially a more restricted set of environmental triggers in UC **[Jostins et al., 2012]**.

The paradoxical observation that UC has lower sequence mimicry rates than even healthy controls (1,338 vs 1,550 per 100,000 ORFs, upon normalization) warrants consideration. One explanation is the depletion in UC of specific commensal taxa that normally contribute human-sequence-mimicking proteins to the gut metagenome — taxa whose loss in UC may itself be part of the disease process. Another possibility is that the smaller mean ORF count in UC samples (similar to CD at ∼40k) does not fully explain the pattern, and that there is a genuine qualitative difference in the functional composition of UC microbiomes. Prospective studies with larger sample sizes and longitudinal sampling will be needed to clarify this observation.

### 4.7 Taxonomic Context and Known Biology

The taxonomic enrichment patterns observed here are generally consistent with prior reports from the HMP2 cohort and independent IBD microbiome studies. The strong enrichment of *Haemophilus parainfluenzae* in CD (log₂FC = 5.96) replicates a finding from the original HMP2 multi-omics study and multiple subsequent reports, lending confidence to the validity of this analysis **[Lloyd-Price et al., 2019].**

The depletion of *Akkermansia muciniphila* in CD (log₂FC = −5.65) is also well-established in the literature and consistent with the role of this keystone commensal in maintaining mucus layer integrity — a function whose loss in CD would be expected to increase microbial access to epithelial surfaces and potentially amplify mimicry-mediated immune interactions **[Zheng et al., 2023].**

The protective functional signatures identified in healthy microbiomes — including enrichment of Class-III pyridoxal-phosphate-dependent aminotransferase activities and specific transport families — point to metabolic functional capabilities that may be lost during IBD-associated dysbiosis. Whether these represent causes or consequences of the inflammatory state cannot be determined from cross-sectional baseline data, and longitudinal studies correlating functional mimicry signals with disease activity would be informative.

### 4.8 Human gut microbiome as a functional mirror of the host

The healthy human gut microbiome is not merely a collection of metabolic contributors. It is a functional mirror of the host — a vast library of microbial proteins that structurally resemble human biological machinery across immunity, autophagy, neuroenteric signalling, hormone response, and developmental regulation. This mimicry library is maintained in health and systematically dismantled in disease.

IBD is not simply characterised by what the microbiome loses. It is characterised by what it replaces the lost mimicry with — and CD and UC replace it with entirely different things, reflecting their distinct pathophysiologies. CD replaces homeostatic mimicry with survival and host-perturbation mimicry — mitochondrial interference, p53 apoptosis manipulation, NOD2 pathway mimicry, appetite dysregulation, bile acid pathway capture. UC replaces homeostatic mimicry with immune amplification mimicry — cytokine signalling, T cell receptor pathways, LPS sensing, histamine production, antigen presentation, B cell maturation. This is a molecular explanation for why CD and UC are observed as different diseases despite both being IBD. They are driven by microbiomes that have diverged not just in composition, but in their entire functional mimicry repertoire.

Thus, the healthy gut microbiome maintains a broad functional mimicry repertoire spanning immunity, autophagy, and neuroenteric signalling, which is systematically dismantled in IBD and replaced by disease-subtype-specific mimicry signatures that reflect the distinct pathophysiologies of Crohn’s disease and ulcerative colitis.

### 4.9 NOD2 mimicry in CD

Nucleotide-binding oligomerization domain-containing protein 2 (NOD2), also known as caspase recruitment domain-containing protein 15 or inflammatory bowel disease protein 1, is a protein that in humans is encoded by the NOD2 gene located on chromosome 16. NOD2 is the primary genetic risk locus for Crohn’s disease or CD. It plays an important role in the immune system. As per our findings, bacteria enriched in CD were found mimicking NOD2, and the subsections below discuss in detail the implications of the same.

#### 4.9.1 NOD2 as an intracellular pattern recognition receptor

NOD2 is an intracellular pattern recognition receptor. Its job is to detect bacterial cell wall components — specifically muramyl dipeptide (MDP), a fragment of peptidoglycan found in both Gram-positive and Gram-negative bacteria — and trigger an innate immune response **[Girardin et al., 2003].** It is the host’s internal bacterial sensor.

Mutations in NOD2 are the single strongest genetic risk factor for Crohn’s disease **[Hugot et al., 2001].** Patients with loss-of-function NOD2 mutations cannot properly sense and respond to bacterial invasion of the intestinal epithelium. The gut becomes permissive to bacterial overgrowth and chronic inflammation follows.

This has been known for over 20 years. The question the field has never been able to fully answer is: why does NOD2 dysfunction lead specifically to Crohn’s disease and not some other inflammatory condition?

#### 4.9.2 Findings from our study and its implications

CD-enriched bacteria — bacteria that are specifically and exclusively present in Crohn’s disease patients and absent from both healthy individuals and UC patients — carry proteins that structurally mimic the NOD2 signalling pathway. Nucleotide-binding oligomerization domain containing 2 signalling pathway is a CD-exclusive GO term in our data. Zero in nonIBD. Zero in UC. Present only in CD.

What we thus propose is: (i) NOD2 normally detects bacteria and triggers immune clearance. In many CD patients, NOD2 is mutated and non-functional — bacteria are not properly detected or cleared. (ii) Into this environment of impaired NOD2 surveillance, CD-specific bacteria expand. (iii) These CD-specific bacteria have evolved proteins that mimic the NOD2 signalling pathway itself. (iv) CD Bacteria mimic the very pathway designed to detect and destroy them. Because if a bacterium carries a protein that looks like a NOD2 pathway component, it can potentially: (a) Interfere with or saturate NOD2 signalling — acting as a decoy, (b) Suppress downstream NF-κB activation that would otherwise clear them, (c) Exploit the already-impaired NOD2 system in genetically susceptible CD patients to establish persistent colonisation, and (d) Trigger aberrant NOD2 signalling in patients with functional NOD2, contributing to chronic inflammation. The same can be visually understood from **Figure 9**.

**Figure 9.**
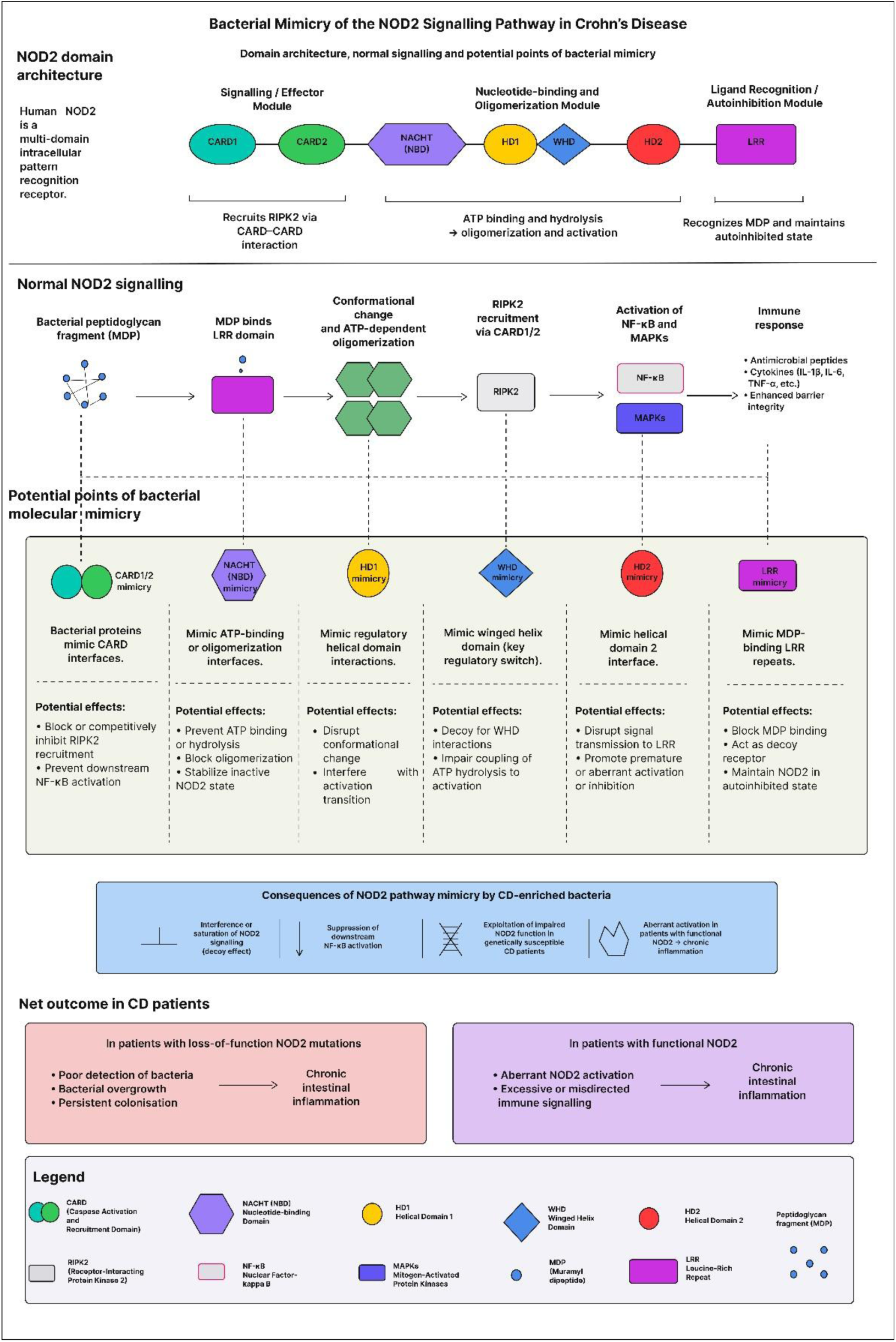
Bacterial mimicry of the NOD2 signalling pathway in Crohn’s disease: domain architecture, normal signalling, and potential points of interference.

#### 4.9.3 The deeper implication

Bacterial Mimicry of NOD2 Signalling Pathway and its impairment creates a vicious cycle that has never been described before at the molecular mimicry level (**Figure 10**).

**Figure 10.**
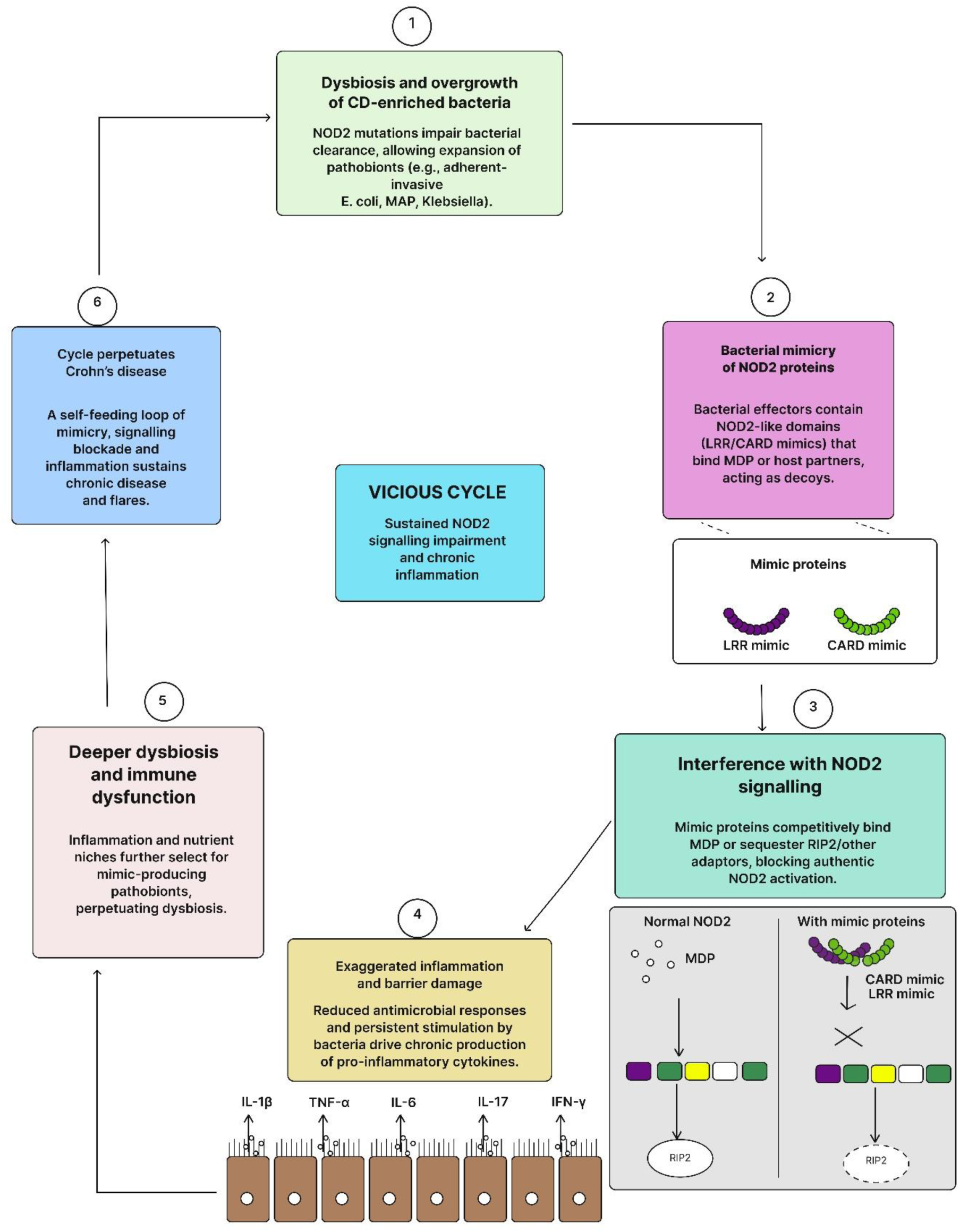
The NOD2 vicious cycle: molecular mimicry as a mechanistic bridge between genetic susceptibility and microbial dysbiosis in Crohn’s disease.

The genetic risk factor and the microbial environment are not independent. The bacteria that thrive in a NOD2-deficient gut have evolved to exploit and amplify exactly that deficiency — through molecular mimicry of the NOD2 pathway itself.

For 20 years, the IBD genetics community and the IBD microbiome community have worked largely in parallel. Genome-wide association studies identified NOD2. Metagenomics identified dysbiosis. The two fields have known they are connected but the molecular mechanism linking them has been elusive.

Our data provides a candidate mechanism: CD-enriched bacteria mimic the NOD2 pathway, creating a molecular bridge between the genetic architecture of CD susceptibility and the functional behaviour of the dysbiotic microbiome. This finding thus reframes how the disease is understood.

#### 4.9.4 Limitations and need for further validation

This is computational — sequence-level mimicry identified through GO term mapping. We have not shown that these bacterial proteins functionally interfere with NOD2 signalling *in vitro* or *in vivo*. That validation is a much necessary future work. Therefore, our study can be understood as a hypothesis-generating computational finding.

### 4.10 The Microbial Signal Recognition and Neuronal Mimicry (SRNM) Axis in IBD

The integrated findings from neuronal, signalling, and signal recognition particle (SRP)-associated GO term analyses reveal a coherent and previously unrecognised mechanistic axis operating at the interface of the gut microbiome and the host neuroimmune system. We designate this the Microbial Signal Recognition and Neuronal Mimicry (SRNM) axis — a dual-component functional framework in which the healthy gut microbiome simultaneously maintains mimicry of host neuronal regulatory machinery and SRP-mediated protein targeting pathways, and in which both components are coordinately dismantled in IBD and replaced by disease-subtype-specific compensatory mimicry signatures. The SRNM axis thus represents not a single pathway but a systems-level property of the gut microbiome — one that distinguishes health from disease, and Crohn’s disease from ulcerative colitis, at the level of microbial functional mimicry.

#### 4.10.1 The healthy gut microbiome as a neuronal mimicry maintainer

A foundational and perhaps counterintuitive finding of the present study is that broad neuronal mimicry is a feature of the healthy gut microbiome, not of the diseased one. GO terms encompassing neuron projection development (GO:0031175), neurogenesis (GO:0022008), neuron differentiation (GO:0030182), and projection morphogenesis (GO:0048812) are present in the nonIBD microbiome and depleted or absent in both CD and UC. The depletion of neuron projection morphogenesis is among the most severe observed in the entire dataset, with log₂fc values of approximately −8.33 in both CD and UC, and neuron development (GO:0048666) is depleted to log₂fc −9.92 in UC. These values indicate near-complete absence of microbial proteins mimicking these pathways in the IBD gut.

This pattern is consistent with emerging evidence that commensal microbiota actively support enteric nervous system homeostasis through direct molecular interactions with host neuronal tissue **[Obata & Pachnis, 2016].** The enteric nervous system contains approximately 500 million neurons and operates semi-autonomously, regulating intestinal motility, secretion, and immune surveillance **[Rao & Gershon, 2018].** The finding that healthy microbiota carry proteins structurally resembling host neuronal projection and differentiation machinery suggests a previously unrecognised mode of microbiome-nervous system crosstalk — one that operates not through metabolite production or immune modulation but through functional molecular mimicry at the protein sequence level. The loss of this mimicry in IBD implicates dysbiosis not merely as a consequence of altered microbial composition, but as a direct contributor to the neurological and visceral sensory disturbances that are increasingly recognised as core features of IBD pathophysiology **[Sun et al., 2025]**.

#### 4.10.2 CD-specific neuronal mimicry: stress, apoptosis, and action potential signatures

While broad neuronal mimicry is lost in IBD, the CD microbiome acquires a distinct and functionally coherent set of neuronal mimicry signals that are entirely absent in both UC and nonIBD individuals.

Neuron cellular homeostasis (GO:0070050) is the most strongly enriched neuronal term in CD (log₂fc = +9.33), and neuronal action potential (GO:0019228), regulation of synapse pruning (GO:1905806), modification of postsynaptic structure (GO:0099010), and regulation of oxidative stress-induced neuron intrinsic apoptotic signalling (GO:1903376) are all CD-exclusive, enriched at log₂fc ≈ +8.33 with zero signal in UC. The action potential signature is particularly striking: every action potential-related GO term in the dataset passes the log₂fc ≥ 1 threshold exclusively in CD, including membrane depolarisation, cardiac muscle action potential, and voltage-gated sodium channel activity. Critically, regulation of neuronal action potential is depleted in UC (log₂fc = −9.33) while enriched in CD — the sharpest single directional divergence between the two disease subtypes observed in the neuronal category.

These findings point to a model in which CD-enriched bacteria have evolved proteins mimicking host machinery involved in neuronal stress responses, apoptotic fate decisions, and electrochemical signalling. This is mechanistically coherent with the biology of Crohn’s disease: CD is characterised by transmural, discontinuous inflammation with pronounced oxidative burden and deep tissue involvement, creating an environment in which bacteria that can interface with host neuronal stress pathways would have a significant survival and colonisation advantage **[Alemany-Cosme et al., 2021].** The enrichment of oxidative stress-induced neuronal apoptosis mimicry in CD bacteria aligns with the documented degeneration of enteric neurons in Crohn’s disease, which has been attributed to oxidative stress and apoptotic signalling **[Jamka & Gulbransen, 2025]**. The possibility that CD-enriched bacteria carry proteins that modulate these very pathways through molecular mimicry warrants direct experimental investigation.

The synapse-level divergence between CD and UC is equally informative. Regulation of synapse pruning and synaptic target inhibition are CD-specific gains, whereas long-term synaptic potentiation (GO:0060291) shows a CD vs. UC log₂fc contrast of −16.66 — enriched in UC at +1.91 and depleted in CD at −8.33. Glutamatergic synaptic transmission and synaptic vesicle endocytosis are depleted in both subtypes, indicating loss of healthy glutamatergic mimicry irrespective of disease type. Together, these synaptic findings suggest that CD microbiota preferentially target pruning and inhibitory synaptic regulation, whereas UC microbiota target excitatory synaptic transmission and potentiation — a divergence that maps strikingly onto the known differences in visceral pain and enteric neuropathy between the two diseases **[Morales-Soto et al., 2023].**

#### 4.10.3 UC-specific neuronal mimicry: the cholinergic anti-inflammatory axis

The UC microbiome acquires a neuronal and neurotransmitter mimicry signature that is qualitatively distinct from CD and centred on the cholinergic system. Synapse assembly (GO:0007416), synaptic vesicle exocytosis (GO:0016079), synaptic vesicle fusion to presynaptic active zone membrane (GO:0031629), regulation of synaptic activity (GO:0060025), postsynaptic modulation of chemical synaptic transmission (GO:0099170), and negative regulation of synaptic transmission (GO:0050805) are all enriched at log₂fc ≈ +8.66 in UC and absent in CD. The acetylcholine subsystem is entirely UC-specific: acetylcholine catabolic process in the synaptic cleft (GO:0001507), acetylcholine binding, acetylcholine receptor signalling pathway, and acetylcholinesterase activity are exclusively enriched in UC. Neuroblast differentiation (GO:0014016) and motor neuron apoptotic process (GO:0097049) are likewise UC-exclusive.

This UC-specific cholinergic signature has clear pathophysiological relevance. The cholinergic anti-inflammatory reflex — mediated by the vagus nerve and enteric cholinergic neurons — is a primary mechanism of mucosal immune suppression in the colon, and its impairment has been documented in UC **[Kanauchi et al., 2022].** Acetylcholine released by enteric neurons acts on α7 nicotinic acetylcholine receptors on macrophages to suppress TNF-α and other pro-inflammatory cytokines **[Wang et al., 2003]**. The finding that UC-enriched bacteria carry proteins mimicking the entire cholinergic synaptic machinery — from vesicle exocytosis to acetylcholinesterase activity to receptor signalling — raises the possibility that these bacteria interfere with or exploit the cholinergic anti-inflammatory reflex, potentially amplifying mucosal inflammation by subverting the neuronal brake on immune activation. The simultaneous depletion of cellular response to acetylcholine in UC (log₂fc = −1.26) alongside enrichment of acetylcholine biosynthetic and catabolic mimicry further supports a model of dysregulated cholinergic neurotransmission in the UC gut microbiome.

The dendritic compartment adds a further dimension to the UC-specific signature. Apical dendrite, distal dendrite, proximal dendrite, and dendritic cell antigen processing and presentation are all UC-enriched (log₂fc ≈ +8.66), while dendrite morphogenesis and dendrite membrane are depleted in CD but neutral in UC. Anterograde dendritic transport (GO:0098937) and dendritic mRNP transport (GO:0098963) represent the only neuronal terms enriched in both IBD subtypes (log₂fc +8.33 CD, +8.66 UC), suggesting that disruption of dendritic transport machinery is a conserved feature of IBD-associated microbial mimicry, irrespective of disease subtype.

#### 4.10.4 The signalling landscape: shared and divergent IBD signatures

Beyond neuronal-specific terms, the broader signalling category reveals the full breadth of the CD vs. UC divergence. CD-exclusive signalling enrichments cluster around intracellular stress and survival pathways: intrinsic apoptotic signalling pathway by p53 class mediator (GO:0097193; log₂fc = +9.33), TORC1 signalling, DNA damage response signalling, Hippo signalling, and — notably — symbiont-mediated suppression of host toll-like receptor signalling are all CD-specific. The suppression of host TLR signalling is mechanistically significant: TLR pathways are primary sentinels of bacterial detection at the intestinal epithelium, and CD-enriched bacteria that mimic host TLR suppression machinery may thereby evade innate immune detection, facilitating persistent colonisation in the transmural environment characteristic of Crohn’s disease **[J Worley, 2023]**.

The UC-specific signalling landscape is dominated by immune amplification pathways. Cytokine-mediated signalling pathway (GO:0019221; term_size = 224; log₂fc = +9.66), T cell receptor signalling pathway (GO:0050852; term_size = 167; log₂fc = +9.66), and lipopolysaccharide-mediated signalling pathway (GO:0031663; log₂fc = +9.66) are all UC-exclusive. The acetylcholine receptor signalling pathway and Toll signalling pathway are further UC-specific enrichments. That UC-enriched bacteria simultaneously mimic LPS sensing pathways — the very immune surveillance system designed to detect them — and T cell receptor signalling pathways suggests a sophisticated strategy of immune pathway infiltration, consistent with the adaptive immune-dominant pathophysiology of ulcerative colitis **[Kałużna et al., 2022]**. Integrated stress response signalling and positive regulation of calcium-mediated signalling are shared between both IBD subtypes, representing pan-IBD features of the dysbiotic microbial signalling mimicry repertoire.

#### 4.10.5 The SRP component of the SRNM axis: coordinated loss and subtype-specific ER manipulation

The signal recognition particle arm of the SRNM axis mirrors the neuronal arm in its overall architecture: a set of mimicry functions present in health is coordinately lost in both IBD subtypes, and each subtype acquires distinct compensatory mimicry of adjacent pathways. Core SRP-mediated protein targeting to membrane (GO:0006612; term_size = 63) is severely depleted in both CD and UC (log₂fc = −8.33 each). The entire ER-associated machinery that depends on SRP function follows the same pattern: ER-Golgi intermediate compartment membrane, host cell ER membrane, mitochondria-associated ER membrane contact site, and ER stress-induced pre-emptive quality control are all lost in both subtypes, with log₂fc values ranging from −8.33 to −9.92. Small GTPase binding (−1.58 CD, −1.26 UC) and ceramide translocation (−9.33 both) complete the picture of comprehensive SRP pathway depletion in IBD.

The SRP is a universally conserved ribonucleoprotein complex responsible for co-translational targeting of membrane and secreted proteins. In bacteria, SRP consists of the Ffh protein and 4.5S RNA, functionally analogous to eukaryotic SRP54 and SRP RNA **[Luirink & Sinning, 2004].** Elevated SRP function in commensal bacteria would facilitate correct membrane localisation of a wide range of host-interacting proteins, including those involved in mucosal adherence, metabolite exchange, and immune modulation. The loss of SRP mimicry in both IBD subtypes therefore implies a fundamental disruption of the membrane protein targeting machinery in dysbiotic bacteria, with downstream consequences for the full spectrum of host-microbe molecular interactions mediated through the bacterial membrane.

Rather than simply losing SRP function, however, CD and UC bacteria acquire distinct alternative strategies. CD exclusively enriches mimicry of the ER chaperone complex (GO:0034663; log₂fc = +8.33), cytosol-to-ER transport (GO:0046967), rough ER lumen, perinuclear ER lumen, cortical ER lumen, protein targeting to peroxisome, and positive regulation of the regulated secretory pathway and phospholipid translocation (log₂fc = +9.33). This CD-specific acquisition of ER chaperone and secretory pathway mimicry, occurring simultaneously with loss of core SRP targeting function, suggests a bacterial strategy of ER functional hijacking: CD bacteria may interfere with host ER-dependent immune signalling, antigen processing, or secretory protein production specifically in the context of the severely inflamed, transmural Crohn’s gut. The concurrent depletion of ER stress-induced intrinsic apoptotic signalling (log₂fc = −1.00) in CD — while ER chaperone mimicry is gained — is consistent with a model in which CD bacteria suppress the apoptotic consequences of ER stress while maintaining chaperone-mediated interactions, thereby prolonging bacterial survival within the ER stress milieu of the inflamed epithelium **[Miao et al., 2025].**

UC bacteria acquire a narrower but biologically coherent SRP-adjacent signature: ER unfolded protein response (GO:0030968; log₂fc = +1.32 UC) and positive regulation of GTPase activity (GO:0043547; log₂fc = +8.66 UC) are UC-exclusive. The UPR is chronically activated in the colonic epithelium of UC patients, particularly in goblet cells, where ER stress has been identified as a key driver of mucosal inflammation and barrier dysfunction **[Deng et al., 2023]**. UC-enriched bacteria carrying proteins that mimic UPR machinery may exploit or amplify this pre-existing ER stress, potentially contributing to the sustained mucosal inflammation characteristic of ulcerative colitis. The depletion of translocation of peptides or proteins into host (log₂fc = −1.67) specifically in UC — absent in CD — suggests that UC bacteria rely less on direct host cell translocation and more on surface and secreted protein interactions, a mechanistic distinction with implications for the design of microbiome-targeted therapeutic strategies.

Shared between both IBD subtypes are GTPase activator activity (log₂fc +9.33 CD, +8.66 UC) and protein targeting to vacuole (log₂fc +8.33 CD, +8.66 UC). The pan-IBD enrichment of GTPase activator mimicry is notable: Rab and Ras family GTPases regulate vesicle trafficking, bacterial autophagy, and innate immune signalling, and bacteria that carry GTPase activator-like proteins may broadly interfere with host vesicular immunity irrespective of disease subtype **[Personnic et al., 2016].**

#### 4.10.6 The SRNM axis as a unifying framework

Taken together, the neuronal, signalling, and SRP findings delineate a coherent systems-level axis operating in the healthy and IBD gut **(Figure 11a)**. In health, the microbiome maintains dual mimicry of host neuronal projection and differentiation machinery on one hand, and SRP-mediated membrane protein targeting on the other. These two mimicry systems are not independent: SRP function directs the localisation of bacterial membrane proteins that interact with host neuronal and immune cells, and the loss of both systems simultaneously in IBD suggests coordinated functional remodelling rather than independent pathway disruption. We propose that the SRNM axis represents an evolved property of the healthy gut microbiome — a molecular interface through which commensal bacteria contribute to the maintenance of neuroimmune homeostasis through structural mimicry of host regulatory proteins.

**Figure 11.**
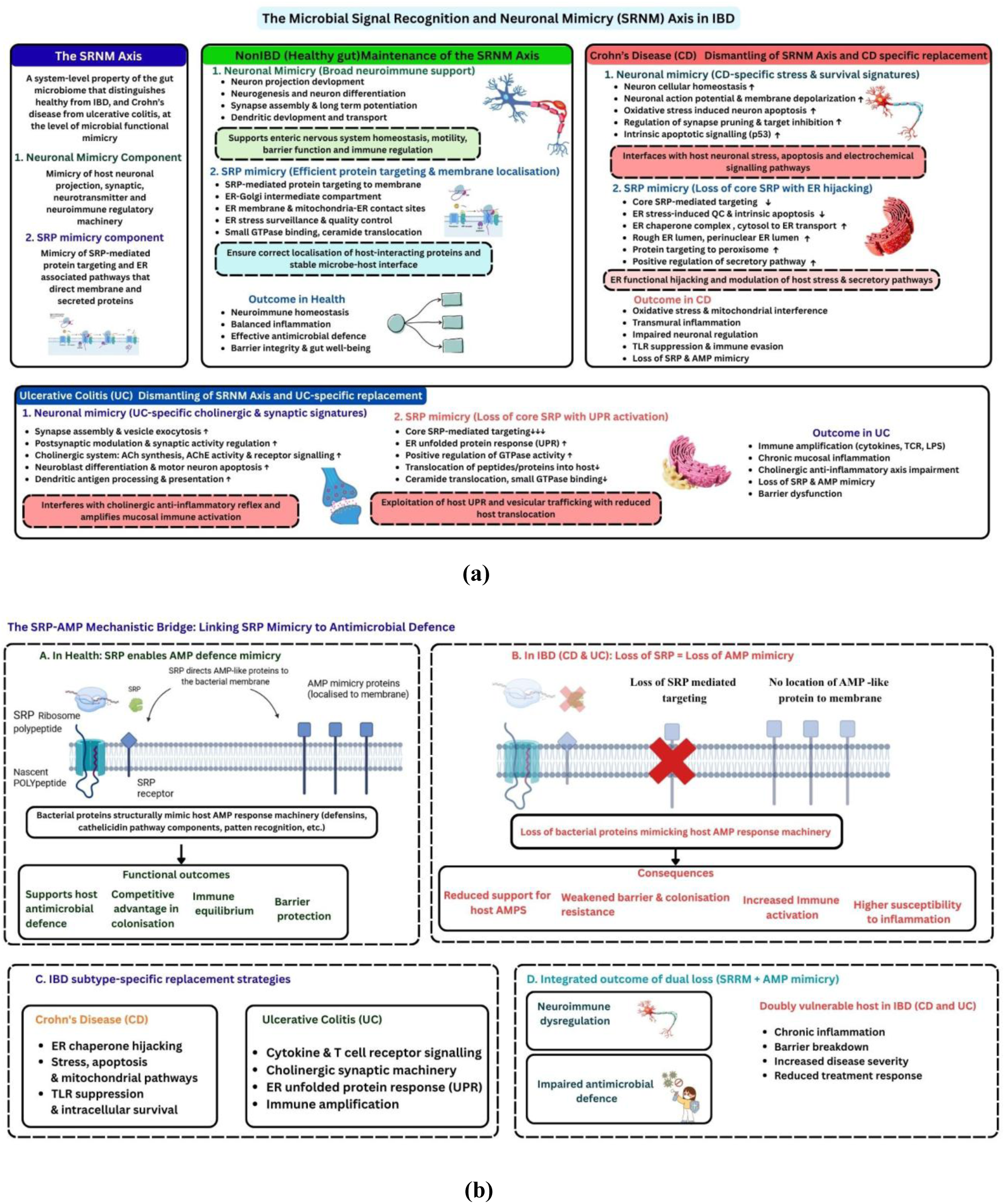
The Microbial Signal Recognition and Neuronal Mimicry (SRNM) axis in IBD and its mechanistic bridge to antimicrobial peptide defence. (a) SRNM axis in healthy and its coordinated loss in IBD with disease subtype-specific replacement in UC and CD. (b) The SRP-AMP mechanistic bridge presenting the role of SRP mimicry in AMP defence

In IBD, this axis is dismantled and replaced by disease-subtype-specific signatures that reflect the distinct inflammatory environments of CD and UC. CD bacteria acquire mimicry of neuronal stress, apoptotic signalling, action potential regulation, ER chaperone function, and TLR suppression — a functional repertoire consistent with adaptation to the oxidative, transmural inflammatory environment of Crohn’s disease. UC bacteria acquire mimicry of cholinergic synaptic machinery, cytokine and T cell receptor signalling, and the unfolded protein response — a repertoire consistent with exploitation of the mucosal immune amplification and ER stress landscape of ulcerative colitis. The divergence between these two disease-specific mimicry signatures is as great as the divergence betwe

en either disease and health, providing a molecular-level explanation for the clinical and pathophysiological distinction between CD and UC that has long been recognised but incompletely understood at the mechanistic level.

These findings open several avenues for future investigation. First, the bacterial taxa contributing to SRNM axis components should be identified through taxonomically restricted analyses, to determine whether specific commensal or pathobiont species are the primary carriers of health-associated or disease-specific mimicry proteins. Second, the functional consequences of SRNM axis mimicry proteins should be tested in cell-based systems, particularly with respect to the CD-specific suppression of TLR signalling and the UC-specific modulation of the cholinergic anti-inflammatory reflex. Third, the SRNM axis provides a rational basis for microbiome-targeted therapeutic strategies aimed not merely at restoring microbial composition but at restoring specific functional mimicry — particularly the neuronal projection and SRP targeting mimicry that characterises the healthy gut. Whether restoration of these mimicry functions through targeted probiotic supplementation or faecal microbiota transplantation can ameliorate the neuroimmune disturbances of IBD is a question of both scientific and clinical importance.

### 4.11 The AMP angle

What is already known is that: (i) AMPs are produced by the host epithelium (defensins, cathelicidins) to control gut microbiota **[Chairatana & Nolan, 2017];** (ii) Some gut bacteria produce their own AMP-like molecules (bacteriocins) **[Heilbronner et al., 2021],** (iii) Dysbiosis in IBD is associated with reduced AMP production by host Paneth cells **[Wehkamp & Stange, 2006],** and (iv) A few studies have shown that specific commensal bacteria can modulate host AMP expression **[Schauber et al., 2006].**

What has never been shown before is that gut bacteria in healthy individuals carry proteins that structurally mimic the host’s own antimicrobial humoral immune response mediated by AMPs — and that this mimicry is entirely absent in both Crohn’s disease and ulcerative colitis. Our findings support this.

It is not about bacteria producing AMPs or about bacteria regulating host AMP gene expression. It is about bacteria carrying proteins that look like the host AMP response machinery at the sequence level — molecular mimicry of the defence pathway itself. And it disappears completely in IBD — in both subtypes (CD as well as UC) simultaneously.

### 4.12 SRP-AMP mechanistic bridge

The unified narrative, thus becomes, that a healthy gut microbiome maintains two parallel mimicry systems: (i) SRP/neuronal mimicry — supporting neuroimmune homeostasis, and (ii) AMP response mimicry — participating in host antimicrobial defence (**Figure 11b**). In IBD, both are dismantled simultaneously. The microbiome loses its contribution to both neuroimmune regulation AND antimicrobial defence. The host is left doubly vulnerable — neurologically dysregulated and immunologically under-supported. In CD and UC, the replacement mimicry diverges — CD toward stress/apoptosis/mitochondrial interference, UC toward cytokine/immune amplification — but both subtypes share the same fundamental loss.

Overall, we can say that the healthy gut microbiome maintains a dual functional mimicry axis — neuroimmune (SRNM) and antimicrobial — that is coordinately dismantled in IBD. One mechanistic bridge between them could be SRP mimicry — because SRP directs AMP-resistance proteins to the bacterial membrane. The loss of SRP mimicry and loss of AMP response mimicry in IBD may not be independent. They could reflect the same underlying shift in bacterial membrane biology.

### 4.13 Limitations and Future Directions

The GO term frequency analysis captures the functional representation of microbial proteins inferred from SwissProt homologs, but does not directly measure the expression or activity of these proteins in the gut environment. Some high-frequency GO terms may reflect proteins present in multiple taxa but expressed at low levels, while rare but highly expressed proteins may contribute functional mimicry signals not captured here. Integration with metaproteomics or metatranscriptomics data would provide a more direct measure of functionally active mimicry.

The human protein mimicry candidates identified here — particularly the lysosomal carbohydrate-processing enzymes most frequently matched across groups — represent a prioritized set for experimental follow-up. Peptide-specific T cell assays and autoantibody profiling in IBD patient cohorts could determine whether cross-reactive immune responses to these microbial-human protein pairs exist and correlate with disease characteristics. Structural mimicry analysis using AlphaFold2-predicted structures and Foldseek-based alignment could complement the sequence-level approach and identify structural mimicry not detectable by sequence comparison. Integration with the HMP2 metatranscriptomics and metaproteomics datasets would allow validation of whether the GO term-inferred functional signals correspond to actually expressed and translated microbial proteins.

## 5. Conclusion

We present the first metagenome-wide analysis of molecular mimicry across IBD subtypes, showing that the healthy gut microbiome maintains broad structural resemblance to host neuronal, antimicrobial, and immune-regulatory machinery, a repertoire that is coordinately lost in disease rather than simply diminished. CD and UC do not merely diverge from health; they diverge from each other, replacing this lost mimicry with distinct, pathophysiologically coherent signatures, CD toward stress and survival pathways, UC toward immune amplification. It was observed that phosphorylation-related binding functions are convergently depleted in both IBD subtypes relative to healthy controls. Interestingly, neuronal, SRP-mediated, and antimicrobial peptide mimicry are dismantled together, defining a coordinated Microbial Signal Recognition and Neuronal Mimicry (SRNM) axis, potentially bridged mechanistically through shared SRP-dependent membrane targeting. A candidate mimicry link to NOD2, the principal genetic risk locus for CD, further connects host genetics to microbial dysbiosis. Finally, CD and UC show distinct pathobiont profiles: CD is characterized by oral-origin bacteria including *Haemophilus parainfluenzae* and *Streptococcus thermophilus*, while UC shows specific enrichment of *Fusobacterium nucleatum* and *Prevotella* species. These findings are computational and hypothesis-generating; the next steps are functional validation and taxonomic attribution of the mimicry signals described here.

## Contributions

Ananya Anurag Anand: Conceptualization; Methodology; Writing-Original Draft, Final Draft, Review and Editing

Payal Mishra: Methodology; Writing-Original Draft

Vejendla Sharmil Srivathsa: Methodology; Writing-Original Draft

Vidushi Yadav: Methodology

Sintu Kumar Samanta: Conceptualization; Supervision; Investigation; Writing-Review and Editing.

## Acknowledgements

Ananya Anurag Anand, Payal Mishra and Vejendla Sharmil Srivathsa are thankful to MoE, Govt. of India for their fellowship. Vidushi Yadav is thankful to UP-CST (Council of Science and Technology, U.P.) for her fellowship. All the authors are thankful to IIIT Allahabad for providing research facility. It is to note that Generative AI (Claude Sonnet 4.6) was used for improving the language wherever necessary.

## Statements and Declarations

### Ethics approval

There were no human subjects or animal experiments in our study.

