## Supplementary File 5 for "Molecular Mimicry in Inflammatory Bowel Disease: Multi-layered Functional and Sequence-level Analysis of Gut Microbial Proteins Mimicking the Human Proteome"

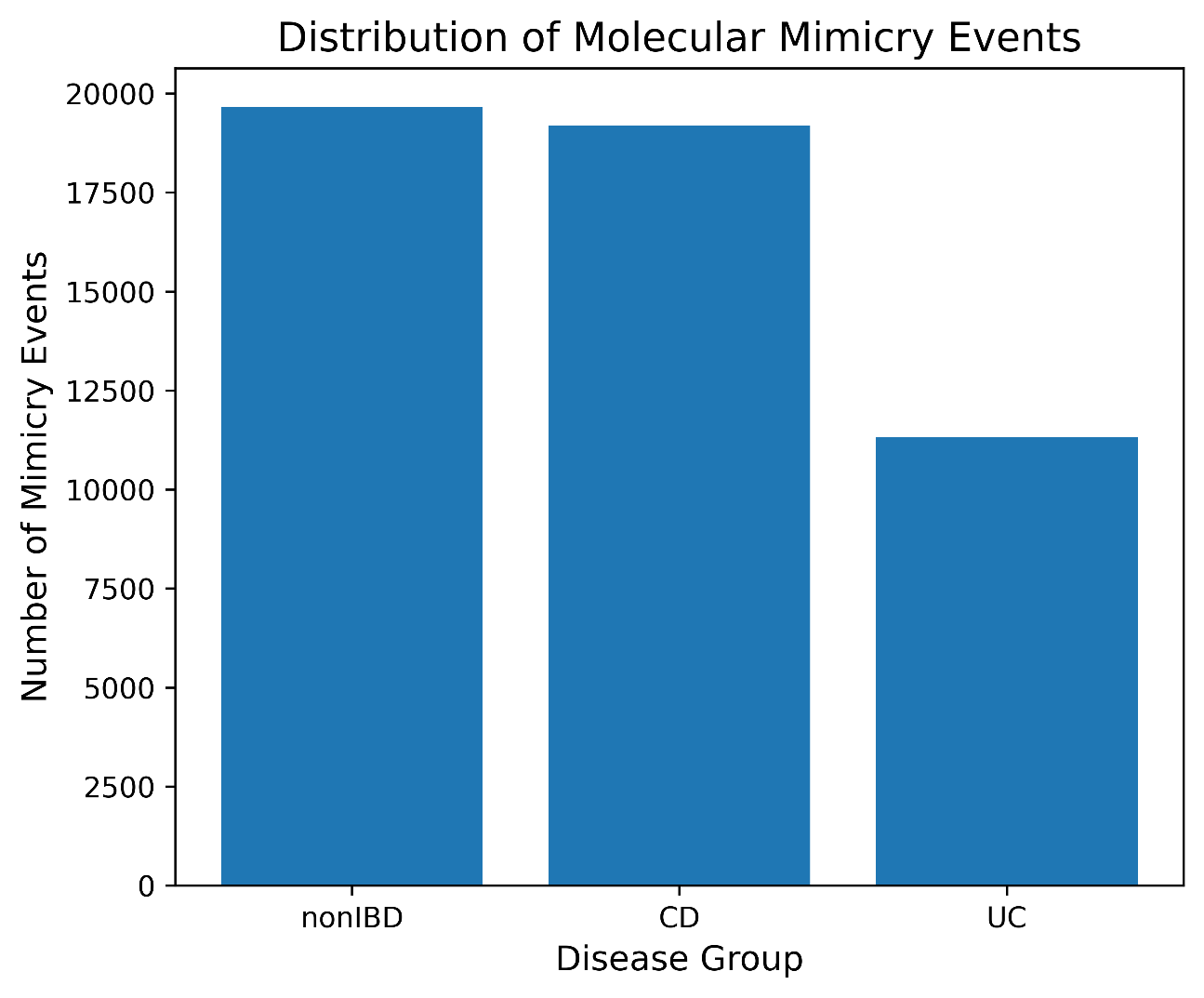

**Figure S1. The distribution of molecular mimicry events**

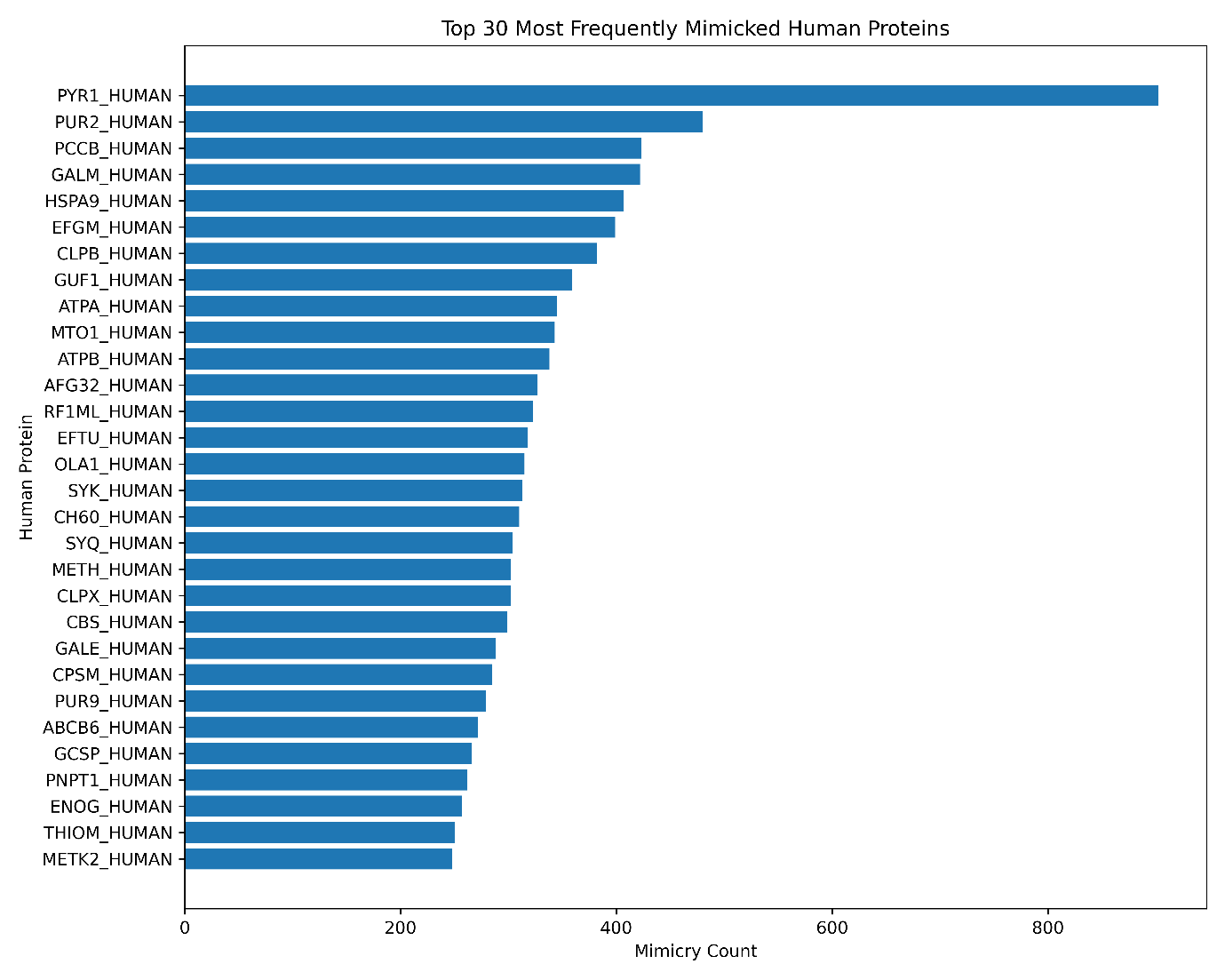

**Figure S2. Top 30 most frequently mimicked human proteins**

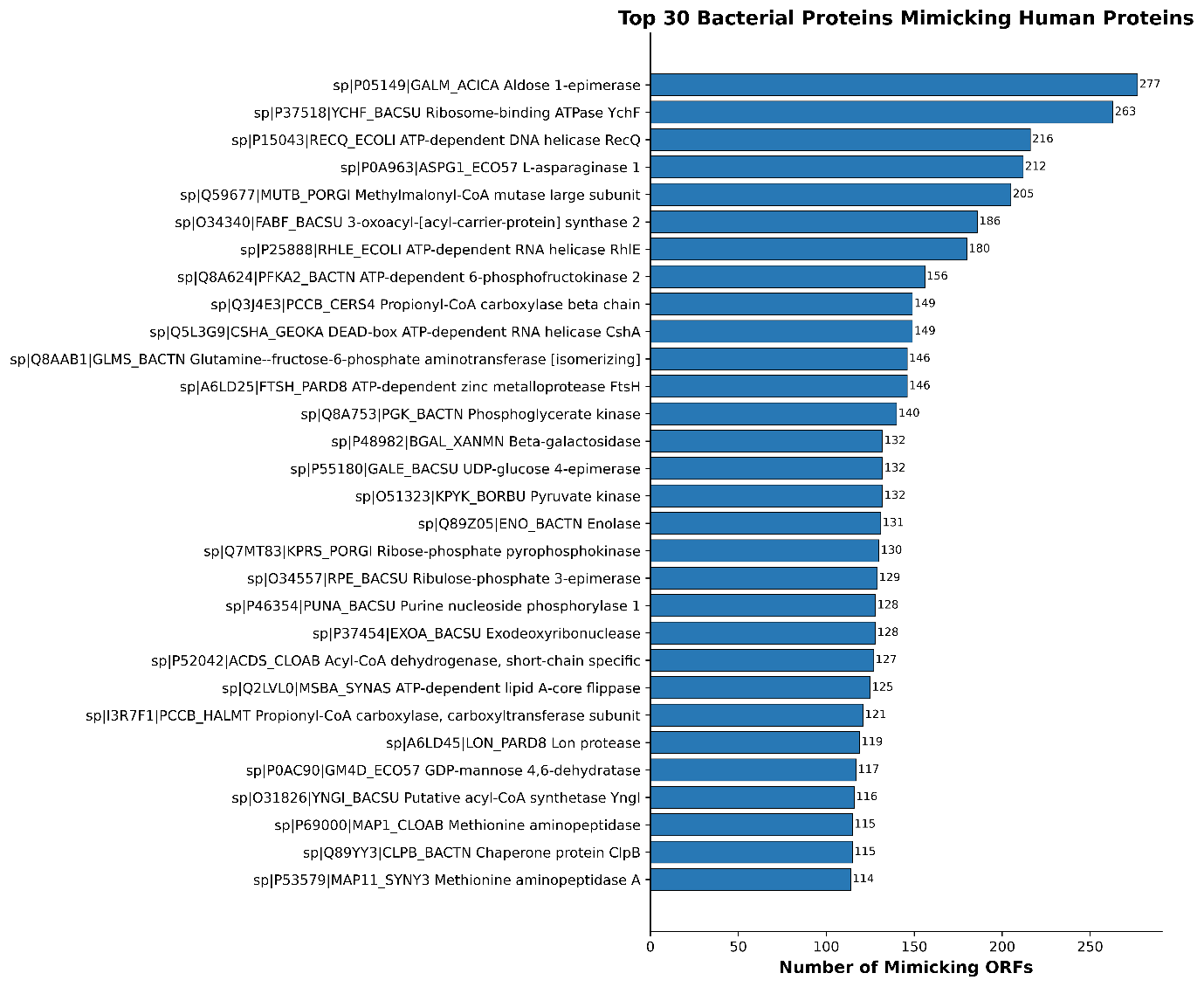

**Figure S3. Top 30 bacterial proteins mimicking the human proteins**

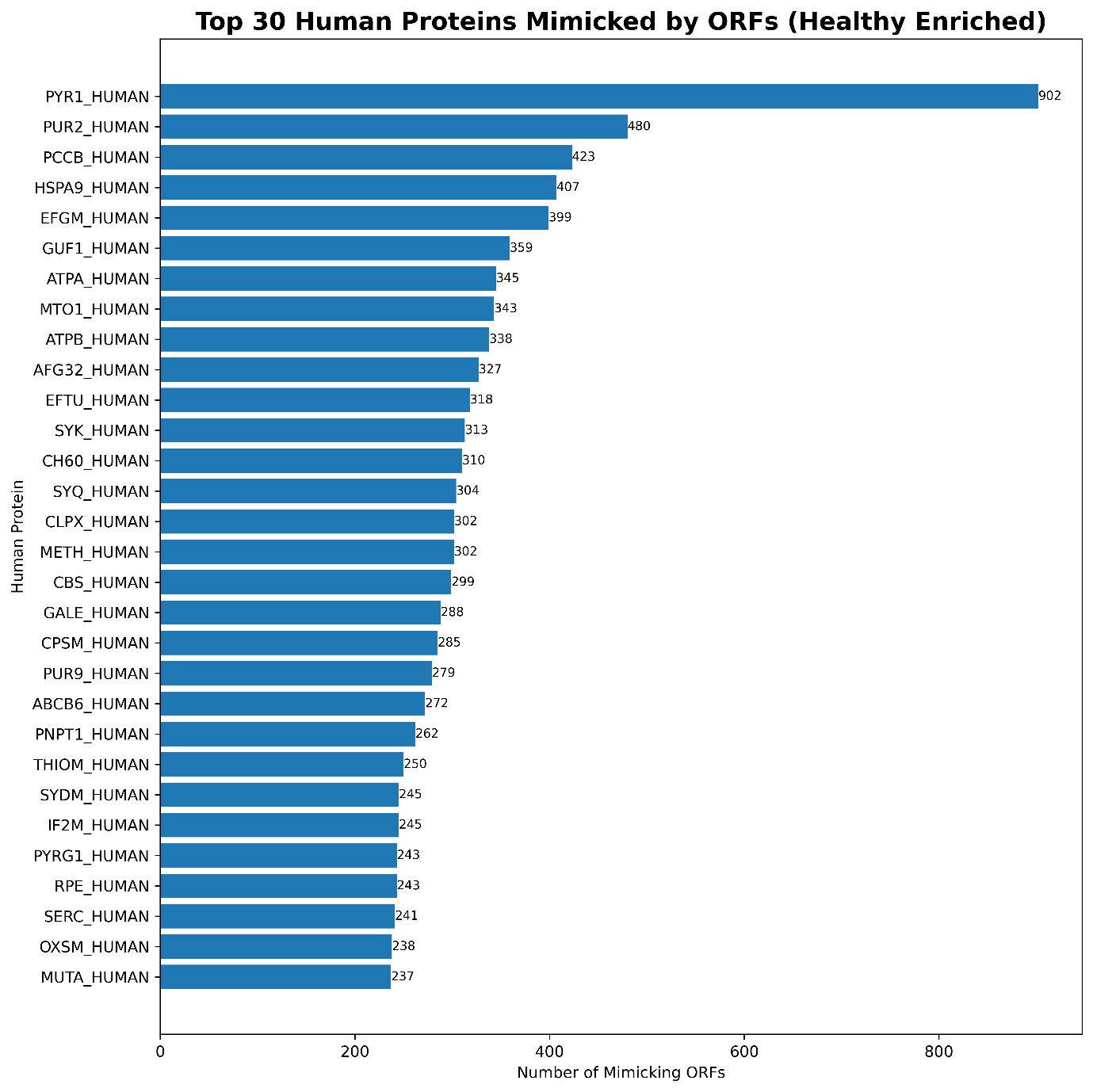

**Figure S4. Top sequence-validated human mimicry targets in the Healthy cohort**

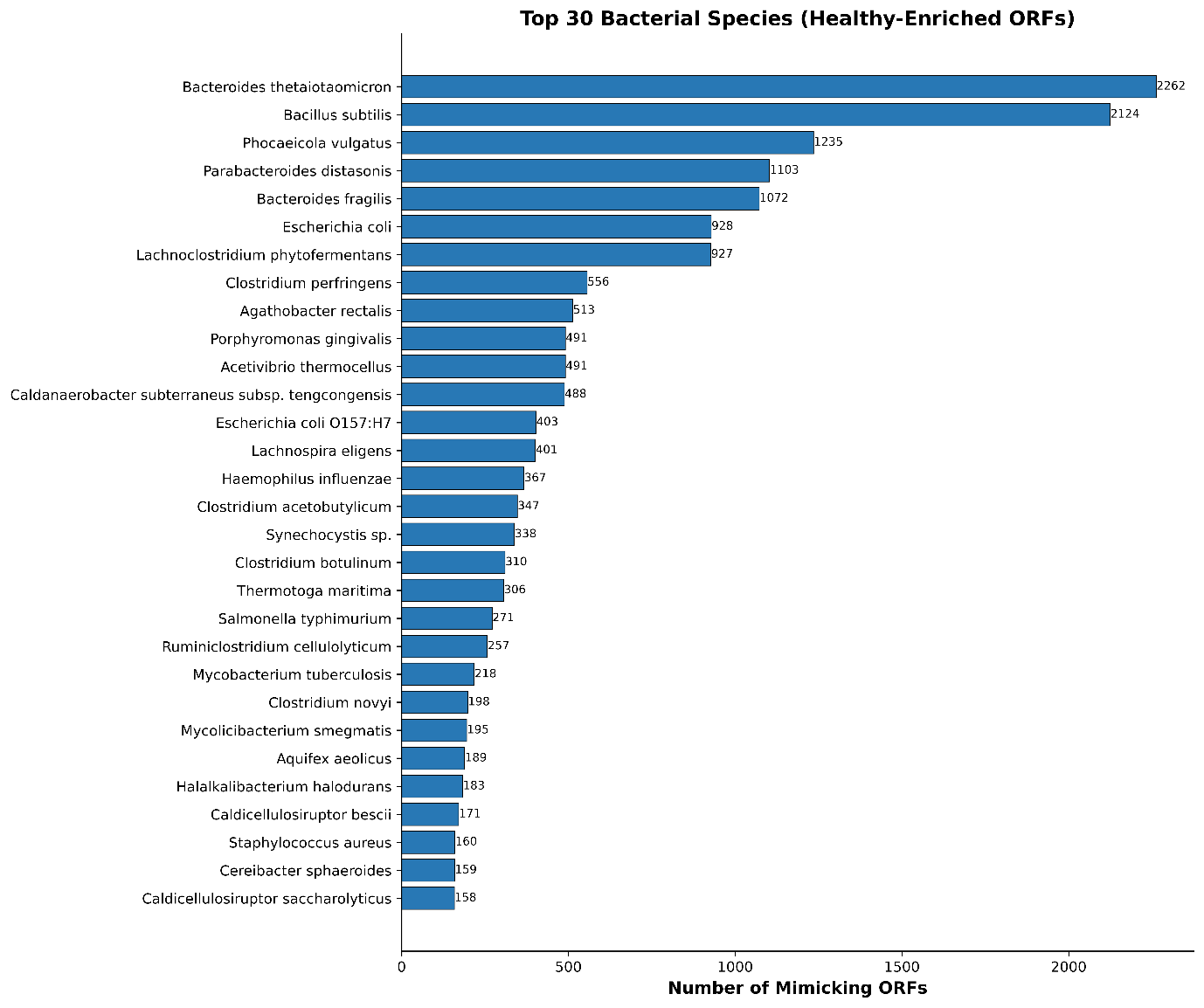

**Figure S5. Taxon-resolved attribution of the sequences from healthy cohort**

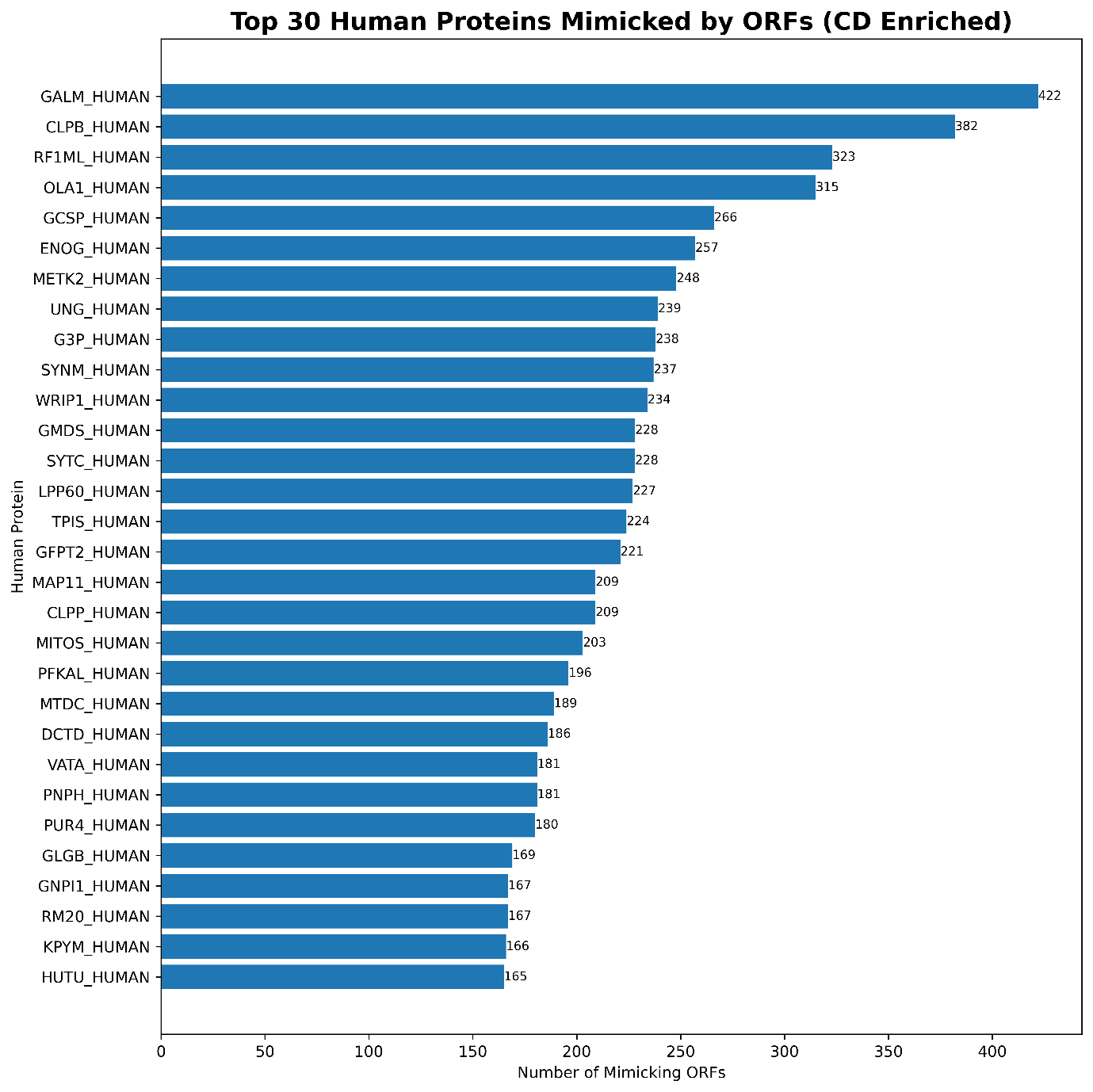

**Figure S6. Top sequence-validated human mimicry targets in the CD cohort**

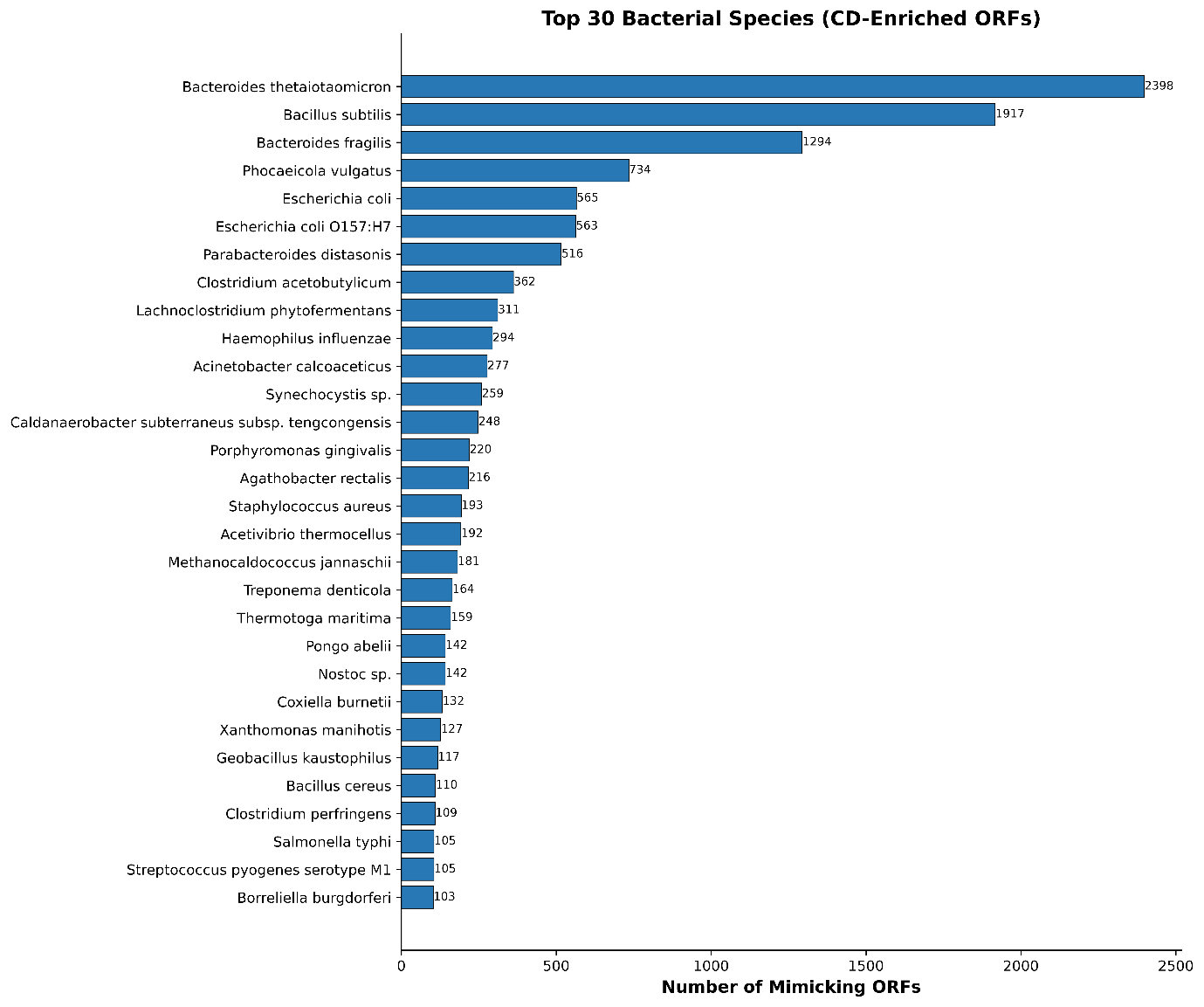

**Figure S7. Taxon-resolved attribution of the sequences from CD cohort**

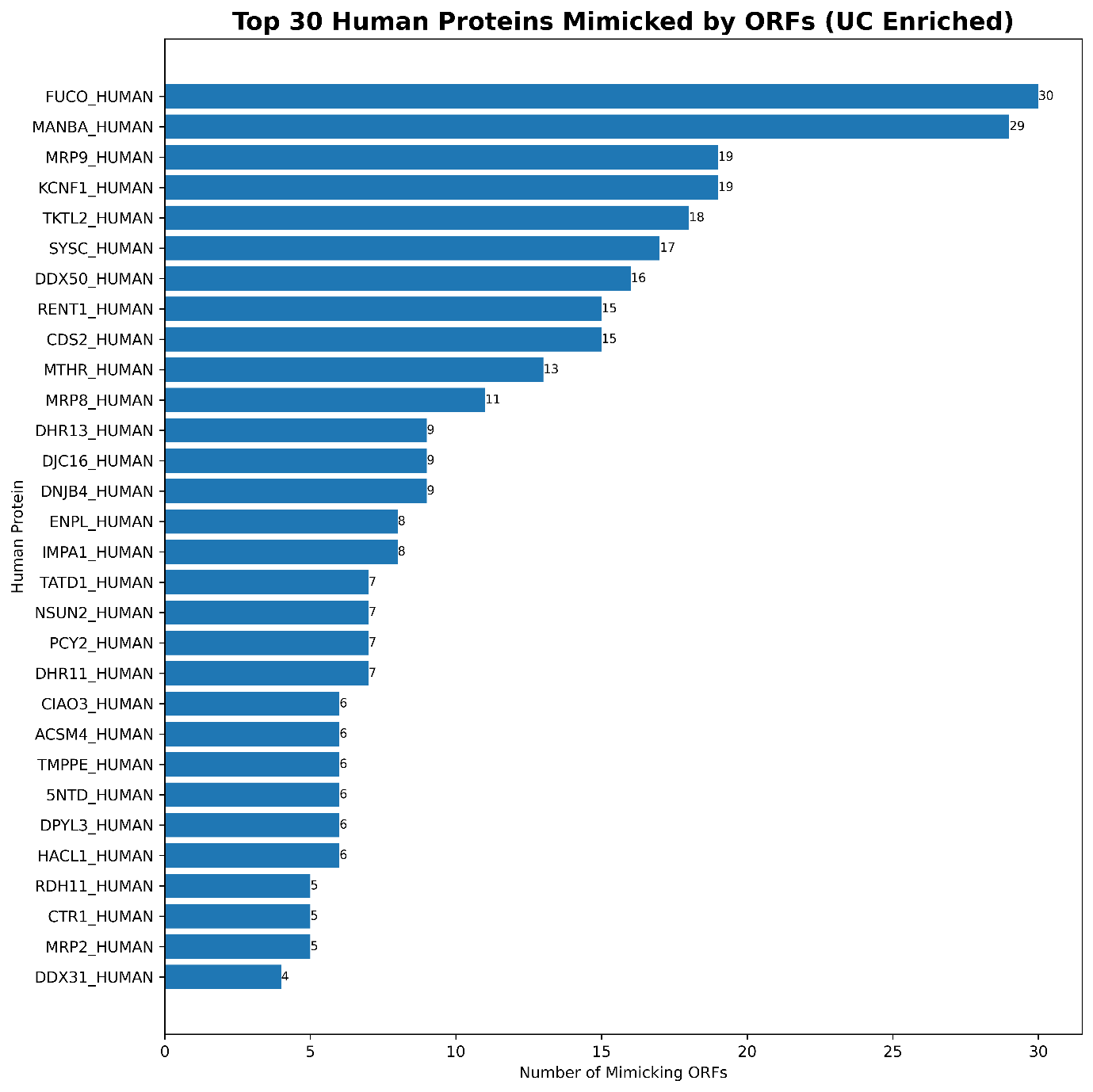

**Figure S8. Top sequence-validated human mimicry targets in the UC cohort**

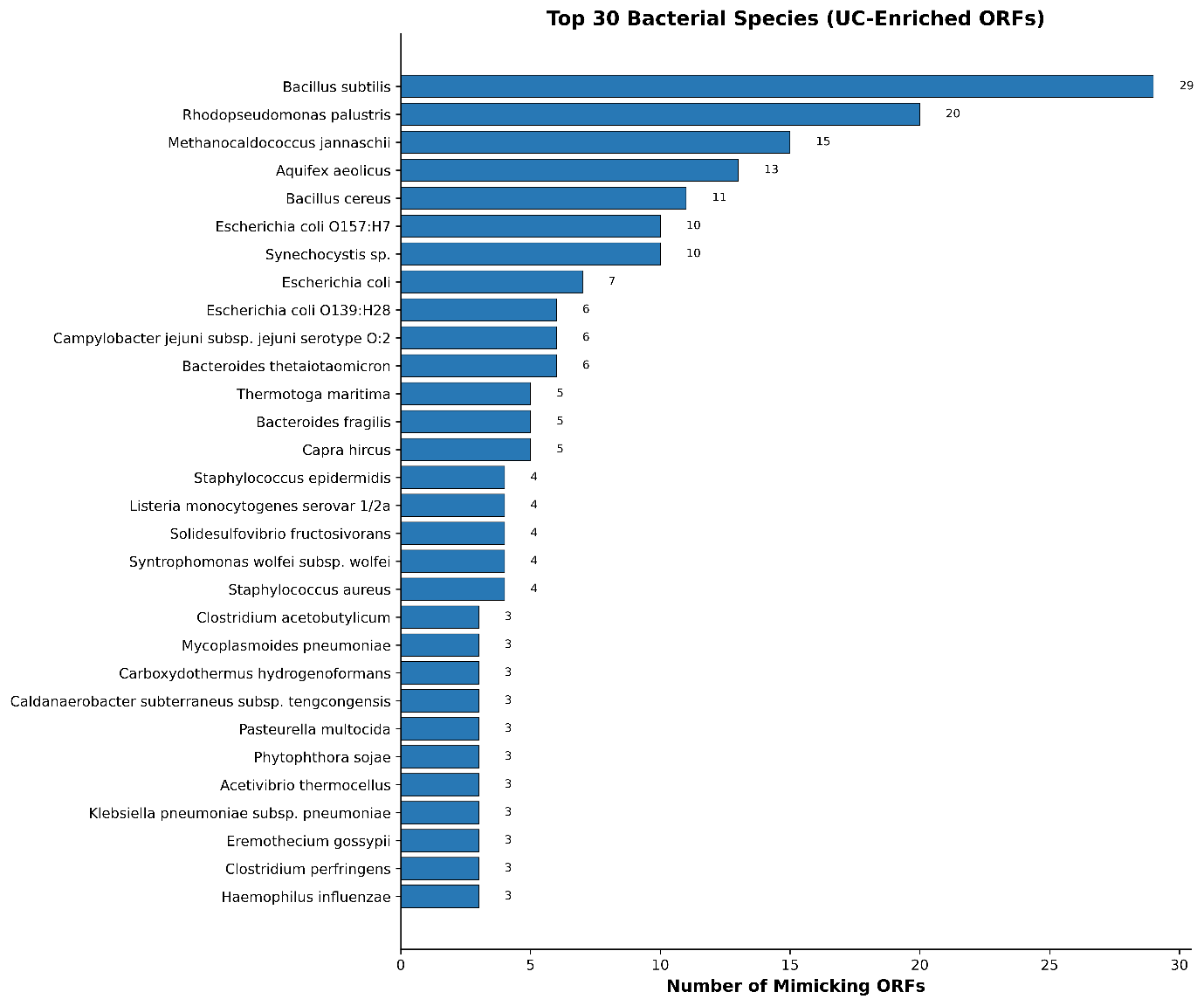

**Figure S9. Taxon-resolved attribution of the sequences from CD cohort**

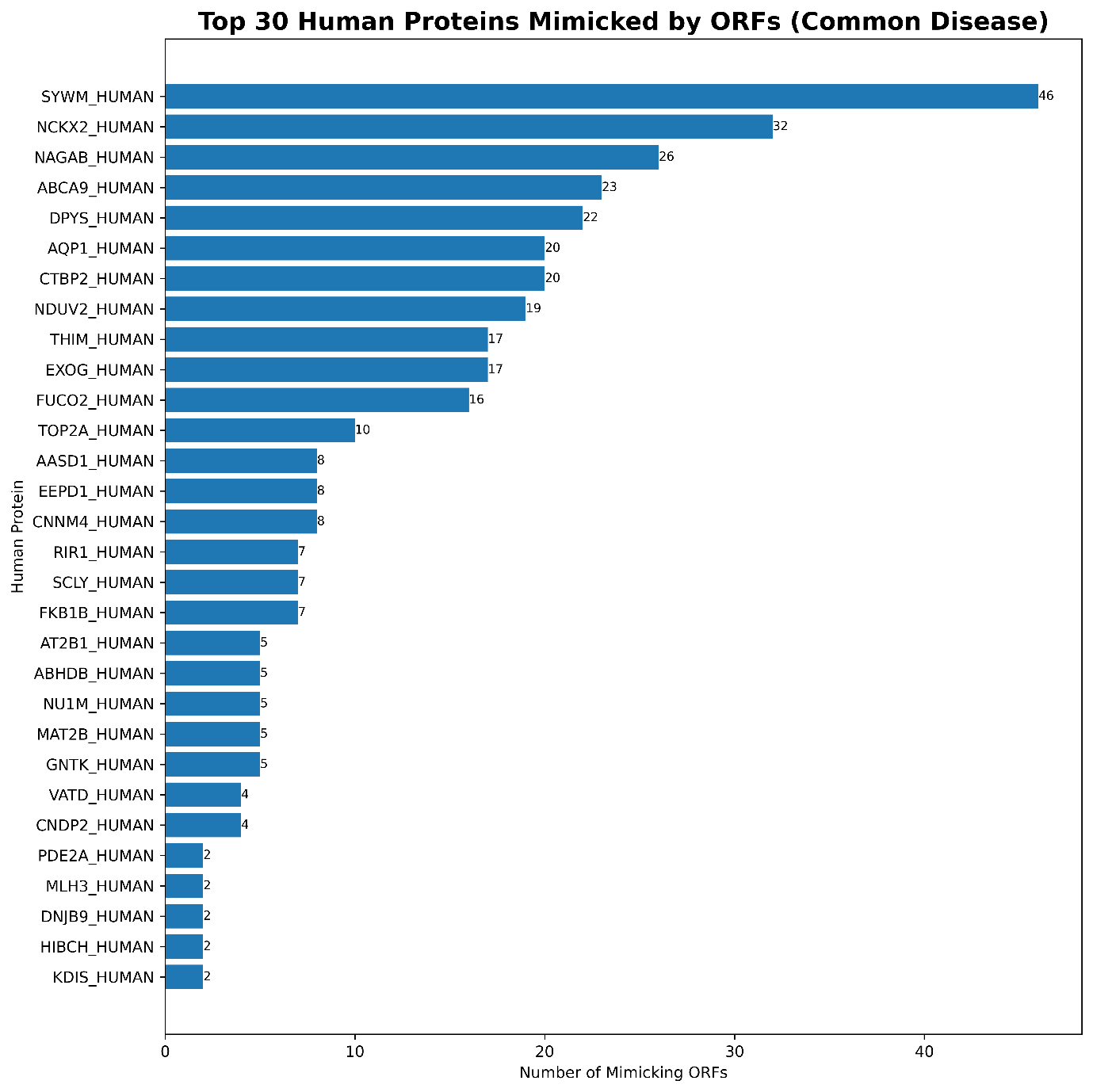

**Figure S10. Top 30 human proteins commonly mimicked by ORFs from both CD and UC gut.**
